# Single-Cell Multimodal Profiling Reveals Maternal Diabetes-*Nkx2-5* Interaction in Congenital Heart Disease in a Genetic Model Recapitulating Human Embryopathies

**DOI:** 10.64898/2026.10.07.757272

**Authors:** Stephanie Ibrahim, Julien Maurizio, Nikita Ved, Thibault Brunet, Camille Humbert, Damien Conrozier, Sofian Ameur, Gwenaelle Collod, Dorota Szumska, Jack J. Miller, Frances M. Ashcroft, Bernard Thienpont, Anne-Karin Kahlert, Duncan B. Sparrow, Sonia Stefanovic

**Author notes:** These authors contributed equally to this work (Stephanie Ibrahim, Julien Maurizio). The MR Research Centre, Department of Clinical Medicine, Aarhus University, Aarhus, Denmark and Clarendon Laboratory, Department of Physics, University of Oxford, UK.

## Abstract

The etiological complexity of congenital heart disease (CHD) arises from the interaction between genetic and environmental factors. We hypothesized that environmental exposure could potentiate a genetic susceptibility, leading to an increased incidence of specific cardiac defects. To test this hypothesis, we used ßV59M+ female mice exhibiting severe hyperglycemia (>28 mmol/L) without dyslipidemia, a model we characterized that faithfully recapitulates human diabetic embryopathy. These females were crossed with males haploinsufficient for *Nkx2-5*, a transcription factor essential for cardiac development and implicated in human CHD. Our results show that the combination of genetic susceptibility and a hyperglycemic maternal environment (G×E) leads to a higher frequency and severity of cardiac malformations, including ventricular septal defects specifically involving the muscular septum, univentricular morphologies, and dilation with thinning of the myocardium. These structural abnormalities were associated with transcriptional changes affecting the contractile identity of the chamber myocardium. Single-nucleus multiomic profiling (snRNA-seq and snATAC-seq) of the E9.5 cardiac region revealed that the interaction between maternal diabetes and *Nkx2-5* haploinsufficiency profoundly reshapes transcriptional and chromatin accessibility landscapes. These changes affect not only metabolic pathways, but also cell cycle progression in second heart field (SHF) progenitors, likely contributing to the observed structural cardiac defects. Multiomic integration further identifies a disrupted indirect *Nkx2-5–Aldh1a2* regulatory axis in posterior SHF progenitors, while demonstrating that maternal diabetes selectively affects a restricted subset of *Nkx2-5*-dependent regulatory elements associated with cardiomyocyte maturation. Our findings establish a comprehensive single-cell multiomic resource that will facilitate the integration of human GWAS and other genomic datasets, enabling the identification and functional interpretation of disease-associated loci.

## Introduction

Congenital heart disease (CHD), characterized by structural malformations of the heart, represents the most prevalent form of birth defect, affecting approximately 1,7% of live births worldwide, with even higher rates reported in miscarriages and stillbirths (GBD 2017 Congenital Heart Disease Collaborators 2020; Diller et al. 2021). Septal defects affecting the atria and/or ventricles constitute over 50% of CHD cases (Diz et al. 2021). Although substantial efforts have been dedicated to identifying genetic contributors, definitive genetic causes are identified in only a minority of cases, despite almost 200 genes being associated with CHD (Yang et al. 2022). The etiological complexity of CHD is further compounded by contributions from epigenetic, environmental, and gene-environment interactions (G×E). Even in families carrying pathogenic sequence variants in a single gene, the phenotype often shows variable penetrance and expressivity (Prendiville et al. 2014). Beyond genetic factors, many non-genetic influences have been implicated in CHD. These include: exposure to teratogenic chemicals such as pesticides and dioxins (Kopf and Walker 2009); maternal consumption of alcohol (Dorsey et al. 2025) or use of medications like isotretinoin (Khiali et al. 2018), thalidomide (Rengel et al. 2025), or anti-epileptic drugs (Petersen et al. 2017); infections such as rubella or influenza during pregnancy (Overall 1972); maternal smoking (Alhindal et al. 2025); residence at high altitudes (Linares et al. 2025); and obesity (Jacinta Isabelle Kalisch-Smith et al. 2020; McMullan et al. 2025). In the present study, we specifically focused on pregestational maternal diabetes, investigating its role both as a causal factor and as a potential modifier of CHD phenotypes. Maternal pregestational diabetes mellitus stands out as a common and significant risk factor, increasing the likelihood of CHD in offspring by approximately 3 to 5 times (Helle and Priest 2020; Persson et al. 2019; Øyen et al. 2016; Letao Chen et al. 2019). Insights from animal studies have demonstrated that pregestational diabetes mellitus can impair heart development by affecting key cellular and molecular processes, particularly within specific cardiac progenitor populations (Manivannan et al. 2022; Nishino et al. 2023). Recent findings highlight that pregestational diabetes mellitus selectively disrupts cardiac formation by altering the development of the second heart field (SHF) and neural crest derivatives, leading to impaired cardiomyocyte differentiation and axis patterning (Nishino et al. 2023). All forms of maternal hyperglycemia are associated with a similar spectrum of foetal cardiac anomalies. These include heterotaxia, conotruncal defects (Ferencz et al. 1990) (double outlet right ventricle, transposition of the great arteries, persistent *truncus arteriosus*), single ventricle, septal defects, and foetal cardiac hypertrophy (Lisowski et al. 2010a; Depla et al. 2021). However, the wide spectrum of CHD observed in the offspring of mothers with pregestational diabetes may not be fully explained by maternal hyperglycemia alone. The phenotypic variability observe suggests that genetic factors may shape the foetal response to the diabetic intrauterine environment, supporting a G×E model of CHD pathogenesis. However, the molecular and cellular mechanisms through which genetic susceptibility modulates environmental effects remain poorly understood, highlighting the need for advanced models that capture this variability and can be used to dissect its biological underpinnings.

The analysis of complex traits in pediatric cardiology poses a significant challenge. Genome-wide association studies (GWAS) typically require cohorts of tens of thousands of individuals to achieve robust statistical significance. However, population genetic heterogeneity, environmental interactions, and various confounding factors greatly limit their power to detect causal loci. In the context of CHD, these difficulties are further exacerbated by the rarity of the condition, the wide clinical heterogeneity of phenotypes, and the diversity of underlying pathogenic mechanisms. In this setting, mouse models offer a valuable strategy to overcome these limitations. Experimental data generated in mice can help guide human clinical investigations, based on the premise that key genetic and molecular pathways involved in cardiac development are conserved across species.

Previously, we and others have used mouse models to show that environmental challenges can interact with genetic susceptibility to modify the incidence or severity of congenital heart defects (Chapman et al. 2020; Moreau et al. 2019a; Basu et al. 2017a). Here we have tested the hypothesis that maternal hyperglycaemia acts as a “second hit”, increasing the penetrance and severity of cardiac defects associated with *Nkx2-5* haploinsufficiency. In humans, pathogenic sequence variants in the *NKX2-5* gene are associated with 9% of familial CHD cases (Ellesøe et al. 2016). *NKX2-5* encodes a transcription factor containing an NK-2 homeodomain that is active in cardiac progenitor cells, the first heart field (FHF) and the anterior part of the second heart field (aSHF), playing a role in the formation of various cardiac regions (Kelly 2024; Harvey 1996). In mouse models, global deletion of *Nkx2-5* leads to death around embryonic day (E) 9.5. This is accompanied by cardiac developmental arrest characterized by abnormal heart looping, reduced incorporation of anterior SHF progenitors into the primary heart tube, and failure of chamber formation, underscoring its vital role in heart morphogenesis (O. W. J. Prall et al. 2007). Additionally, *Nkx2-5* is crucial for the development, maturation, and maintenance of the cardiac conduction system, as well as for effective myocardial contraction (Briggs et al. 2008; Biben et al. 2000). *Nkx2-5* haploinsufficiency has also been associated with ventricular non-compaction (Choquet et al. 2018b; Morlanes-Gracia et al. 2021), highlighting its crucial role in myocardial maturation and compaction during cardiac development. To date, over 100 mutations in the *NKX2-5* gene have been reported in patients with CHD (Kalayinia et al. 2019). These present with diverse clinical manifestations, including atrial septal defects (ASDs), ventricular septal defects (VSDs), tetralogy of Fallot (TOF), patent *ductus arteriosus* (PDA), hypoplastic left heart syndrome, dilated cardiomyopathy, as well as cardiac conduction system defects, which are frequently associated with an increased risk of sudden cardiac death (Ellesøe et al. 2016; Reamon-Buettner and Borlak 2010; Benson et al. 1999). This phenotypic diversity is frequently observed even among members of the same family carrying the same *NKX2-5* sequence variant (Reamon-Buettner and Borlak 2010; Benson et al. 1999).

SHF cardiac progenitor cells, originating from the splanchnic mesoderm, play a crucial role in the formation of several cardiac structures (Kelly et al. 2014). Anterior SHF cells are progressively incorporated into the arterial pole of the linear heart tube, giving rise to the OFT, right ventricular myocardium, right bundle branch, and most of the right Purkinje fibre network (Liang et al. 2013; Watanabe et al. 2012a). Cells from the posterior SHF migrate into the venous pole, contributing to atrial structures, including the atrial septum (Stefanovic et al. 2020). The addition of SHF-derived cells establishes the structural basis for subsequent cardiac septation, and is essential for proper cardiac looping and coordinated OFT development alongside neural crest cells. Therefore, any disruption in SHF deployment, be it genetic or environmental, can result in a variety of CHD, especially those affecting the conotruncal region, ventricular septum, semilunar valves, and great arteries (Kelly 2024; Guijarro and Kelly 2024; Lin et al. 2012).

The impact of perturbed glucose metabolism on cardiac progenitor cell fate remains unresolved. We demonstrate that maternal hyperglycemia exacerbates both the severity and prevalence of cardiac defects in genetically-predisposed embryos. In particular, we show that the interaction between maternal hyperglycemia and embryonic *Nkx2-5* haploinsufficiency leads to an increased incidence of muscular ventricular septal defects (muscVSDs), but not membranous VSDs (mVSDs), as well as a univentricular-like morphology, myocardial wall thinning, and atrial distension. These structural anomalies are associated with a switch in contractile identity in the chamber myocardium and impaired retinoic acid (RA) signaling in cardiac progenitor cells, ultimately disrupting SHF patterning. Multiomic integration identifies a disrupted indirect Nkx2-5–*Aldh1a2* regulatory axis in posterior SHF progenitors, and highlights a selective alteration of a restricted subset of Nkx2-5-dependent regulatory elements associated with cardiomyocyte maturation. Together, these findings provide novel mechanistic insights into the variable penetrance and expressivity of monogenic CHD, and highlight maternal hyperglycemia as a potential contributor to loss of foetuses.

## Methods

### Mice

All animal procedures were either compliant with the UK Animals (Scientific Procedures) Act 1986 and approved by the University of Oxford animal welfare review board and the Home Office (project licenses PPL3003411 and PP7967396), or conducted in France in accordance with protocols approved by the nationally appointed ethics committee for animal experimentation (Ministère de l’Éducation Nationale, de l’Enseignement Supérieur et de la Recherche; Authorization No. APAFIS #33927-2021111715507212). Hemizygous mice expressing an inducible *Kir6.2-V59M* transgene (*Gt(ROSA)26Sor^tm1(Kcnj11*ßV59M)Fmas^*) specifically in pancreatic β-cells were generated using a Cre-lox system driven by the tamoxifen-inducible rat insulin promoter II (*Tg(Ins2-cre/ERT)1Dam/*J, also known as RIPII*-Cre-ER*; Dor et al. 2004), as previously described (Brereton et al. 2014, Girard et al. 2009). These are referred to as βV59M+Cre or βV59M-Cre hereafter. Tamoxifen (Sigma) was administered as a single subcutaneous injection (50-200 mg/kg body weight) in 6- to 14-week-old female mice. The tamoxifen was prepared at 10 mg/mL in corn oil. The *Nkx2-5^GFP/+^* (B6.129S1(C)-*Nkx2-5^tm4Rph^*/J) allele has been previously described (Biben et al. 2000). All experiments were conducted on mice with a C57BL/6J genetic background. Wild-type C57BL/6J mice were obtained from Charles River Laboratories. All mice were maintained under standard conditions on a 12-h light/dark cycle with *ad libitum* access to food and water. 60-day slow-release glibenclamide pellets (25 mg kg^−1^ per day ; Innovative Research of America) were implanted subcutaneously in the scruff of E3.5 pregnant females under anaesthesia as described previously (Brereton et al. 2014).

### Blood glucose measurement

Blood glucose levels were measured from tail vein blood using a glucometer (AccuCheck, Roche Diagnostics). Measurements were performed prior to injection, and at 1-2 days post-injection, as well as throughout pregnancy, including on the day of the dissection, to monitor glycaemic status. Body weight and blood glucose levels were monitored regularly. Blood glucose levels were measured in non-fasted mice in the morning, at a fixed time for all animals. After gentle warming of the tail to enhance blood circulation, a capillary blood sample (∼10 µL) was obtained by a superficial incision of the tail vein and immediately applied to a test strip (Accu-Chek). Mice with blood glucose levels ≥15 mmol/L were considered diabetic. For the G×E study, analyses were restricted to animals with hyperglycemia ≥20 mmol/L, which were used for timed matings and embryo collections.

### Collection of embryos and foetuses

For the validation of the diabetic model in studies of embryos and foetal development, C57BL/6J males and tamoxifen-treated βV59M+ or βV59M-females were used. For the G×E study, timed matings were performed between *Nkx2-5^GFP/+^* males and either tamoxifen-injected diabetic βV59M females or control females (wild-type, *RIPII-Cre-ER*, and floxed *Kir6.2-V59M* littermates, with or without tamoxifen treatment). Mating pairs were maintained on a 12-h light/dark cycle. The day of vaginal plug detection was designated as E0.5. Pregnant females were euthanized by cervical dislocation.

### Heart morphology determination

Whole embryos or isolated hearts were fixed in 4% paraformaldehyde (PFA) for over 24 hours, followed by thorough washing in phosphate-buffered saline (PBS). The morphology of whole embryos was determined by micromagnetic resonance imaging (µMRI) as previously described (Jacinta I. Kalisch-Smith et al. 2021). Fixed hearts were either paraffin-embedded, sectioned and stained with haemotoxlin and eosin, as previously described (Jacinta I. Kalisch-Smith et al. 2021), or for HREM analysis, dehydrated in a stepped series of methanol concentrations and embedded in methacrylate resin (JB4, Polysciences) containing Eosin and Acridine Orange as contrast agents, as previously described (Le Garrec et al. 2017). Single-channel images of the surface of the resin block were acquired using the optical high-resolution episcopic microscope (Indigo Scientific) and a 1× Apo objective repeatedly after removal of on average 1.7 µm thick sections. The datasets comprise 1000–2000 images of 1.83–2.89 µm resolution in x and y depending on the stage. HREM was performed on E14.5-E16.5 hearts as described previously (Mohun and Weninger 2012), using the optical high-resolution episcopic microscope (Indigo Scientific). Three-dimensional reconstruction of HREM and µMRI data was carried out using Imaris, Horos 3.3.6 (https://horosproject.org) and/or Amira for Life & Biomedical Sciences version 2019.4 (Thermo Fisher Scientific) software.

### *In situ* hybridization

RNA-FISH was performed according to the protocol of the RNAscope Multiplex Fluorescent v2 Assay (Acdbio; cat. no.323110), which detects single mRNA molecules. In briefly, E9.5 embryos and E14.5 hearts were fixed for 20–30 hr in 4% paraformaldehyde and then dehydrated in methanol. Whole-mount RNA-FISH was performed as previously described (Stefanovic et al. 2020). Embryos were imaged using an AxioScan Z1 scanner microscope (Zeiss). The following probes were used: mm-Nkx2-5-C2 (cat no. 428241), mm-Slc2a1 (cat no. 458671), mm-Slc2a3-C3 (cat no. 451571), mm-Slc2a4 (cat no. 476571), mm-Nppa (cat no. 418691), mm-Nppb (cat no. 425021), mm-Aldh1a2 (cat no. 447391) and mm-Osr1-C2 (cat no. 496281).

### RNA sequencing

RNA-Sequencing and bioinformatics analysis was performed by the Genomics and Bioinformatics facility (GBiM) from the U 1251/Marseille Medical Genetics laboratory. Hearts were isolated from E14.5-15.5 foetuses dissected in ice-cold phosphate-buffered saline (PBS). Total RNA was extracted from foetal hearts of *Nkx2-5^GFP/+^* and *Nkx2-5^+/+^* foetuses, either exposed or unexposed to maternal hyperglycemia, using the NucleoSpin RNA Plus Kit (Macherey-Nagel), following the manufacturer’s protocol. For each experimental condition, RNA was isolated from three individual foetuses (n*=* 3). Before sequencing, the quality of total RNA samples was assessed by using the Agilent Tapestation (Agilent Technologies, USA): only RNAs with RNA Integrity Numbers (RIN) above 8 were deemed suitable for sequencing and used for library preparation. For each sample, a library of stranded mRNA was prepared from 300 ng of total RNA after capture of RNA species with poly-A tail poly(A), using the Kapa mRNA HyperPrep kit (Roche, Switzerland), and following the manufacturer’s instructions. The quality and profile of individual libraries have been quantified and visualized using Qubit™ and the Agilent Tapestation (Agilent Technologies, USA) respectively. Indexed libraries were pooled and sequenced (2*100 bp paired-end sequencing) on an Illumina NovaSeq60000 platform (Illumina, USA).

### RNA-seq data processing and differential gene expression (DGE) analysis

The quality of sequencing reads was assessed using fastQC v0.11.5. Raw sequencing reads were mapped to the mouse reference genome (Mus Musculus genome assembly GRCm39 (mm39) using STAR v2.7.2b (Dobin and Gingeras 2015) and bam files were indexed and sorted using Samtools v1.73. After mapping, the number of reads per feature (ensembl release-018 mus_musculus annotation) was determined using Stringtie v2.1.6 (Pertea et al. 2015). Differential gene expression analysis was performed using a Wald test from the DESeq2 package (v1.34.0)(Love et al. 2014). P-values were adjusted for multiple testing using the method described by Benjamini and Hochberg (Benjamini and Hochberg 1995). Only transcripts with an adjusted P-value (FDR, False Discovery Rate) below 0.05 were considered as significantly differentially expressed. Differentially Expressed Genes were visualized in the form of heatmaps generated using the Pheatmap R package, or volcano plots, representing the Log2FC (log2 of the expression fold change) and the adjusted p-value, prepared using the EnhancedVolcano R package.

### Single-cell Multiome

Single-cell multiome libraries combining RNA expression and chromatin accessibility (scMultiome) were prepared from approximately 7000 nuclei isolated from the E9.5 mouse embryonic heart region, with 2 replicates (n=2) for each of the 4 experimental groups, as described in (Stephanie Ibrahim et al. 2023). Nuclei were exposed to Tn5 transposase for chromatin fragmentation and tagging, then encapsulated in Gel Beads-in-Emulsion (GEMs) using the Chromium Next GEM Single Cell Multiome ATAC + Gene Expression kit (10x Genomics), following the manufacturer’s protocol. Barcoding, pre-amplification, and construction of ATAC and GEX libraries were carried out according to 10x Genomics guidelines. Library concentration was measured using the Qubit High Sensitivity assay (Thermo Fisher Scientific), and fragment size distribution was verified with a Bioanalyzer (Agilent Technologies) prior to sequencing.

### Single-cell Multiome sequencing

The resulting libraries were pooled and submitted to IntegraGen (Evry, France) for high-throughput sequencing. Sequencing was performed on an Illumina NovaSeq™ 6000 platform using paired-end reads (2 × 100 bp). A preliminary quality control (QC) was conducted on an aliquot of the library pool (1–10 million read pairs) to assess performance on the flow cell. Only libraries passing this QC step proceeded to full sequencing. The sequencing run produced data from a single pooled library comprising four indexed samples, each thoroughly validated for compatibility with Illumina TruSeq technology.

### Single-cell Multiome bioinformatic analysis

We obtained 81,618 high quality nuclei (normoglycemic *Nkx2-5^+/+^*10,386; normoglycemic *Nkx2-5^GFP/+^* 18,886; hyperglycemic *Nkx2-5^+/+^* 21,146; hyperglycemic *Nkx2-5^GFP/+^* 31,200). Raw sequencing data were processed using the Cell Ranger ARC pipeline (10x Genomics, v2.0.2, 2022), which performs demultiplexing, alignment to the mm10 reference genome, barcode assignment, and quantification of both transcriptomic (including intronic reads) and chromatin accessibility data. Summary reports generated by Cell Ranger ARC were reviewed for initial quality assessment. Downstream analyses were performed using R software (v 4.3.2) and especially Signac (v1.12) and Seurat (v5.0.0) bioconductor packages. Ambient RNA contamination was removed using SoupX (v1.6.2) (Young and Behjati 2020), and non-informative genes, including ribosomal, pseudogenes, and mitochondrial genes, were excluded from the analysis. Quality control was jointly performed on both RNA and ATAC modalities. For each replicate, lower and upper thresholds were set individually based on total RNA counts, number of detected features, number of ATAC fragments in peaks, transcription start site (TSS) enrichment scores, and nucleosomal signal distributions. Additionally, cells with fewer than 15% of reads in peaks or with more than 1% of reads mapping to ENCODE blacklist regions were excluded. After filtering, single-cell RNA-seq dataset was processed independently from single-cell ATAC-seq.

#### Single-cell RNA-Seq

Each single-cell RNA-seq replicate was processed independently using the standard Seurat pipeline: normalization, selection of the top 2,000 variable features, data scaling, dimensionality reduction via PCA, identification of nearest neighbors and clustering using the Louvain algorithm, and projection into two dimensions using UMAP. For each replicate, cell types were annotated based on specific marker genes drawn from the literature, the UCSC Cell Browser, and gene signature scoring with the UCell R package (v2.6.2) (Andreatta and Carmona 2021). The same standard Seurat pipeline used for individual replicates (normalization, scaling, PCA, clustering, and UMAP) was then applied to each sample. Cells were annotated based on the cell type labels assigned in the replicate-level analysis, using the clustering results. To assess potential batch effects, cell type consistency within clusters and sample origin biases were qualitatively evaluated. Optionally, a quantitative step was added by adjusting the clustering resolution to match the expected number of cell types and testing for the overrepresentation of multiple cell types within individual clusters. As no significant batch effect was observed, no batch correction algorithms were employed. All samples were then merged and processed again using the standard Seurat pipeline (normalization, scaling, PCA, clustering, and UMAP). Cell types were assigned based on known markers from the literature, the UCSC Cell Browser, and UCell module score calculations. Ambiguous clusters (with multiple potential identities) were further subclustered and re-analyzed using the same Seurat workflow to refine cell type annotations. In addition, the FindMarkers function from the Seurat package was used to support these assignments. These strategies were particularly helpful for distinguishing cardiomyocyte and cardiac progenitor subpopulations. To account for the high variability in cell numbers across samples, as well as the presence of a single-replicate sample, differential abundance testing was performed using the sccomp R package (Mangiola et al. 2023). Sccomp fits a hierarchical Bayesian model in which posterior probabilities are inferred using a reference beta-binomial distribution, explicitly modeling overdispersion across features. Differential expression analysis was performed using the FindMarkers function from the Seurat R package, applying the MAST statistical test. To reduce noise from lowly expressed genes, only genes detected in at least 10% of cells in either group (min.pct = 0.1) were tested for differential expression. For functional enrichment analysis, genes with an absolute difference of at least 5% in the percentage of expressing cells between condition A (pct.1) and condition B (pct.2) were considered differentially expressed. These genes were subjected to Gene Ontology (GO) enrichment analysis using the enrichGO function from the ClusterProfiler R package (v4.10.1)(Yu et al. 2012). GO terms with an adjusted p-value < 0.05 were considered significantly enriched.

#### Single-cell ATAC-Seq

From the merged samples, single-cell ATAC-seq data were first normalized using the TF-IDF (Term Frequency–Inverse Document Frequency) method. Dimensionality reduction was then performed using Singular Value Decomposition (SVD) on the peak-by-cell matrix, producing latent semantic indexing (LSI) components representing cells in a reduced-dimensional space. These were used to embed cells in a two-dimensional UMAP space. To assess the correlation between gene expression and chromatin accessibility, and thereby estimate scATAC-seq quality based on higher-confidence scRNA-seq data, cell annotations were transferred from matched scRNA-seq profiles. First, gene activity scores were computed from scATAC-seq data using the GeneActivity function in the Signac package, quantifying fragment counts within gene bodies and 2 kb upstream. These gene activity matrices were normalized, scaled, and used as input to the FindTransferAnchors function with canonical correlation analysis (CCA) reduction, using highly variable genes from the scRNA-seq dataset. Cell labels were then transferred using Seurat’s TransferData function, which performs weighted label prediction in the LSI space. Annotation transfer achieved ∼80% accuracy compared to ground truth labels, suggesting strong concordance between transcriptomic and chromatin accessibility signals. Finally, batch effect correction was applied on the LSI embeddings using Harmony (v1.2.3), leading to improved cluster homogeneity in the UMAP projection. It was further verified that marker gene-associated peaks were enriched in ATAC clusters corresponding to the expected cell types. Peak calling was conducted with MACS (v 3.0.2) (Y. Zhang et al. 2008) on selected clusters of interest (namely, annotated cardiomyocytes, aSHF, and posterior SHF (pSHF). For each selected cluster, a FeatureMatrix was constructed by counting fragments overlapping the called peaks. This matrix was normalized using TF-IDF for downstream comparisons between conditions. Although logistic regression is more common in scATAC-seq differential accessibility analysis, MAST was used here for consistency across modalities and because the data matrix was normalized and filtered accordingly. To reduce noise from lowly accessible regions, only peaks detected in at least 10% of cells in either group (min.pct = 0.1) were tested. Gene annotation was performed using Ensembl v79, defining promoter regions as ±2 kb around the transcription start site (TSS). Peaks were associated with the nearest gene promoter using the ClosestFeature function from Signac. Visualization of chromatin accessibility tracks was carried out with the CoveragePlot function (Signac).

#### Annotation of differentially accessible regions (DARs)

A set of 260 501 peaks was constituted from peak calling with a wrapper of MACS2 (Y. Zhang et al. 2008) (CallPeaks function) from Signac (Stuart et al. 2021) on all cell types, performing independent peak calling across replicates for each condition. Overlapping peaks were merged and combined with non-overlapping ones. The following steps were performed independently for the pSHF annotated cluster and the cardiomyocyte (CM) annotated cluster : Based on this set of 260 501 peaks, DARs were identified for all pairwise comparisons : GxE *vs* control, GxE *vs* Environment, GxE *vs* Gene, Gene *vs* control and Environment *vs* Control. These comparisons were conducted with FindMarkers function from Signac (v 1.12.0) using a logistic regression and a likelihood ratio test, introducing sequencing depth as a latent variable and setting a minimum percentage cutoff of 5% across the compared conditions.Peaks were annotated with submitGreatJob and getRegionGeneAssociations from rGREAT package (v 2.4.0) (Z. Gu and Hübschmann 2023). The top 15 peaks (ranked by best p-values) showing significant chromatin changes in GxE, but not in G alone compared to the control, were selected. The log2 fold change was plotted against mean accessibility. Each point represents a peak labeled with the associated genes, with coordinates defined by the specific comparison. These 15 peaks are highlighted in blue or red according to their increased or decreased status in GxE. Peak-associated genes showing significant correlation via LinkPeaks are also labeled on the MAplots. For regions with dual gene annotations, only the gene(s) expressed or of particular biological interest are displayed on the MA plots.

#### Gene Ontology enrichment of DARs

Significantly altered regions were selected using distinct criteria for each cell type: for cardiomyocytes, regions with p-value < 0.01 and log2FC < 0 were retained as significantly less accessible (closed), whereas for pSHF, all DARs with p-value < 0.05 were included. To evaluate the relevance of these regions in cardiac development, a Gene Ontology (GO) term analysis was performed using the enrichGO function from the clusterProfiler package (v4.10.1;(Yu et al. 2012)) to highlight Biological Processes involved in chromatin remodeling. For this, the 704 (CM) less accessible peaks and the 1,392 (pSHF) DARs were assigned to their nearest gene using the ClosestFeature function. The 18,829 (CM) and 16,258 (pSHF) peak-associated genes tested for differential accessibility served as the background set. P-values were adjusted using the Benjamini-Hochberg method (Benjamini & Hochberg, 1995) and GO terms with adjusted p-value < 0.05 were selected as significant. To verify the robustness of these results, a random set of 704 (CM) and 1,392 (pSHF) peak-associated genes was tested as a negative control to ensure that no GO terms were enriched by chance.

#### Identification of candidate Nkx2-5 regulatory elements

Then, peaks harboring an Nkx2-5 binding motif were selected as potential regulatory candidates, identifying 172 peaks for CM and 201 for pSHF. First, Position Frequency Matrices (PFMs) for Mus musculus were retrieved from the JASPAR2020 database (Fornes et al. 2020) using the TFBSTools package (v1.40.0). The FindMotifs function from Signac was used to identify over-represented motifs in the selected peaks against the genomic background. The 172 (CM) and 201 (pSHF) peaks containing Nkx2-5 motifs were retained for further analysis. Putative Nkx2-5 co-binding factors were subsequently searched using the same strategy within these 172/201 peaks, this time querying the entire vertebrates collection. Finally, genes with GxE-regulated expression in snRNA-seq (1,453 for CM / 326 for pSHF; relative to control, adjusted p-value < 0.05) were linked to the Nkx2-5+ peaks (172 for CM / 201 for pSHF) using the LinkPeaks function from Signac (with p-value and score cutoffs of 0.05). This method identifies peaks correlated with regulated expression within a 500 kb genomic window around the gene. 3 genes with significant links were found for CM, and 0 for pSHF.

### Genome browser visualization

The following priviously-published genome browser ChIP-seq tracks were used: NKX2-5 (He et al. 2014), GATA4 (He et al. 2014), HDAC1 (Chronis et al. 2017), and HDAC2 (Whyte et al. 2012).

## Results

### Expression patterns of glucose transporters during mouse cardiogenesis

*In utero* exposure to hyperglycemia is likely to have the greatest effect on cells with an active glucose-uptake system. In mouse embryos, this likely occurs via the glucose transporter proteins *Slc2a1* (GLUT1), *Slc2a3* (GLUT3) and *Slc2a4* (GLUT4). However, the spatiotemporal expression patterns of these genes has not been well described. Therefore, we used RNAScope FISH on wild-type mouse embryos to further investigate. At E9.5, *Slc2a1* was expressed in a range of tissues, including the cardiac progenitors (Supplementary Fig. 1A). Specifically, *Slc2a1* was present in the SHF, the ventral wall of the pharyngeal endoderm, the primitive atria and ventricle, and the outflow tract (OFT), with clear co-localization with *Nkx2-5* in the OFT and atrial/ventricular myocardium (Fig. 1A-A’, B-B’’). By contrast, at E9.5 *Slc2a3* was only detected in the ventral wall of the pharyngeal endoderm, and not in the adjacent SHF (Fig. 1C-C”). These results are consistent with previous reports, which lacked spatial resolution to define its detailed expression pattern (Joyner and Smoak 2004; Grover-McKay et al. 1999; Ma et al. 2022; Mlynarski et al. 2015; Smith and Gridley 1992). Proximity to the ventral pharyngeal endoderm is essential for maintaining SHF progenitor cell properties and for proper SHF deployment. Thus, these transporters are expressed in cells that contribute to the maintenance of SHF progenitor cell properties via ventral endoderm–derived signaling. Consistent with this, human genetic studies have associated *SLC2A3* sequence variants with syndromic CHD, with carriers exhibiting a spectrum of cardiac anomalies, including atrial septal defects (ASD), persistent *truncus arteriosus*, and multiple ventricular septal defects (VSDs) (Conte et al. 2023a; Ma et al. 2022; Mlynarski et al. 2015). From E10.5 (Fig. 1E-E”) to E12.5 (Fig. 1F-F”). In addition, *Slc2a1* and *Nkx2-5* expression largely overlapped within the compact myocardium and the developing interventricular septum. By E14.5, *Slc2a1* expression persisted, whereas *Slc2a4* emerged in the atria and the trabecular myocardium (Fig. 1G-H). While these patterns are in agreement with published single-cell RNA-seq data (de Soysa et al. 2019), our analysis further delineates expression within and surrounding the SHF (Supplementary Fig. 1A), and our spatial mapping provides the missing anatomical context. Collectively, *Slc2a1* predominates during early cardiogenesis, while *Slc2a3* is more restricted and may support arterial-pole progenitors.

**Fig. 1.**
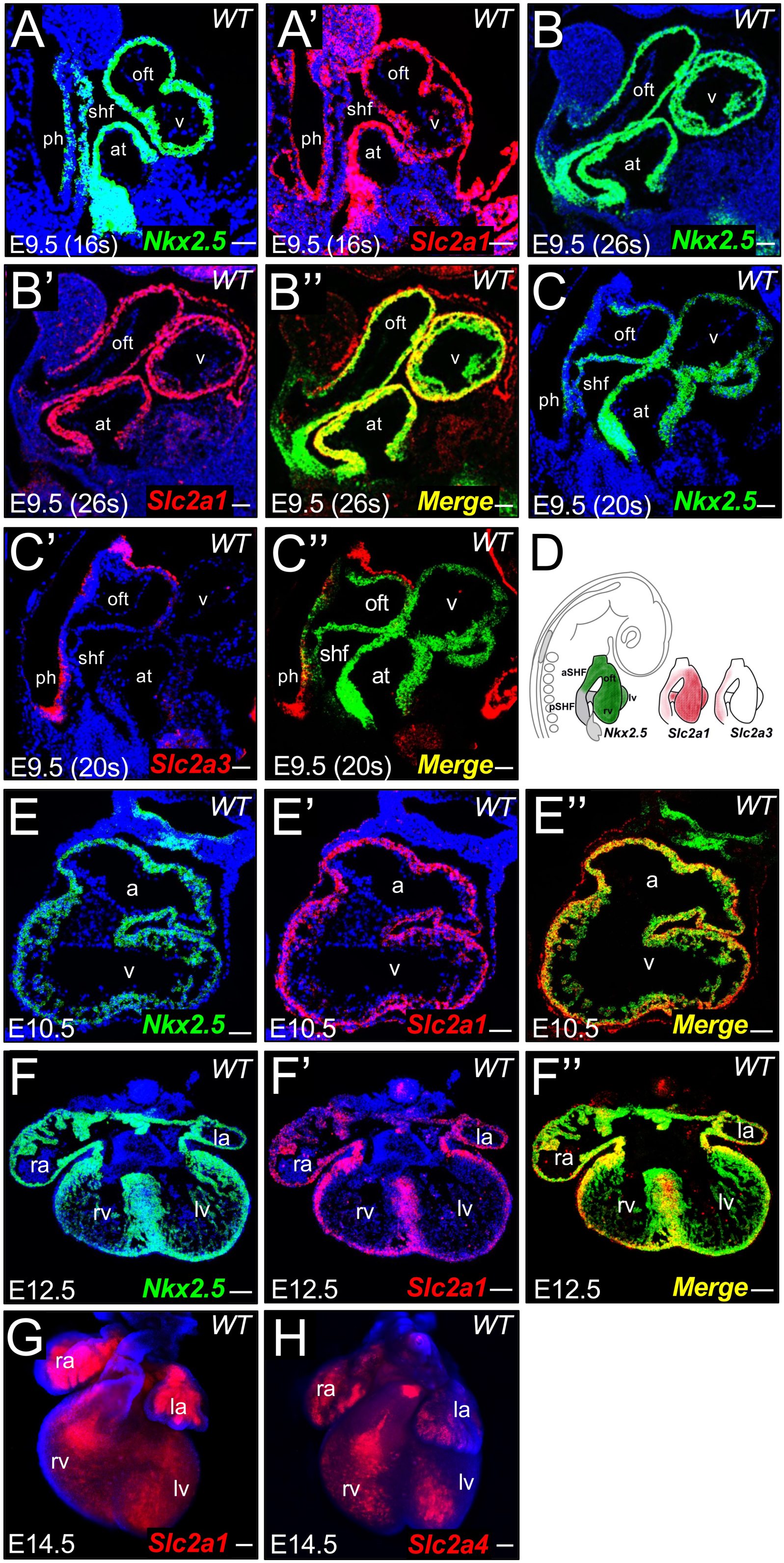
Spatial and temporal co-expression of glucose transporters and *Nkx2-5* during murine heart development. RNAScope fluorescence in situ hybridization (FISH) was performed on wild-type embryonic and foetal hearts to examine *Slc2a1 (Glut1), Slc2a3 (Glut3), Slc2a4 (Glut4) expression and* co-localization with *Nkx2-5* throughout development. (A-A’) Serial sections of an E9.5 (16-somite stage) embryonic heart show *Nkx2-5* and *Slc2a1* expression in the second heart field (shf), the ventral wall of the pharyngeal endoderm (ph), as well as the atria (at), ventricle (v), and outflow tract (oft). (B-B”) At E9.5 (26s), *Nkx2-5* and *Slc2a1* co-localize in the atria, ventricle, OFT (separate channels in B–B’, merged in B’’). (C–C’’) At E9.5 (20s), *Slc2a3* expression overlaps with *Nkx2-5* primarily in the ventral wall of the pharyngeal endoderm (ph) adjacent to the second heart field (separate channels in C–C’, merged in C’’). (D) Schematic representation of *Nkx2-5*, *Slc2a1*, and *Slc2a3* expression domains in the embryonic heart. (E–E’’) By E10.5, *Slc2a1* expression overlaps with *Nkx2-5* in the myocardial walls of both atria and ventricles (separate channels in E–E’, merged in E’’). (F–F’’) At E12.5, *Slc2a1* co-localizes with *Nkx2-5* in the right and left atria (ra, la), ventricles (rv, lv), and more prominently in the muscular interventricular septum and in the compact zone of the myocardium (separate channels in F–F’, merged in F’’). (G–H) Whole-mount staining at E14.5 shows persistent *Slc2a1* expression, while *Slc2a4* begins to be expressed in the atria and likely in the trabecular myocardium. Scale bars: 50 µm for panels A, B, C; 100 µm for panel E; 200 µm for panels F, G, H.

### The ßV59M mouse model reflects human diabetic embryopathy

Female ßV59M+ mice develop severe hyperglycemia (>28 mmol/l) without dyslipidaemia 48 hours after injection with 200 mg/kg tamoxifen (Brereton et al. 2014). Although control female C57BL/6J mice have a much higher baseline BGL than humans (∼10 mmol/l compared to 5.5 mmol/l in humans)(“Phenome.Jax.Org/Measures/43901,” n.d.), the BGL in injected ßV59M+ mice is elevated far above the clinical range. We reasoned that injection with lower doses of tamoxifen might result in a more clinically-relevant elevation of BGL. ßV59M+ female mice were injected with 50-200 mg/kg tamoxifen at 6-10 weeks of age. Blood glucose levels (BGL) were measured 24 and 48 hours post-injection. Injected ßV59M+ mice had significantly higher BGL compared to injected ßV59M-mice lacking the *Cre* transgene (Supplementary Fig. 2A). Injected ßV59M+ mice had a wide range of BGL, with higher tamoxifen doses generally leading to higher BGLs. BGL were stable over time, and the hyperglycemic mice did not lose weight (Supplementary Fig. 2B). To test the effects of maternal hyperglycemia on embryonic development, hyperglycemic female mice were timed mated with wild-type C57BL/6J males, and maternal BGL measured at plugging, E9.5 and immediately prior to embryo collection at E15.5-16.5. For ease of analysis, we divided litters into four categories based on maternal BGL at E9.5: control (<15mmol/l), moderate (15-21.4 mmol/l), high (21.5-28 mmol/l) and severe (≥28mmol/l) (Supplementary Fig. 2C). As noted for other mouse models of maternal diabetes mice (Endo 1966; Moazzen et al. 2014) with severe hyperglycemia were less likely to initiate or maintain pregnancy to post-implantation stages. Based on external morphology (Geyer et al. 2017), embryos from hyperglycemic mothers at E15.5 were smaller and developmentally delayed by approximately 24 hours. This is in line with previous studies with chemically-induced diabetic mothers (Eriksson et al. 1982; Endo 1966; Kumar et al. 2007). Therefore, to allow a more direct comparison of embryo phenoytype between hyperglycemic and control embryos, for all subsequent analyses we collected hyperglycemic embryos at E16.5, which were stage-matched to E15.5 controls by external appearance. Hyperglycemic embryos had significantly increased BGL (Supplementary Fig. 2D), with the absolute embryonic BGL correlated to maternal BGL (r2 = 0.4969, Supplementary Fig. 2E).

Macroscopic observation at dissection revealed significant embryonic lethality and abnormality, with increasing penetrance and severity as maternal BGL increased (Supplementary Fig. 2F, Supplemental Table 1). In moderate litters, 6/27 embryos were abnormal; in high litters, 19/48 embryos were abnormal and 4/48 were dead; and in severe litters, 13/33 were abnormal and 11/33 were dead; compared to 0/76 abnormal or dead for controls. The most common abnormalities were craniofacial (exencephaly, and micrognathia), eye (anopthalmia and micro-opthalmia), or lymphatic (gross sub-cutaneous oedema and blood-filled lymphatics) (Supplementary Fig. 2 I-L, Supplementary Table 1). Defect penetrance was greater in litters from mothers with higher BGL. These observations are in keeping with observations of both human diabetic embryopathy (Zhao and Reece 2013) and previous studies of different animal models of maternal hyperglycemia. We conclude that the ßV59M mouse is a useful model of diabetic embryopathy. Diabetic embryopathy is also associated with highly penetrant CHD in humans (Hoang et al. 2017; Lisowski et al. 2010b), and in streptozotocin (STZ) (Basu et al. 2017b; Wang et al. 2015) and high-fat diet (HFD) mouse models (Y. Wu et al. 2016; Schulkey et al. 2015a). We therefore examined cardiac morphology in a subset of surviving embryos (Supplementary Fig. 2G, Supplementary Table 2). In moderate litters, 4/25 hearts were abnormal; in high litters, 26/32 hearts were abnormal; and in severe litters, 14/21 were abnormal; compared to 0/44 controls. Whereas STZ-induced diabetic models typically yield cardiac defects in 30–50% of offspring with substantial inter-litter variability (Nishino et al. 2023; Basu et al. 2017c), the βV59M model, upon selection of animals with hyperglycemia >20 mmol/L, results in cardiac abnormalities in 65-80% of the offspring (Table 1, Supplementary Table 2). This high penetrance highlights the robustness of our genetic approach for studying embryonic heart defects.

**Table 1.** Incidence of cardiac malformations across genotypes and maternal conditions, showing the highest incidence and broadest spectrum of defects when maternal hyperglycemia co-occurs with *Nkx2-5* haploinsufficiency. Percentages exceed 100% within one experimental group, because some hearts exhibited multiple defects simultaneously thereby counted in more than one category (ASD, atrial septal defect; AVSD, atrioventricular septal defect; DOLV, double outlet left ventricle; DORV, double outlet right ventricle; TGA, transposition of the great arteries; TOF, tetralogy of Fallot; VSD, ventricular septal defect; UV, univentricular heart).

| Types of defects | NG |  | HG |  |
| --- | --- | --- | --- | --- |
|  | Nkx2.5<sup>+/+</sup> | Nkx2.5<sup>GFP/+</sup> | Nkx2.5<sup>+/+</sup> | Nkx2.5<sup>GFP/+</sup> |
| ASD | 0 | 0 | 5 | 3 |
| AVSD | 0 | 0 | 3 | 0 |
| DOLV | 0 | 0 | 0 | 3 |
| DORV | 0 | 0 | 2 | 2 |
| TGA | 0 | 0 | 0 | 3 |
| TOF | 0 | 0 | 0 | 3 |
| muscvSD | 0 | 0 | 10 (29.4%) | 24 (60.0%) |
| mVSD | 0 | 0 | 2 | 3 |
| UV | 0 | 0 | 1 (3%) | 8 (23.5%) |
| Hypertrophy | 0 | 0 | 2 | 2 |
| Wall thinning | 0 | 0 | 8 (24%) | 28 (70%) |
| Total abnormal | 0 | 0 | 27 | 40 |
| Total normal | 0 | 0 | 7 | 0 |
| P value vs Nkx2.5<sup>+/+</sup> | - | - | - | p < 0.05 |

### Reversal of maternal hyperglycemia at E3.5 rescues developmental abnormalities

A unique property of the ßV59M+ hyperglycemia model is that the inhibition of insulin secretion caused by the β-cell-specific expression of a human activating KATP channel mutation is reversable by sulfonurea treatment (Brereton et al. 2014). Evidence in humans suggests that elevated maternal BGL alone causes offspring birth defects (Priest et al. 2015). We therefore used this feature of the model to test the effects of reducing severe hyperglycemia in early pregnancy prior to embryo implantation on subsequent embryonic development. 6 ßV59M+ females with severe hyperglycemia and 3 ßV59M-controls (normal BGL) were timed mated, then implanted with a glibenclamide pellet at E3.5. BGLs decreased to normal in 48 hours (14.8 mmol/l, Supplementary Fig. 2H). 5/6 implanted diabetic and 3/3 implanted control females were still pregnant at litter collection. Embryo BGL in severe-reversed embryos was not significantly difference to controls, indicating a dramatic reduction compared to embryos from untreated mothers with severe hyperglycemia (Supplemental Fig. 2D). 13/18 embryos from these litters were normal, 4/18 abnormal and 1/18 were dead compared to 21/22 normal and 1/22 abnormal embryos from the control reversed mothers (Supplemental Table 1, Supplemental Fig. 2F). This is a highly significant reduction of death and abnormalities compared to offspring from mothers with severe hyperglycemia. Heart morphology was examined in a subset of these embryos: 3/14 embryos from reversed severe hyperglycemia mothers had heart defects, a significant drop from embryos from mothers with severe hyperglycemia, although rescue was not complete (Supplemental Table 2). These data support the hypothesis that maternal hyperglycemia by itself is teratogenic.

### Gene-environment interaction (G×E) between maternal hyperglycemia and offspring *Nkx2-5* haploinsufficiency

We next asked whether *in utero* hyperglycemic exposure of embryos with a genetic susceptibility to CHD increases the effects of either factor alone on cardiac development and risk of heart defects. Tamoxifen-injected ßV59M+ females with blood glucose levels >20 mmol/L and control normoglycemic females (Fig. 2A-B) were mated with *Nkx2-5^GFP/+^* males, generating four groups of embryos: normoglycemic *Nkx2-5^+/+^* (Control); normoglycemic *Nkx2-5^GFP/+^* (Gene); hyperglycemic *Nkx2-5^+/+^*(Environment); and hyperglycemic *Nkx2-5^GFP/+^* (G×E) (Fig. 2A). Similarly to our results above, we observed a significantly increased foetal mortality from E16.5 in foetuses from hyperglycemic mothers compared to those from normoglycemic mothers (Supplementary Fig. 3). Foetal genotype did not influence this temporal pattern, with no earlier mortality detected, statistically (Supplementary Table 3). Using HREM at E14.5-E17.5, we found that combining genetic predisposition with maternal hyperglycemia increased both prevalence and severity of CHD, with penetrance increasing from 79.4% (27/34) in the environment group (*n* = 34) to 100% (40/40) in the G×E group (*n* = 40) (Table 1). However, no increase was observed in the frequency of ASD (3/40 *vs* 5/34) AVSD (0/40 *vs* 3/34), DORV (2/40 *vs* 2/34) and hypertrophy (2/34 *vs* 2/40) (Fig. 2C, Table 1). Compared with hyperglycemic *Nkx2-5^+/+^* foetal hearts (Environment), the G×E group showed more frequent muscular VSD (24/40 60% *vs* 10/34 29,4%) a higher rate of univentricular morphologies (8/40 23.5% *vs* 1/34 3%), and markedly increased chamber dilation with wall thinning (28/40 70% *vs* 8/34 24%, Fig. 2C, Table 1). These categories often co-occurred within the same heart, accounting for totals exceeding 100% within groups. Together these data show that foetal genetic susceptibility and altered maternal environment have a synergistic effect compromising cardiac development and elevating heart defect risk.

**Fig. 2.**
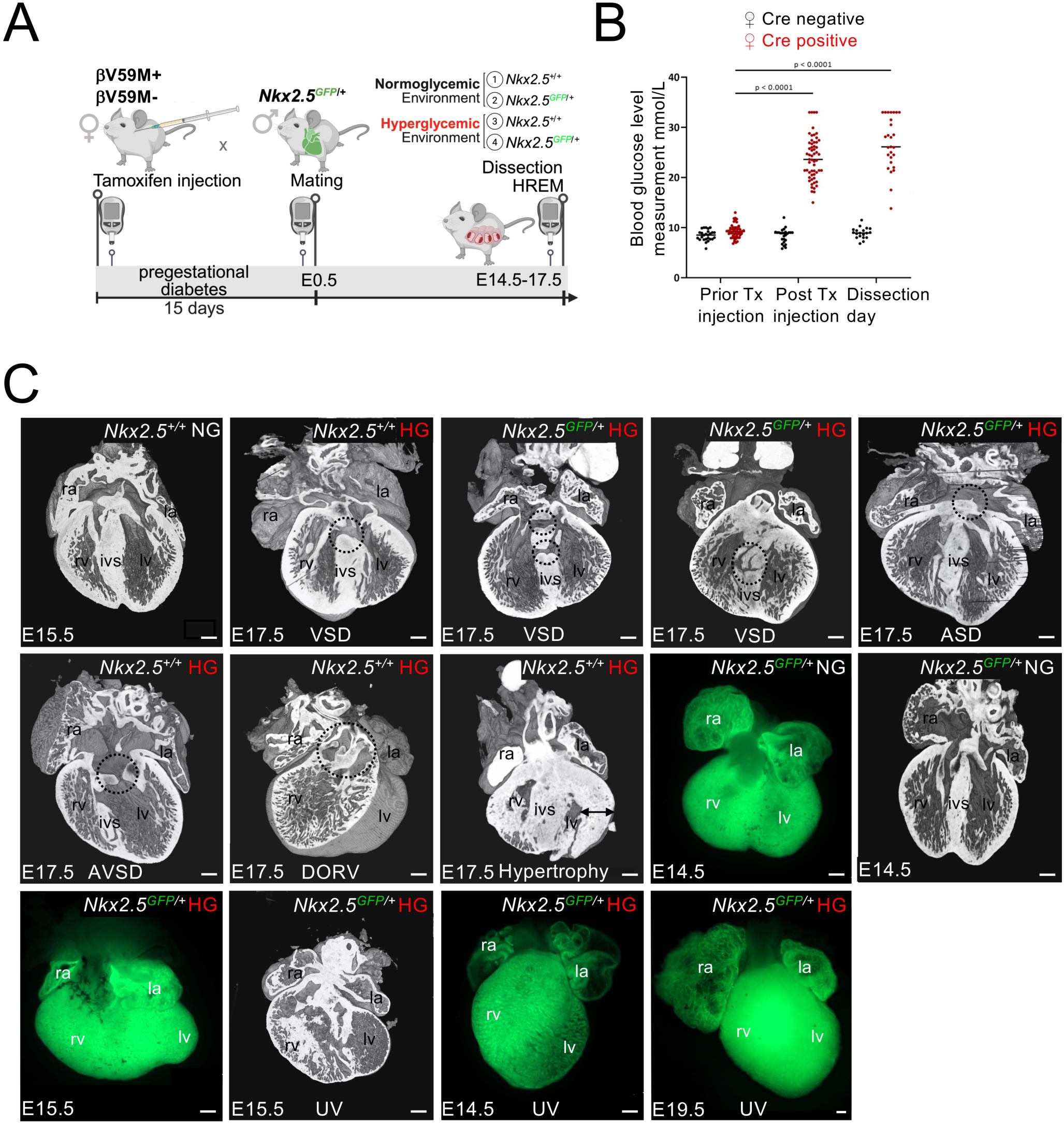
Increased susceptibility to specific cardiac malformations in *Nkx2-5* haploinsufficient foetuses exposed to maternal hyperglycemia. (A) Schematic representation of the experimental design and breeding strategy. Diabetes was induced in female βV59M mice by tamoxifen injection. Fifteen days after induction, diabetic females were mated with males heterozygous for *Nkx2-5* (*Nkx2-5^GFP/+^*) to generate foetuses with genetic susceptibility to CHD and exposed *in utero* to hyperglycemia. Foetuses were collected between E14.5 and E17.5 for HREM-based analysis of cardiac morphology. Elements of this panel were created in BioRender.com (https://BioRender.com/i72e213). (B) Blood glucose measurements confirmed that Cre-positive females became hyperglycemic (HG) after tamoxifen induction, while Cre-negative controls remained normoglycemic (NG). (C–N) Representative HREM images illustrating the spectrum of cardiac malformations observed under different maternal (NG or HG) and genetic conditions (*Nkx2-5^+/+^* or *Nkx2-5^GFP/+^*), including atrial septal defect (ASD), atrioventricular septal defect (AVSD), ventricular septal defect (VSD), double outlet right ventricle (DORV), univentricular (UV) morphology, hypertrophy, and dilated chambers with wall thinning; Scale bars: 200 µm.

### G×E drives transcriptional changes affecting the contractile identity of chamber myocardium

For molecular characterization of the observed phenotypes, we performed bulk RNA-seq on E15.5 whole hearts from the 4 experimental groups, normoglycemic *Nkx2-5^+/+^* (Control), hyperglycemic *Nkx2-5^+/+^* (Environment), normoglycemic *Nkx2-5^GFP/+^* (Gene), and hyperglycemic *Nkx2-5^GFP/+^* (G×E) (*n*=3 per group, for a total of 12 samples, Supplementary Fig. 3). We analyzed four comparisons: Environment *vs* Control; Gene *vs* Control; G×E *vs* Environment; and G×E *vs* Gene. Principal component analysis (PCA) of the bulk RNA-seq data, performed on biological triplicates, revealed some dispersion among replicates across all experimental groups, including G×E, Environment, Gene, and Control. This variability likely reflects natural biological differences between individual foetal hearts, which were obtained from different litters. Importantly, despite this intra-group variability, the overall global trends and separation between experimental groups remain clearly detectable (Supplementary Fig. 3A). Data revealed widespread transcriptional alteration, especially when maternal hyperglycemia co-occurred with embryonic *Nkx2-5* haploinsufficiency (G×E). Skeletal muscle contraction was found within the top10 terms for upregulated genes when comparing G×E to Environment.

GO terms related to cellular respiration and mitochondrial ATP production were among the most recurrent in the Environment *vs* Control, G×E *vs* Gene, and G×E *vs* Control comparisons. Their enrichment in these comparisons is consistent with altered metabolic gene expression in embryonic hearts exposed to maternal hyperglycemia (Fig. 3A, Supplementary Table 4). At the gene level, differentially expressed genes (DEG) analysis (Fig. 3B) and normalized counts (Fig. 3G-I-K) showed *Tnni2, Tnnt3* upregulation and *Tnnt1* downregulation due to G×E (Supplementary Fig. 3B). The altered expression of troponin genes likely reflects a shift from a typical cardiac program toward a more skeletal muscle–like transcriptional profile. RNAScope at E14.5 confirmed this pattern *in situ*: in the G×E group we observed loss of *Tnnt1* (Fig. 3H) and gain of *Tnni2* (Fig. 3J), especially in the right ventricular myocardium and left atria, while *Tnnt3* (Fig. 3L) was upregulated in both atria. Together, these data indicate coordinated shifts in myofibrillar identity and cellular respiration, reflecting altered myocardial phenotype. To determine whether this deregulation occurs at earlier stages, we examined *Tnni2* and *Tnnt3* expression at E10.5, but no comparable changes were detected (Supplementary Fig. 3C). We also assessed *Hand1* expression to distinguish the left from the right ventricular myocardium and observed reduced *Hand1* expression in the left ventricle (Supplementary Fig. 3C). We also assessed E12.5 hearts in wild-type conditions to rule out the possibility that the observed expression patterns at E14.5 result from persistent early expression or developmental delay. At E12.5, *Tnnt3* was absent and *Tnni2* was only very weakly expressed (data not shown), supporting the conclusion that the observed shifts at E14.5 are specific to the G×E condition rather than a consequence of delayed myocardial maturation.

**Fig. 3.**
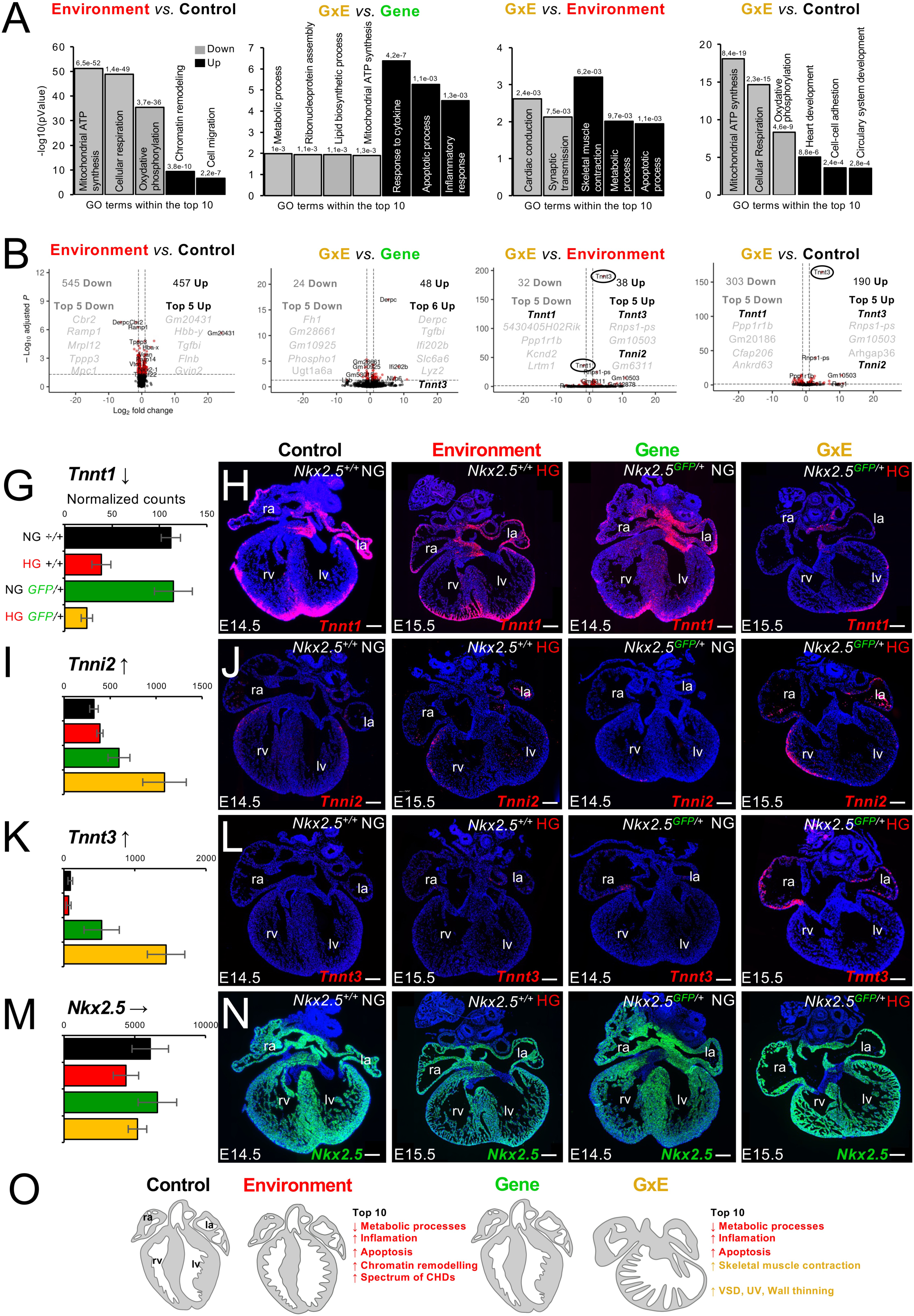
Transcriptomic alterations in foetal hearts reveal gene-environment dependant alteration of contractile identity. Bulk RNA-seq was performed on E15.5 whole hearts from four experimental groups: normoglycemic (NG) *Nkx2-5^+/+^* (black, Control group), hyperglycemic (HG) *Nkx2-5^+/+^* (red, Environment), NG *Nkx2-5^GFP/+^* (green, Gene group), and HG *Nkx2-5^GFP/+^* (yellow, G×E group). Each group included three biological replicates (hearts), for a total of 12 samples. Five pairwise comparisons were analyzed: environment *vs*. control, gene *vs*. control, G×E *vs*. environment, G×E *vs*. gene and G×E *vs*. control. (A) Gene Ontology (GO) enrichment showing representative upregulated (black) and downregulated (grey) terms within the top 10 for each of 4 comparisons. GO analysis was performed using (Enrichr); only terms meeting adjusted *P* < (0,05) are shown. (B) Volcano plots displaying significantly upregulated (right) and downregulated (left) transcripts and highlighting the top five genes of each category per comparison. Differentially expressed genes (DEGs) were called using DESeq2 with FDR < 0.05 (Benjamini–Hochberg; (Wald test)). (G, I, K, M) Normalized read counts of *Tnnt1*, *Tnni2*, *Tnnt3*, which recur among the most deregulated genes across comparisons, and of *Nkx2-5*. Values are (DE DESeq2)-normalized counts, bars/lines indicate mean ± SD. (H, J, L, N) Spatial validation by RNAScope on E14.5-15.5 heart sections confirming downregulation of *Tnnt1* (H), upregulation of *Tnni2* (J) and *Tnnt3* (L) in the G×E group, while *Nkx2-5* (N) expression remained unchanged. Scale bars: 200 µm.

### Developmental origins of the observed malformations: G×E alters second heart field patterning, impairs progenitor proliferation and survival, and remodels chromatin accessibility

To investigate how maternal hyperglycemia reshapes transcriptional and chromatin states during early cardiac development, particularly on a sensitized *Nkx2-5* background, we performed a multiome analysis (RNA +ATAC) on micro-dissected embryonic cardiac regions at E9.5 from all four experimental groups (2 technical replicates per group, 8 libraries in total) (Fig. 4A). We removed the head folds and posterior trunk via microdissection to increase the proportion of cardiac progenitors captured for sequencing. Isolated nuclei were processed using single nucleus Multiome workflow, yielding paired gene expression (snRNA-seq) and chromatin accessibility (snATAC-seq) measurements from the same tissues. Quality control analyses demonstrated good coverage in ATAC and RNA values, robust TSS enrichment and ATAC counts (Supplementary Fig. 4), validating the quality of our snATAC-seq data. Integration of transcriptomes with ATAC-derived gene-activity score, graph-based unsupervised clustering produced a comprehensive cellular atlas (Fig. 4B, Supplementary Table 5). Major cell types were annotated based on an extensive set of known marker genes. For clarity, only a few representative markers are shown (Fig. 4C,D, Supplementary spreadsheet 1), including cardiomyocytes (*Nkx2-5, Ttn*), cardiac progenitors (*Aldh1a2, Tbx1*), endocardium (*Kdr*), pro/epicardium (*Tbx18, Wt1*), neural crest-derived cells (*Tfap2b*), fibroblast-like cells (*Col1a1*), and red blood cells (*Hbb-y*). A finer subclustering further subdivided the cardiomyocyte cluster into outflow tract (OFT), atrioventricular canal (AVC), atrial, ventricular, and sinus venosus (SV) domains (Fig. 4B). These data confirm that all expected cell types from our micro-dissected embryonic hearts were captured in the snMultiome experiment, providing a cellular map to trace early deregulations within specific cell types and link them to the observed structural defects.

**Fig. 4.**
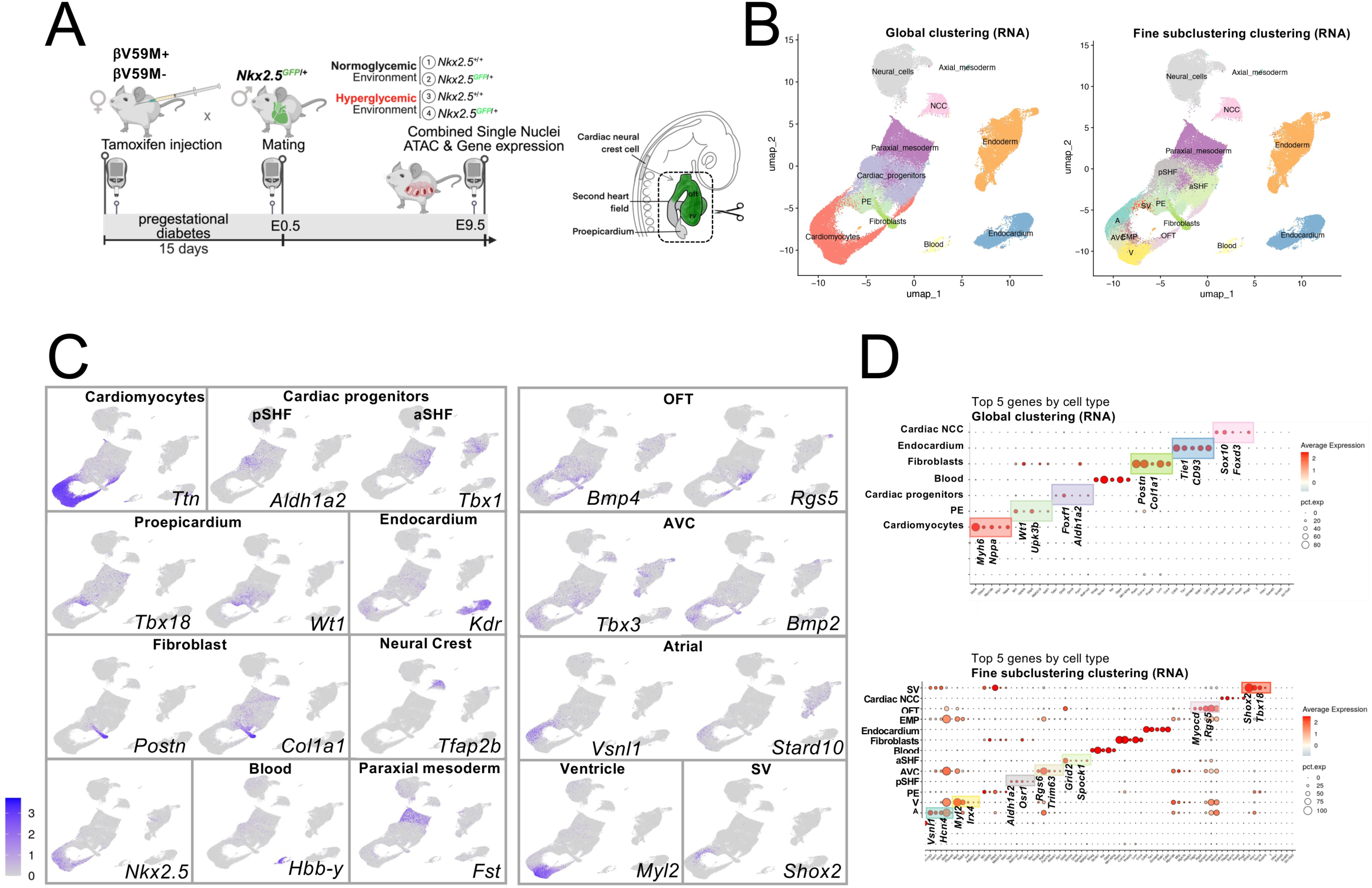
Single-nucleus Multiome at E9.5 maps cardiac lineages and reveals G×E-dependent shifts in cardiac gene expression. (A) Schematic representation of the experimental design and the microdissected embryonic cardiac region used for nuclei isolation used for single-nucleus Multiome (RNA+ATAC) sequencing. Diabetes was induced in female βV59M mice by tamoxifen. Fifteen days later, females were mated with *Nkx2-5^GFP/+^*males. Embryos were collected at E9.5, carefully staged to ensure the same number of somites (22-24 somites), and belonged to four experimental groups: normoglycemic (NG) *Nkx2-5^+/+^* (Control), hyperglycemic (HG) *Nkx2-5^+/+^*(Environment), NG *Nkx2-5^GFP/+^* (Gene), and HG *Nkx2-5^GFP/+^* (G×E). (B) Two-dimensional UMAP representation of single-nucleus transcriptomes from all experimental groups. A global graph-based clustering delineates the principal cardiac lineages and anatomical regions. A second, high-resolution UMAP representation highlights fine subclustering of selected populations. NCC, cardiac neural crest cells; PE, proepicardium; A, atria; V, ventricles; aSHF, anterior second heart field; pSHF, posterior second heart field; AVC, atrioventricular canal; OFT, outflow tract; SV, sinus venosus; EMP, early myocardium progenitors. (C) Marker-based annotation of clusters/subclusters using selected lineage and cell-specific markers (examples shown). (D) Dot plot of the top five markers for each identified cluster. For each cluster, the relative gene expression in percent is represented by the size of dots, e.g., a value of 100 means that each cell within this cell type expressed this gene. The average expression level for the listed genes is indicated by the color.

We first sought to investigate the molecular origins of the structural defects by examining the effects of G×E on cardiac progenitor cells. A higher proportion of aSHF cells was observed in the G×E group compared with the Gene, Environment, and Control groups, whereas a reduced proportion of pSHF cells was seen in the G×E *vs* Gene comparison, indicating that SHF progenitor patterning is altered under G×E conditions (Supplementary Fig. 5A,B).

More genes were downregulated than upregulated in both aSHF and pSHF cells across the G×E *vs.* Control, Environment *vs.* Control, and Gene *vs.* Control comparisons (Supplementary spreadsheet 2 and 3). This might suggest that G×E preferentially suppresses developmental gene expression programs (Supplementary Fig. 6). Comparing G×E *vs* Control, Environment *vs* Control, and Gene *vs* Control, the top 10 downregulated GO terms in the aSHF and pSHF included “chromatin remodeling”. The genes driving this enrichment included *Eed, Hdac1, Hdac2, Sirt1, Sin3A,* and *Trim28* (Supplementary Fig. 5C, Supplementary Fig. 6, Supplementary Table 5). This might suggest that the G×E condition triggers a reorganization of chromatin accessibility. Downregulation of the GO term “Chromatin remodeling” was also observed in the pSHF, suggesting that chromatin remodeling is broadly perturbed across cardiac progenitors. This result is consistent with the context-dependent function of NKX2-5, a transcription factor capable of acting as either an activator or a repressor in SHF cells (Watanabe et al. 2012b). For both the G×E *vs.* Control and Environment *vs.* Control comparisons, the top 10 downregulated GO terms shared by aSHF and pSHF included “chromosome segregation,” “regulation of chromosome organization,” “regulation of cell-cycle phase transition,” and “sister chromatid segregation.” Together, these terms indicate reduced expression of genes involved in cell-cycle progression and mitosis, consistent with decreased proliferative activity in both progenitor populations. This finding is particularly relevant because appropriate proliferation of SHF progenitors, partly regulated by FGF signalling, is essential for normal OFT and RV development (Rochais et al. 2014; Shi et al. 2016; Watanabe et al. 2010; Marguerie et al. 2006). Accordingly, reduced SHF progenitor proliferation has been associated with structural abnormalities similar to those observed in our G×E model, including membranous and muscular VSDs, myocardial thinning, and trabecular defects. Consistent with a reduced progenitor-cell pool, in the pSHF, apoptosis-related GO terms were among the top 10 terms enriched in upregulated genes in the G×E *vs.* Control comparison. This was not observed in the top 10 terms for the Environment *vs.* Control or Gene *vs.* Control comparisons, or in the aSHF. Increased apoptosis of SHF progenitors has previously been associated with impaired ventricular development and cardiac malformations, including a thin ventricular myocardium and ventricular septal defects (C. Leung et al. 2014; Tsuchihashi et al. 2011). These findings raise the possibility that increased apoptosis within the SHF may contribute to the ventricular and septal abnormalities observed in our model. Interestingly, in the aSHF comparison of G×E *vs* Control, the top 10 downregulated GO terms included genes associated with WNT signaling, such as *Fzd3, Fzd4, Fzd7, Gsk3a,* and *Wnt5b*, as well as *Isl1* and *Fgf10*, indicating a perturbation of these genes specific to the aSHF (Supplementary table 5). This finding is particularly relevant given the established role of WNT signaling in regulating SHF progenitor expansion and the development of its cardiac derivatives. Conditional disruption of canonical WNT/β-catenin signaling in the SHF impairs right ventricular and interventricular myocardial expansion, while perturbation of non-canonical WNT signaling through loss of *Wnt5a* results in ventricular septal defects and abnormal outflow tract development (Li Chen et al. 2012; Ai et al. 2007; Cohen et al. 2012).

### G×E alters chromatin accessibility in cardiac progenitors at loci associated with developmental and stress-response pathways

To gain insight into the regulatory mechanisms underlying these transcriptional changes, we next analyzed the scATAC-seq dataset. To assess whether chromatin accessibility distinguished the cell populations identified by scRNA-seq, we examined specific accessible regions in the scATAC-seq dataset. The heatmap revealed distinct accessibility patterns across the identified populations, including cardiac progenitors and cardiomyocytes (Supplementary Fig. 7A). GREAT analysis of regions assigned to the cardiac progenitor cluster identified terms related to development and morphogenesis, including tissue morphogenesis, embryonic morphogenesis, tube morphogenesis, and heart development (Supplementary Fig. 7B, Supplementary Table 6). Transcription factor motif analysis by HOMER using the JASPAR motif database revealed distinct regulatory signatures across embryonic cardiac cell populations (Supplementary Fig. 7D-C). Motif analysis of these regions identified MEIS, Sp1, and FOX motifs among the most enriched (Supplementary Fig. 7C). Consistent with distinct regulatory profiles, targeted motif analysis showed enrichment of MEF2 and GATA motifs in cardiomyocytes (Supplementary Fig. 7D). Cardiac progenitors were specifically enriched for motifs recognized by the MEIS, Sp/KLF, GATA, HAND, and T-box transcription factor families (Supplementary Fig. 7D). This is consistent with established cardiac developmental regulatory networks (Li Chen et al. 2009; Darieva et al. 2025; Nemer and Horb 2007; Zhou et al. 2017; Tsuchihashi et al. 2011) and further validates the biological relevance of the scATAC-seq dataset.

As *Nkx2-5* is expressed in aSHF progenitors, we first performed functional enrichment analysis of the DARs identified in this cell population. Analysis of all DARs in the G×E versus Control comparison, including both regions with increased (logFC > 0) and decreased (logFC < 0) accessibility, identified 1,839 DARs in aSHF cells (Figure 5A). Among these, 279 regions (15.2%) displayed increased accessibility, whereas 1,560 regions (84.8%) showed decreased accessibility, indicating a predominant loss of chromatin accessibility under G×E conditions (Supplementary Fig. 8A). In the aSHF, integration of transcriptional changes with differentially accessible regions containing NKX2-5 motifs highlighted *Rhoa*, *Rock2*, and *Rian* as candidate regulatory targets. *Rhoa* and *Rock2* are of particular interest because RHO–ROCK signaling controls second heart field epithelial organization (Francou et al. 2017). *Rian* encodes an imprinted long noncoding RNA within the *Dlk1–Dio3* locus, which has been implicated in embryonic heart development (Teng et al. 2024; T. Gu et al. 2012). Although *Rhoa*, *Rian*, and *Rock2* showed both transcriptional and chromatin accessibility changes, their associated regions did not rank among the 15 regions with the largest accessibility changes. We therefore prioritized loci with stronger chromatin accessibility changes for further analysis.

**Fig. 5.**
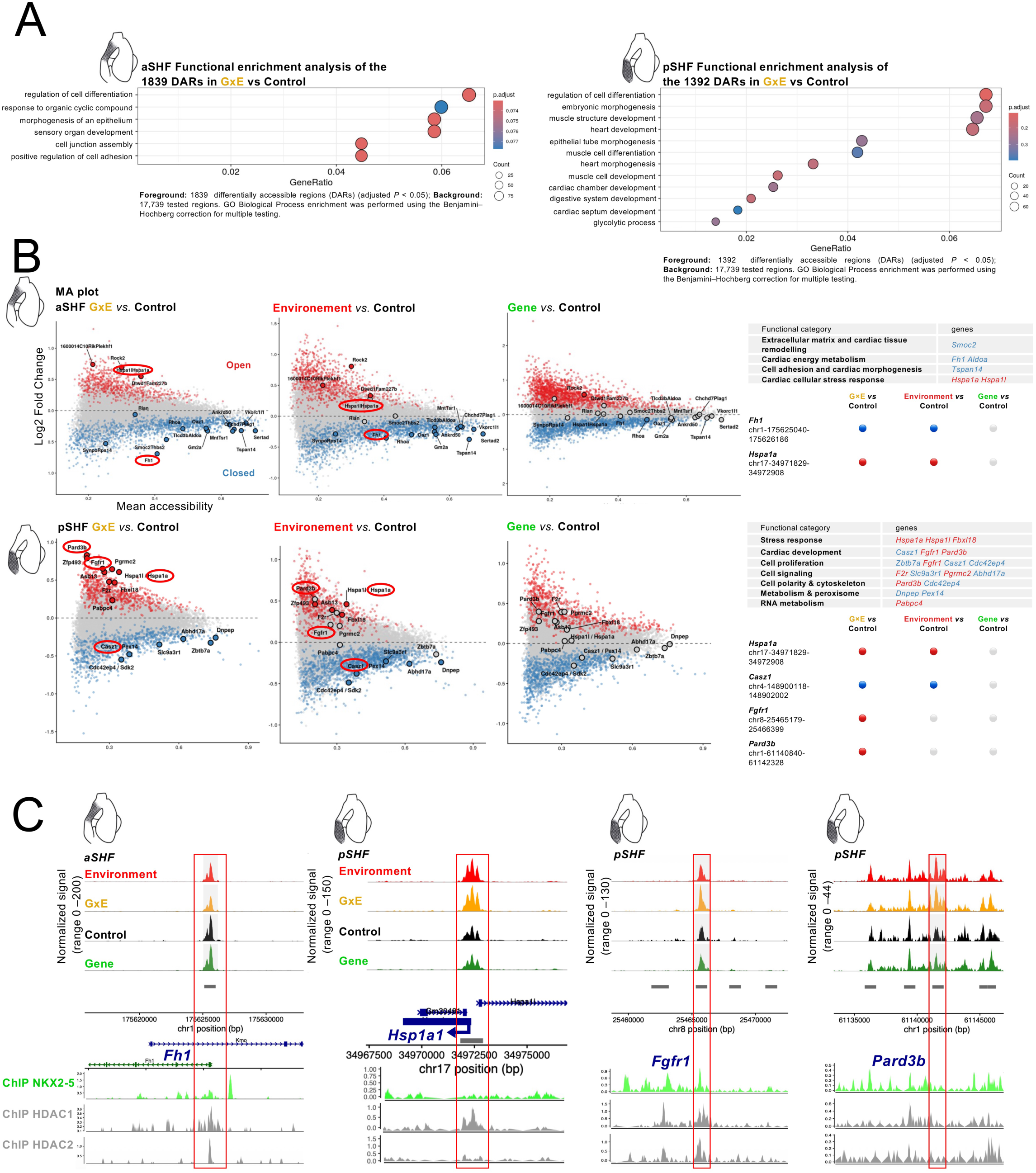
Multi-omic integration identifies candidate regulatory regions selectively remodeled by the interaction between maternal hyperglycemia and *Nkx2-5* haploinsufficiency in SHF progenitors. (A) Gene Ontology (GO) enrichment analysis of differentially accessible regions (DARs) identified in aSHF and pSHF progenitor populations from G×E embryos compared with controls. (C) MA plots showing differential chromatin accessibility across all pairwise comparisons in aSHF and pSHF progenitors. Regions with increased accessibility are shown in red and regions with decreased accessibility in blue (adjusted *P* < 0.05). Comparison across experimental groups identifies *Fh1, Hspa1a, Casz1, Fgfr1* and *Pard3b* as representative loci exhibiting condition-specific chromatin accessibility changes. (D) Genome browser tracks showing representative candidate regulatory regions associated with *Hspa1a*, *Casz1*, *Fgfr1* and *Pard3b*. DARs identified in GxE progenitors are displayed together with previously published NKX2-5, HDAC1/2 ChIP-seq tracks. Red boxes highlight the DARs shown in the analysis.

The pSHF was also examined to determine whether G×E-induced regulatory alterations extended beyond the principal *Nkx2-5*-expressing domain. Filtering pSHF DARs for NKX2-5 motifs failed to identify candidate direct NKX2-5 regulatory targets associated with transcriptional changes (Supplementary Fig. 8A). The absence of such candidates suggests that the transcriptional and chromatin changes observed in the pSHF predominantly arise through indirect or non-cell-autonomous mechanisms, consistent with *Nkx2-5* expression being largely restricted to the aSHF during early cardiogenesis (Watanabe et al. 2012b; O. W. Prall et al. 2008). Functional annotation of aSHF DARs highlighted biological processes related to cell differentiation, epithelial morphogenesis, cell-junction assembly and cell adhesion (Fig. 5A). In the pSHF, DAR-associated genes were linked to broader developmental processes, including embryonic and cardiac morphogenesis, muscle differentiation, cardiac chamber and septum development, and glycolysis (Fig. 5A).

Comparison of accessibility changes across the G×E *vs.* Control, Environment *vs.* Control and Gene *vs.* Control enabled discrimination of environmentally driven alterations from those preferentially associated with the G×E condition (Fig. 5B). In the aSHF, reduced accessibility at a region associated with *Fh1*, a gene involved in mitochondrial energy metabolism (Ashrafian et al. 2012), and increased accessibility near the stress-response gene *Hspa1a* (Kim et al. 2006) were detected in both G×E *vs.* Control and Environment *vs.* Control comparaisons, but not in Gene *vs.* Control. This suggests that these alterations are primarily associated with maternal hyperglycaemia. Additional candidate DARs were associated with genes involved in extracellular-matrix remodelling, and cell adhesion, including *Smoc2* and *Tspan14* (Supplementary spreadsheet 4).

A similar pattern was observed in the pSHF, where the *Hspa1a*-associated region displayed increased accessibility, whereas a region associated with the cardiac developmental regulator *Casz1* (Kennedy et al. 2017) showed reduced accessibility in both G×E and Environment embryos. By contrast, increased accessibility at regions associated with *Fgfr1* and *Pard3b* was detected specifically in the G×E *vs.* Control comparison. Increased accessibility at regions associated with *Fgfr1* and *Pard3b* was detected specifically in the G×E *vs.* Control comparison. *Fgfr1* is relevant to FGF signalling, which contributes to second heart field development (Park et al. 2008). *Pard3b*, a gene associated with epithelial cell polarity (Kohjima et al. 2002), points instead to a potential change in progenitor organisation, a process important for second heart field morphogenesis (Francou et al. 2014). Together, these loci suggest distinct regulatory changes in the G×E condition, whose functional consequences remain to be determined. These genes contribute to developmental signalling, polarity and cytoskeletal organisation, suggesting that their altered regulatory landscapes may reflect molecular effects arising specifically from the interaction between maternal hyperglycaemia and *Nkx2-5* haploinsufficiency. Genome browser views illustrate representative candidate regulatory regions associated with *Fh1, Hspa1a*, *Fgfr1*, *Pard3b* (Fig. 5C) and *Casz1* (Supplementary Fig. 8). These loci recapitulate the chromatin accessibility changes identified by differential accessibility analysis, including increased accessibility at the *Hspa1a*, *Fgfr1* and *Pard3b* loci and reduced accessibility at the *Fh1* and *Casz1* loci in GxE embryos. The DAR at the *Fgfr1* locus overlaps previously published NKX2-5, HDAC1, and HDAC2 ChIP-seq peaks. Although none of these regulatory regions fulfilled our criteria for high-confidence direct NKX2-5 targets, all four genes have previously been implicated in biological processes relevant to cardiac development or stress adaptation. *Hspa1a* encodes a stress-inducible molecular chaperone that protects cardiomyocytes from proteotoxic and oxidative stress (Willis and Patterson 2010; Benjamin and McMillan 1998; Xiao and Benjamin 1999). *Casz1* is an essential regulator of cardiomyocyte differentiation and cardiac morphogenesis, controlling ventricular wall development and myocardial compaction (Sojka et al. 2014; Z. Liu et al. 2014; Dorr et al. 2015). *Fgfr1* mediates FGF signaling required for second heart field maintenance and outflow tract development (J. Zhang et al. 2008; Park et al. 2008). Finally, *Pard3b*, a component of the PAR polarity complex, contributes to epithelial polarity and tissue morphogenesis and has recently been implicated in cardiomyocyte polarization and cardiac morphogenesis (Francou et al. 2017; Kamakura et al. 2024). Collectively, these candidate loci suggest that the G×E condition perturbs metabolic and stress-response pathways, as well as developmental signalling and cell-polarity programmes, largely through regulatory mechanisms that do not involve direct NKX2-5 binding.

### Multi-omic integration identifies an indirect *Nkx2-5–Aldh1a2* regulatory axis disrupted by maternal diabetes in posterior second heart field progenitors

Within the pSHF, 193 genes were downregulated and 60 were upregulated in the G×E versus Control comparison (Supplementary Fig. 6). Among the downregulated genes, *Aldh1a2* was of particular interest because it encodes a key retinoic acid-synthesizing enzyme during embryogenesis, and retinoic acid signaling plays an important role in the patterning of second heart field progenitors (Sirbu et al. 2008; Ryckebusch et al. 2008). Moreover, maternal diabetes and hyperglycemia have previously been shown to perturb embryonic retinoic acid homeostasis and modify susceptibility to retinoic acid signaling (Nishino et al. 2023; Chan et al. 2002; M. B. W. Leung et al. 2004; Lee et al. 2017). We therefore examined *Aldh1a2* expression more closely and found that it was significantly downregulated in the pSHF in the G×E condition (Fig. 6A). This gene encodes the enzyme retinaldehyde dehydrogenase 2 (RALDH2) involved in retinoic acid synthesis. The RA pathway is essential for OFT development and myocardial trabeculation (Rhinn et al. 2011; Niederreither et al. 2001). To validate these results, RNAscope was performed on E9.5 whole-mount embryos (ventral view) and lateral sections. The aSHF domain was marked with *Nkx2-5*, and the pSHF domain with *Osr1* and *Aldh1a2*. *Aldh1a2* expression was reduced in the pSHF in the G×E group (Fig. 6B–C). In the G×E condition, the aSHF domain abnormally overlapped with the pSHF, extending to the venous pole at the level of the atrium, unlike in the Environment and Control groups (Fig. 6B, yellow bar) (Fig 6B, indicated by the yellow bar). At this developmental stage, the aSHF and pSHF domains are normally spatially distinct. However, *Aldh1a2* expression in the pSHF is rapidly extinguished by E10.5, therefore embryos were carefully stage-matched based on somite number to ensure accurate comparison of developmental stages between experimental groups. A similar, albeit less pronounced, overlap was observed in the Environment condition in the ventral view (Fig. 6B), consistent with previous reports in an STZ-induced maternal diabetes model. Notably, in the G×E condition, the *Nkx2-5-*positive aSHF domain abnormally overlapped with the *Aldh1a2*-positive pSHF domain, extending towards the venous pole at the level of the atrium (Fig. 6B–C). ScATAC-seq analysis revealed a strong chromatin accessibility signal at the *Aldh1a2* locus, centred on its promoter (Fig. 6D). Although this region remained accessible across all experimental conditions, differential accessibility analysis showed reduced accessibility in G×E cells compared with each of the Control, Gene, and Environment groups (Fig. 6D; This quantitative rather than binary change is consistent with previous single-cell analyses showing that cardiac progenitors progress through transitional developmental states accompanied by dynamic and gradual changes in chromatin accessibility (Ameen et al. 2022). The residual accessibility observed under G×E conditions may therefore reflect the developmental plasticity and heterogeneous commitment states of these embryonic progenitors, rather than complete promoter closure across the entire cell population.

**Fig. 6.**
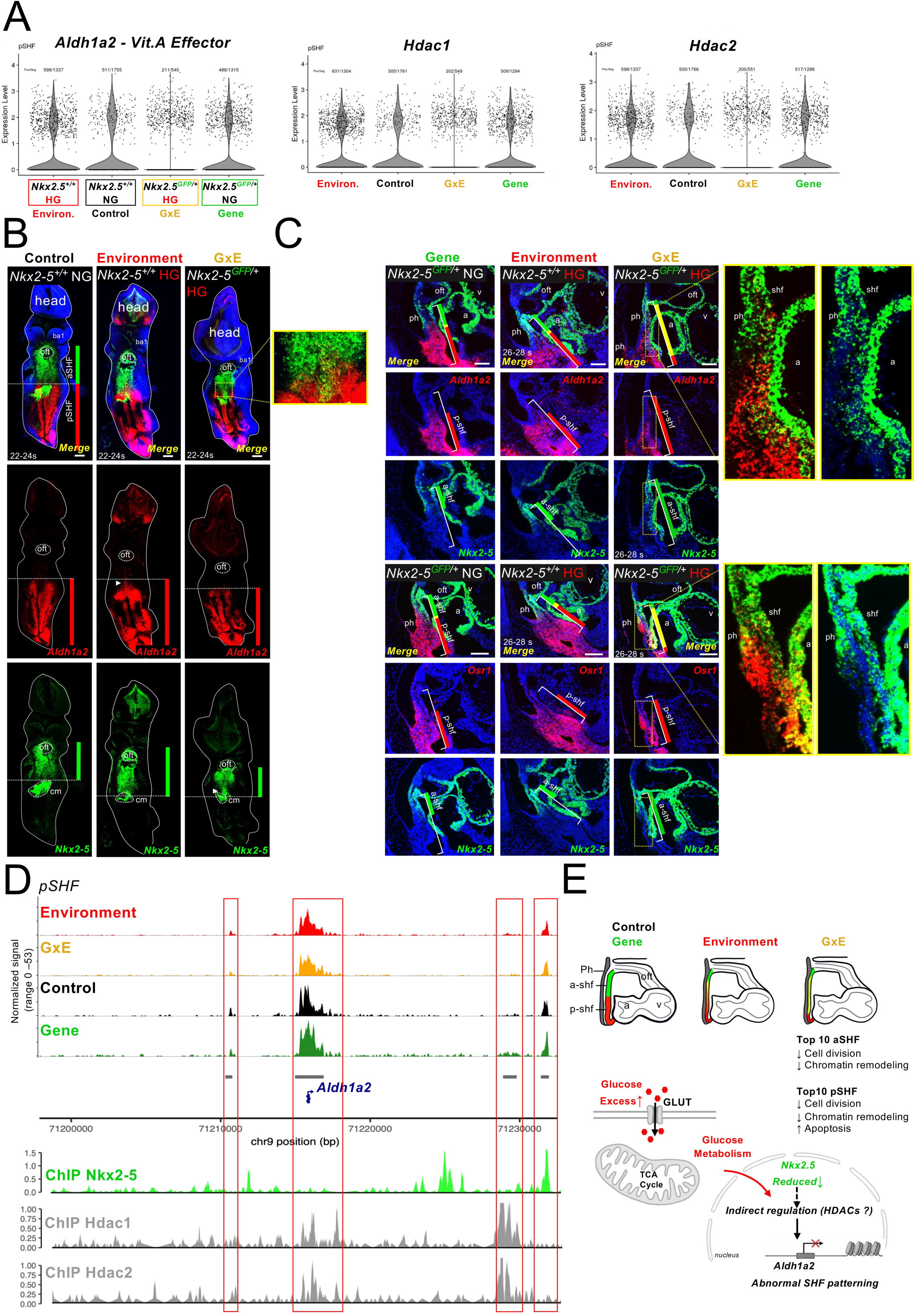
Second heart field programs are perturbed under G×E conditions, involving the retinoic-acid pathway. (A) Violin plots for *Aldh1a2*, *Hdac1*, and *Hdac2* within the posterior second heart field (pSHF) subcluster. (B) Spatial localization of *Aldh1a2* by RNAScope in E9.5 hearts. Whole-mount ventral views (22–24 somites) and (C) lateral sections with *Nkx2-5* co-detection (26–28 somites) highlight SHF patterning across Control, Environment, Gene, and G×E conditions. (D) Genome browser view of the *Aldh1a2* locus showing chromatin accessibility profiles across the four experimental groups, together with single-cell RNA-seq expression data and previously published NKX2-5 and HDAC1/2 ChIP-seq datasets. Candidate regulatory regions identified in this analysis are highlighted by red boxes. (E) Schematic summary of altered SHF patterning and the proposed underlying mechanism. Maternal hyperglycemia, combined with *Nkx2-5* haploinsufficiency (G×E), is associated with abnormal overlap of the aSHF domain with the pSHF, extending to the venous pole at the level of the atrium, and reduced distance between the pharyngeal endoderm and the SHF.Transcriptional changes include downregulation of genes involved in cell division and chromatin remodeling in both populations, and upregulation of apoptosis-related genes in the pSHF. The proposed model links excess glucose and altered glucose metabolism to reduced *Nkx2-5* expression and altered *Aldh1a2* regulation, potentially involving HDACs, thereby contributing to abnormal SHF patterning. Ph, pharynx; oft, outflow tract; a, atrium; v, ventricle; GLUT, glucose transporter; TCA, tricarboxylic acid.

The shortened aSHF and its abnormal overlap with the *Aldh1a2*-positive pSHF domain suggest defective segregation of anterior and posterior SHF identities. In mouse embryos, initially overlapping progenitor domains normally segregate during heart tube elongation, a process regulated by RA signaling (De Bono et al. 2018). Consistent with this role, RA deficiency results in posterior expansion of aSHF markers and defective ventricular trabeculation (Sirbu et al. 2020; Niederreither et al. 2001; Ryckebusch et al. 2008). Thus, disruption of RA-dependent SHF patterning may contribute to both the early progenitor-domain abnormalities and the later ventricular defects observed in the G×E condition. We also detected reduced chromatin accessibility at the *Sin3a* locus in cardiac progenitors from the G×E group compared with each of the other three groups. (Supplementary Fig. 8D). SIN3A is a scaffold protein that associates with HDAC1/2 in transcriptional repressor complexes. Separately, SIN3A has been shown to interact with TET1 and facilitate transcription of shared target genes in mouse embryonic stem cells (F. Zhu et al. 2018; Chandru et al. 2018). The functional consequence of the accessibility change at *Sin3a* in cardiac progenitors remains to be determined. Existing ChIP-seq data shows NKX2-5 binding at the *Sin3a* promoter in murine embryonic hearts ((Williams et al. 2011), Supplementary Fig. 8E); and SIN3A and its cofactors HDAC1 and TET1 can bind to the *Aldh1a2* promoter (Supplementary Fig. 8E). These data suggest that dysregulated recruitment of HDAC/TET1/SIN3A to the *Aldh1a2* locus, resulting from the combined effects of *Nkx2-5* haploinsufficiency and hyperglycemia, may alter gene expression and disrupt SHF organization (Supplementary Fig. 8F). In humans, SIN3A deficiency is known to cause Witteveen-Kolk syndrome, which manifests with cardiovascular anomalies including thickened aortic valve, atrial septal defects, patent *ductus arteriosus*, and patent *foramen ovale* (Coenen-van der Spek et al. 2023; Uctepe et al. 2024). In addition, sequence variants in *SAP130*, which encodes a SIN3A-associated protein in the HDAC repressor complex, have been identified in patients with the severe CHD hypoplastic left heart syndrome (X. Liu et al. 2017). Furthermore, in both mouse and zebrafish, loss of the *Sap130* gene causes heart defects (DeMoya et al. 2023; X. Liu et al. 2017). Together, this suggests that the HDAC repressor complex is broadly required for proper cardiac septation and valve development, and its haploinsufficiency or dysfunction may contribute to the spectrum of conotruncal and septal defects observed in human patients.

### Gene–environment interaction reshapes chromatin accessibility, metabolism, cell division, and trabeculation in the developing cardiac chambers

The functional cardiac defects observed in GxE may have arisen indirectly as a result of the structural heart defects, or may be a direct result of the GxE. To distinguish between these possibilities, we investigated changes in gene expression (Supplementary spreadsheet 6) and chromatin accessibility (Supplementary spreadsheet 7) in cardiomyocytes. In G×E condition *vs* Gene, among the top downregulated GO terms in cardiomyocytes, *generation of precursor metabolites and energy* ranked first, pointing to a profound alteration of cellular energy metabolism (Supplementary Table 7). Given the predominant reliance of embryonic cardiomyocytes on glycolysis (Stéphanie Ibrahim et al. 2023; Nakano et al. 2021), this finding strongly suggests an impairment of glycolytic pathways in the G×E condition (Supplementary Fig. 9, Supplementary Table 7).

*Chromatin remodeling* emerged as a top enriched term in all comparisons involving the G×E condition, but not in Environment *vs* Control or Gene *vs* Control. Key genes contributing to this term include *Hira*, *Hdac1*, *Suz12*, *Hdac2*, and *Ezh2* (Supplementary Fig. 9). This indicates that, while chromatin remodeling–related genes may still be affected by hyperglycemia, their contribution is not dominant, as they do not rank among the top ten enriched terms (Fig. 7A). These findings suggest that the G×E condition triggers a profound reorganization of chromatin accessibility, which is absent when genetic or environmental factors act alone. This result is consistent with the context-dependent function of NKX2-5, a transcription factor capable of acting as either an activator or a repressor in cardiomyocytes (Hiroi et al. 2001; Dupays et al. 2015; Nakashima et al. 2014; Robbe et al. 2022).

**Fig. 7.**
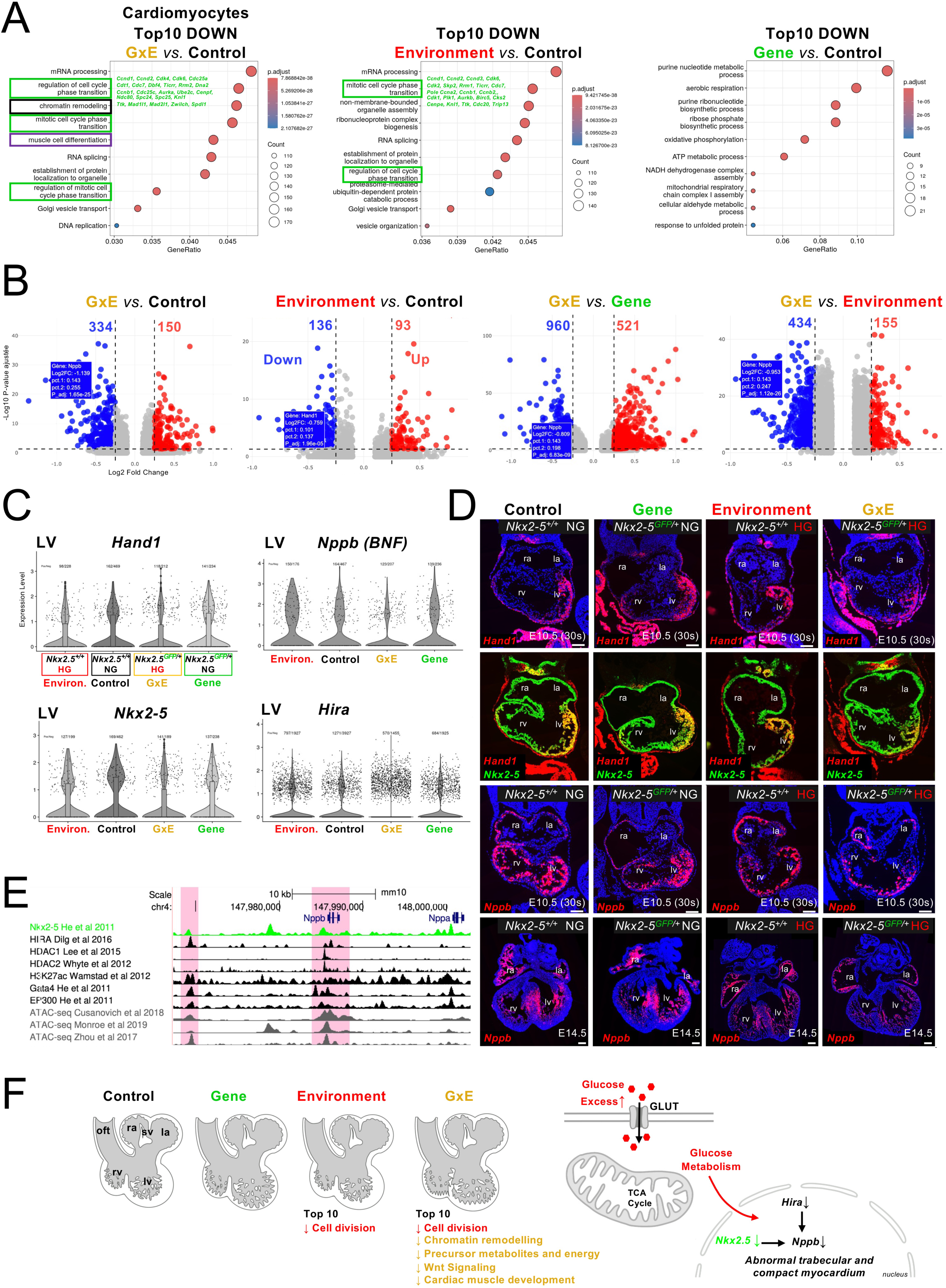
Chromatin-remodelling, cell-cycle, contractile, and trabecular gene programs are downregulated in G×E cardiomyocytes. (A) Differentially expressed genes were analyzed to identify significantly enriched GO terms and biological pathways, highlighting the biological processes and molecular functions associated with the observed transcriptional changes. (B) Volcano plots of differentially expressed genes within the cardiomyocyte cluster for each comparison, with downregulated (left, blue) and upregulated (right, red) genes (thresholds: |log2FC| ≥ 0.5, adjusted *P* < 0.05). Genes highlighted in blue boxes indicate *Hand1* and *Nppb*. (C) Violin plots of *Hand1, Nppb, Nkx2-5 and Hira* in the ventricular subcluster reveal significant perturbation in the G×E group (MAST, adjusted P < 0.05; y-axis: log-normalized counts). (D) Spatial validation by RNAScope of *Hand1* expression at E10.5 and *Nppb* at E10.5 and E14.5–E15.5 across the four experimental groups; scale bar = [µm]. (E) Analysis of existing ChIP-seq datasets indicates co-occupancy of Nkx2-5 and HIRA at the *Nppb* locus. Publicly available ChIP-seq data were analyzed within Cistrome Data Browser v2.0(Taing et al. 2024) using the UCSC Genome Browser(Perez et al. 2025) and genome annotations from the mm10 reference assembly. (F) Schematic representation summarizing the main enriched Gene Ontology (GO) terms identified in the cardiomyocyte cluster across the four experimental groups. Combined hyperglycemia and *Nkx2-5* haploinsufficiency downregulate *Hira* and subsequently the trabecular marker *Nppb*, resulting in defective trabeculation and myocardial compaction.

The term *mitotic cell phase transition* was significantly represented in all comparisons involving the G×E condition, as well as in the Environment *vs*. Control comparison, but not in Gene *vs*. Control, as expected. The coordinated downregulation of genes involved in cell-cycle entry, DNA replication, and mitosis is consistent with reduced proliferative activity or a lower proportion of cycling cells. This finding is consistent with the observed alterations in myocardial compact zone thickness under the Environment and G×E conditions (Fig. 7A, Table 1, Fig. 3).

In contrast to the other comparisons, in the G×E *vs*. Environment comparison the top 10 downregulated genes include those associated with WNT signaling, such as *Wnt11* and *Lef1*, as well as *Nkx2-5* itself (Supplementary Fig. 9). The decrease in *Nkx2-5* is expected given the difference in gene dosage between these groups. This is consistent with the notion that NKX2-5-dependent perturbation of the WNT signaling pathway promotes cardiac dysfunction through alterations in cardiomyocyte metabolism (Furtado et al. 2017). Interestingly, among the top 10 upregulated terms when comparing G×E *vs* Gene, the term *skeletal muscle development* emerges, which is consistent with the bulk RNA-seq results obtained at the foetal stage (Supplementary Fig. 9, Fig. 3), suggesting a persistence or reactivation of early myogenic programs. Among the top 10 upregulated genes when comparing Environment *vs* Control, the terms *mesenchymal development* and *heart morphogenesis* emerge, suggesting an environment-driven remodeling of developmental programs and cardiac differentiation pathways, potentially reflecting enhanced activation of mesenchymal and extracellular matrix–related processes associated with cardiac tissue remodeling (Supplementary Fig. 9).

In cardiomyocytes, the observation of a higher number of downregulated than upregulated genes in Environment *vs* Control suggests a global transcriptional repression in the environmental condition (Fig. 5B). Analysis of DEG in the ventricular cardiomyocytes showed significant perturbation of *Hand1*, and *Nppb* in the G×E group compared to the other 3 groups (Fig. 7C-D, Supplementary Spreadsheet 6). Spatial assays at E10.5 confirmed reduced *Hand1* and *Nppb* trabecular markers, consistent with the compaction defect observed at E14.5 (Fig. 7D).

Within the downregulated chromatin remodelers, we found *Hira* (Fig. 7C). NKX2-5 and HIRA co-occupy at a shared enhancer regulating *Tnni2-Tnnt3* expression (Dilg et al. 2016; Dupays et al. 2015)(Supplementary Fig. 3D). This suggests two non-exclusive possibilities: either the reduced NKX2-5 dosage under hyperglycemia prevents recruitment of the HIRA/NKX2-5 complex to this enhancer, leading to de-repression of skeletal-type troponins, consistent with the observations at E15.5; or the hyperglycemia-induced reduction of *Hira* itself allows ectopic expression of *Tnni2* and *Tnnt3*. Similar mechanisms involving NKX2-5, HIRA, and HDAC1/2 may operate at the *Nppb* locus (Fig. 7F-E). *Hand1* and *Nppb* are essential for ventricular trabeculation, altered expression of these genes can correlate with thin myocardial walls (Choquet et al. 2018a; Firulli et al. 2020), consistent with the increased incidence of dilated chambers we observed in the G×E group. Violin plots of *Nkx2-5* in the ventricular subcluster reveal significant perturbation in the G×E group (Fig. 7C; MAST, adjusted P < 0.05; y-axis: log-normalized counts) in the most severe G×E condition compared to the other groups. This reduction is consistent with the diabetes-induced loss of the compact myocardial zone and is further amplified by *Nkx2-5* haploinsufficiency, together accounting for the observed decrease in *Nkx2-5* expression (Fig. 7C). This also aligns with our previous findings combining gestational hypoxia and *Nkx2-5* haploinsufficiency (Moreau et al. 2019a) where both transcript and protein levels decrease in G×E conditions. In line with these findings, cell-type–specific estimated effect analyses revealed a reduced proportion of ventricular cardiomyocytes in both the G×E *vs* Control and Environment *vs* Control comparisons (Supplementary Fig. 5A-B).

### Maternal diabetes selectively disrupts a small subset of *Nkx2-5-*dependent regulatory elements controlling cardiomyocyte maturation

To investigate whether the transcriptional alterations observed in GxE cardiomyocytes were associated with changes in chromatin accessibility, we analyzed the snATAC-seq dataset generated from cardiomyocytes isolated from control and GxE embryos. Differential accessibility analysis identified 966 DARs between the two conditions (Fig. 8A-B). Functional enrichment analysis revealed that these DARs were highly enriched for biological processes governing cardiac muscle development, cardiomyocyte differentiation, sarcomere organization, myofibril assembly and cardiac contraction (Fig. 8A, Supplementary Spreadsheet7, Supplementary Table 8). This suggests that maternal diabetes primarily remodels regulatory elements involved in cardiomyocyte maturation rather than general cellular functions. Because *Nkx2-5* is the sensitizing genetic factor in our model, we next asked which accessibility changes might represent direct NKX2-5-dependent regulatory events. Motif analysis showed that only 228 of the 966 DARs (24%) contained a canonical NKX2-5 binding motif (Fig. 8B). We integrated these data with the matched single-nucleus RNA-seq dataset to identify chromatin changes associated with transcriptional output. Remarkably, only five NKX2-5 motif-containing DARs were associated with dysregulated genes, representing 0.5% of all DARs (Fig. 8B). Among the less accessible DARs containing an NKX2-5 consensus binding motif and associated with differentially expressed genes were regions linked to *Myl2*, *Mybpc3*, *Myo18b*, and *Actn2*, which all encode key structural and regulatory components of the cardiac sarcomere. This striking reduction from hundreds of chromatin accessibility changes to only 5 transcriptionally responsive loci suggests that most chromatin-remodelling events induced by maternal diabetes are unlikely to reflect direct NKX2-5-mediated transcriptional regulation. However, it is formally possible that NKX2-5 may be recruited to regulatory regions indirectly through interactions with DNA-bound cofactors. The affected regulatory program appears to converge on a restricted network controlling cardiomyocyte contractile maturation. Consistent with this interpretation, motif enrichment analysis of the 228 DARs containing an NKX2-5 motif identified significant enrichment for GATA4 and SMAD4 binding motifs. This finding is consistent with previous studies demonstrating functional cooperation between NKX2-5 and other cardiac transcription factors. In particular GATA4 is known to physically interact with NKX2-5 and facilitate its transcriptional activity and DNA binding at cardiac regulatory elements (Sepulveda et al. 1998; Durocher et al. 1997). Furthermore, genome-wide studies have shown extensive co-occupancy of NKX2-5 and GATA4 at cardiac regulatory regions, supporting a combinatorial mode of transcriptional regulation (He et al. 2011; Schlesinger et al. 2011). Thus, the enrichment of GATA4 and SMAD4 motifs suggests that these factors may contribute to the recruitment or stabilization of NKX2-5 at a subset of differentially accessible regulatory regions.

**Fig. 8.**
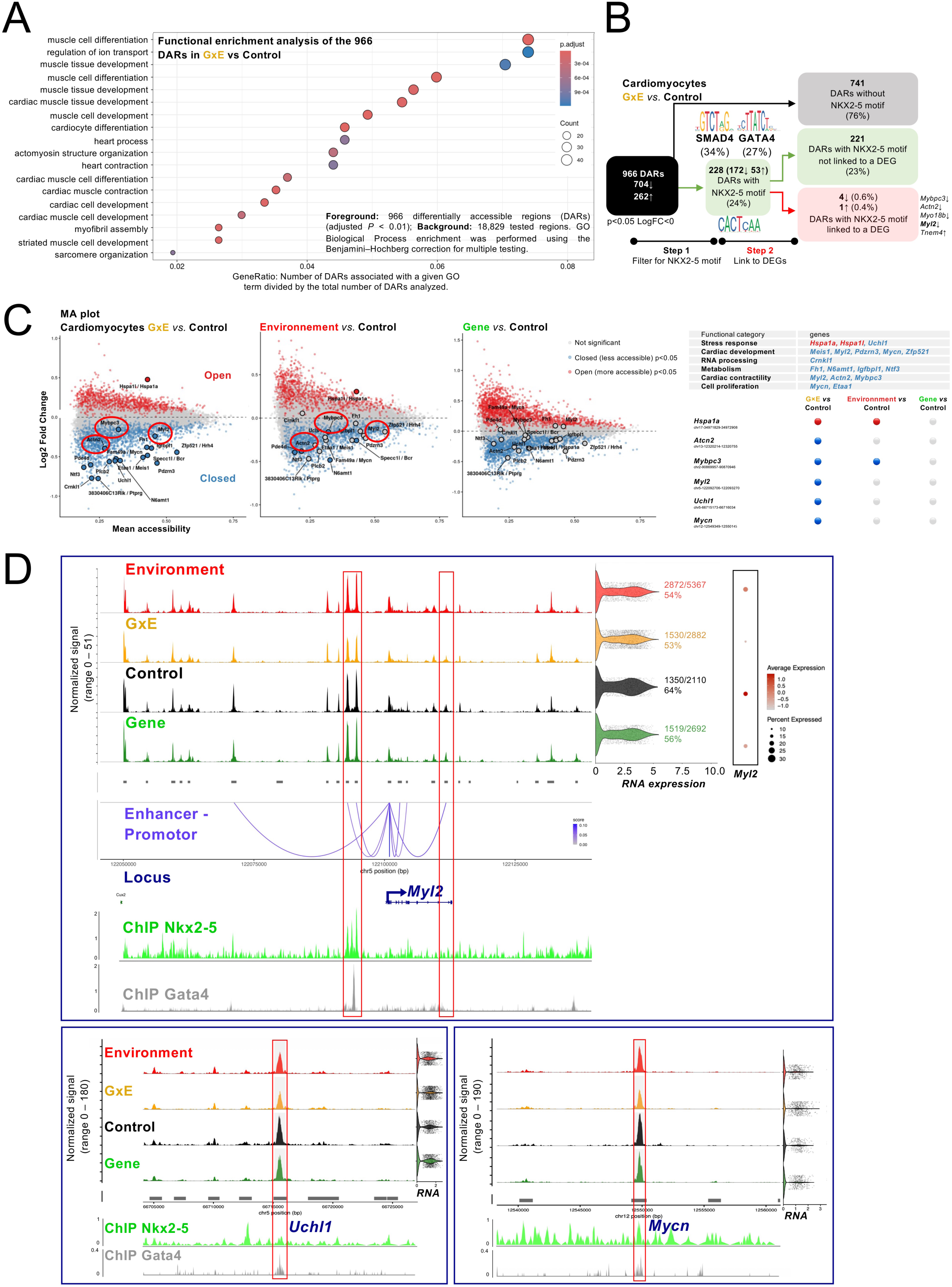
Multi-omic integration identifies a restricted Nkx2-5-dependent regulatory network disrupted by maternal diabetes. (A) Gene Ontology (GO) enrichment analysis of the 966 differentially accessible regions (DARs) in GxE cardiomyocytes compared with controls. Enriched biological processes are predominantly related to cardiac and striated muscle development, sarcomere organization, muscle differentiation, and heart contraction (B) Strategy used to prioritize candidate direct Nkx2-5 regulatory targets. Among the 966 DARs, 228 contained an Nkx2-5 consensus binding motif. Integration with differentially expressed genes identified only five candidate loci (*Actn2*, *Mybpc3, Myo18b, Tnem4* and *Myl2*) displaying coordinated changes in chromatin accessibility and gene expression. Motif enrichment analysis further identified significant enrichment for NKX2-5, SMAD4 and GATA4 binding sites within these regions. (C) MA plots showing differential chromatin accessibility in the G×E *vs.* Control, Environment *vs.* Control, and Gene *vs.* Control comparisons. Regions with increased accessibility are shown in red, whereas regions with decreased accessibility are shown in blue (adjusted *P* < 0.05). Differential accessibility is most pronounced in the GxE versus Control comparison. Functional annotation highlights representative genes involved in stress response, cardiac development, metabolism, RNA processing, cardiac contractility and cell proliferation. Comparison of chromatin accessibility changes across experimental groups identifies *Hspa1a*, *Actn2*, *Mybpc3* and *Myl2* as representative loci displaying condition-specific alterations. (D) Genome browser views of the *Myl2*, *Uchl1*, and *Mycn* loci showing chromatin accessibility profiles across the four experimental groups together with paired single-cell RNA expression. Genome browser views showing chromatin accessibility profiles, predicted enhancer–promoter interactions, and previously published NKX2-5 and GATA4 ChIP-seq datasets. Candidate regulatory regions highlighted in the analysis are outlined by red boxes.

To further visualize chromatin accessibility changes across cardiomyocytes, MA plots were generated for the GxE *vs*. Control, Environment *vs*. Control and Gene *vs*. Control comparisons (Fig. 8C). DARs (adjusted *P* < 0.05) are highlighted according to increased (red) or decreased (blue) accessibility. The GxE *vs*. Control confirmed the reduced chromatin accessibility of *Mybpc3*, *Actn2* and *Myl2*, as well as highlighting additional loci also involved in cardiac contractility, such as *Uchl1* and *Mycn*. By contrast, the Environment *vs*. Control comparison showed fewer decreases in accessibility at these loci, and in the Gene *vs*. Control comparison most of these regions were not significantly altered at all. Conversely, the *Hspa1a* locus displayed increased chromatin accessibility in both the GxE and Environment conditions (p< 0.05), but not in the Gene condition. Finally, genome visualization revealed reduced chromatin accessibility in the G×E condition compared with the Control, Environment, and Gene conditions, specifically within predicted enhancer regions overlapping previously published NKX2-5 ChIP-seq peaks and enhancer–promoter interactions (Fig. 8D). Among these candidate loci, the *Myl2* locus provided particularly strong evidence for direct regulatory disruption. A region showing reduced chromatin accessibility in G×E embryos overlaps a previously reported NKX2-5 ChIP-seq peak in fetal hearts, indicating that NKX2-5 binds this regulatory region during cardiac development. Together with the reduced accessibility observed at the *Myl2* locus in G×E embryos and the established role of *Myl2* in ventricular chamber identity, these findings support the possibility that altered NKX2-5-dependent regulation of *Myl2* may contribute to the impaired contractile program observed in the G×E condition. These chromatin changes were accompanied by decreased *Myl2* expression in cardiomyocytes, supporting a model in which maternal diabetes impairs activation of NKX2-5-dependent regulatory elements controlling sarcomeric gene expression. Together, these findings indicate that maternal diabetes induces widespread chromatin remodeling in cardiomyocytes, yet only a highly restricted subset of regulatory regions containing consensus NKX2-5 binding sites is associated with measurable transcriptional changes. This selective vulnerability identifies sarcomeric maturation genes as primary downstream effectors of the interaction between maternal hyperglycemia and *Nkx2-5* haploinsufficiency. Genome browser views identified additional candidate regulatory elements associated with *Uchl1* (involved in stress response (P. Wu et al. 2022)), and *Mycn* (involved in cell proliferation (Muñoz-Martín et al. 2019)) (Fig. 8D). In both loci, chromatin accessibility was selectively reduced in GxE cardiomyocytes compared with the other experimental groups. These DARs overlapped published SMAD4 ChIP-seq peaks, but did not coincide with NKX2-5 binding site ChIP-seq peaks, sugesting that they are unlikely to represent direct NKX2-5 targets. The *Mycn* locus is of particular interest given the established role of MYCN as a downstream effector of BMP/SMAD4 signaling that promotes ventricular cardiomyocyte proliferation, ventricular wall morphogenesis and trabeculation during embryonic heart development (Harmelink et al. 2013).

## Discussion

Consistent with our observations, other environmental modifiers have been shown to critically influence the penetrance and severity of cardiac defects in genetically-predisposed contexts. In mouse, maternal age has been significantly correlated with an increased risk of both membranous and muscular VSDs in *Nkx2-5*^+/−^ offspring (Winston et al. 2012; Schulkey et al. 2015b, 2015c). Furthermore, previously we have shown massive embryonic lethality in a model combining *Nkx2-5* haploinsufficiency with severe gestational hypoxia (5.5% O₂). In this model approximately half of the *Nkx2-5*^+/−^ embryos exhibited dilated cardiac chambers at E10.5 prior to death (Moreau et al. 2019b). By contrast, our new data indicate a later onset of lethality, occurring at foetal stages, suggesting that different environmental stressors differentially modulate both the timing and nature of cardiac pathology.

*Nkx2-5^GFP/+^* foetuses exposed to maternal hyperglycemia show a higher penetrance of hypertrabeculated ventricular walls, accompanied by an increased incidence of ventricular septal defects, including muscular VSDs, compared to controls. Ventricular noncompaction is a cardiomyopathy characterized by the persistence of the trabecular layer of the ventricular walls, leading to both diastolic and systolic dysfunction in some patients. Under normal conditions, the ventricular trabecular layer develops around murine E10.5, coinciding with a rise in myocardial mass prior to the formation of a distinct coronary circulation (Goodyer and Wu 2018). As cardiac development progresses, the compact myocardium thickens while the trabecular myocardium becomes less prominent. Although the pathogenesis of ventricular noncompaction remains poorly understood, it is thought to result from an early arrest in myocardial maturation, marked by a failure in the compaction and cessation of trabecular proliferation (Goodyer and Wu 2018). *In Nkx2-5^GFP/+^* foetuses exposed to hyperglycemia, the ventricular septum appears irregular in shape, or displays multiple muscular defects. Since the ventricular septum itself forms through compaction of the trabecular myocardium (Sedmera et al. 2000), this supports a mechanistic link between persistent trabecular layers and the observed muscular abnormalities. Proper development of the ventricular septum requires coordinated interactions between the myocardium and the epicardium, including contributions from epicardium-derived cells (Gittenberger-de Groot et al. 2014; Poelmann et al. 2016, 2014). The defects observed under the gene-environment (G×E) condition are unlikely to result from epicardial dysfunction, as *Nkx2-5* is not expressed in epicardial cells, nor are these cells derived from *Nkx2-5*-positive progenitors.

Our observation of a reduced expression of *Aldh1a2* in GxE embryos at E9.5 might be expected to lead to impaired RA signaling, and this may contribute to the ventricular trabeculation defects detected at later stages. Indeed, RA deficiency in *Raldh2-*null embryos results in severely abnormal ventricular trabeculation, with the developing myocardium forming a disorganized layer rather than normal trabeculae (Captur et al. 2016; Niederreither et al. 2001; Ryckebusch et al. 2008). Because early ventricular trabeculae contribute to the formation of the muscular interventricular septum, disruption of this process could also contribute to the muscular ventricular septal defects observed in our model. However, a direct causal relationship remains to be established. The anomalies observed under G×E conditions are primarily concentrated in the ventricles and the myocardial wall, without a significant increase in conotruncal malformations, suggesting that the contribution of SHF progenitors is largely preserved. Nevertheless, recent studies show that cells expressing the SHF program have been identified in the interventricular septal region using an intersectional lineage reporter approach (Devine et al. 2014). Studies employing Cre recombinase under the control of an aSHF enhancer (*Mef2c-AHF-Cre*) have demonstrated that TBX5 function within the SHF is essential for the development of the muscular portion of the ventricular septum (Koshiba-Takeuchi et al. 2009). RA signaling plays an upstream role in establishing a *Tbx5*-positive pSHF progenitor population (De Bono et al. 2018). A transient population of cells coexpressing *Tbx1* (aSHF) and *Tbx5* (pSHF) is specifically observed in the posterior dorsal pericardial wall (DPW), corresponding to an overlapping region between aSHF and pSHF progenitors. This coexpression defines an intermediate stage in the progressive segregation of progenitor cells fated for the arterial and venous poles. Retinoic acid, and particularly the enzyme RALDH2 that catalyzes its synthesis, marks this overlapping region and acts upstream to regulate *Tbx5* expression within this territory. *Nkx2-5* expression overlaps with *Tbx1* in the aSHF, and likely also in the transitional zone where pSHF markers *Tbx5* and *Aldh1a2* (encoding RALDH2) are coexpressed. Our data indicate that maternal hyperglycemia, in combination with *Nkx2-5* haploinsufficiency, affect the organization of the SHF, potentially by altering the identity of this transient overlapping region between the aSHF and pSHF, and could contribute to impaired formation of the muscular ventricular septum.

The ectopic expression of the fast skeletal troponin isoforms *Tnni2* and *Tnnt3* in the embryonic heart under conditions of *Nkx2-5* haploinsufficiency and maternal hyperglycemia may indicate a convergence between genetic and environmental stressors affecting cardiac gene expression. *Tnni2* and *Tnnt3* encode fast skeletal muscle–specific isoforms of troponin I and T, respectively, and are normally restricted to skeletal muscle, although *Tnni2* is transiently expressed in embryonic hearts between E9.5 and E16.5 (L. Zhu et al. 1995). Their ectopic activation in cardiac tissue under stress or pathological conditions indicates that, while they may not be essential for normal cardiac development, their misexpression could interfere with cardiac function. Notably, increased expression of *Tnnt3* has been linked to impaired cardiac contractile function (Q.-Q. Huang et al. 2008).

Although often assumed to be a transcriptional activator, NKX2-5 has also been shown to function as a transcriptional repressor (Prall et al., 2007). In this case, its role appears critical in maintaining proper muscle fiber-type identity. The marked upregulation of *Tnnt3* and *Tnni2* observed in the developing atria of *Nkx2-5^GFP/+^* embryos exposed to hyperglycemia closely resembles the expression pattern seen in the *Nkx2-5^GFP/IRES-cre^*hypomorphic model. Therefore, both *Nkx2-5^GFP/+^* hearts exposed to hyperglycemia and *Nkx2-5^GFP/IRES-cre^* are likely to exhibit impaired contractile performance, potentially driven by altered myofibrillar calcium sensitivity and excitation-contraction coupling. The morphological abnormalities observed at E14.5 in both *Nkx2-5^GFP/IRES-cre^* hearts and *Nkx2-5^GFP/+^* hearts exposed to hyperglycemia include wall thinning and atrial distension, features indicative of impaired cardiac function.

At the molecular level, ChIP-seq analyses reveal strong enrichment of both NKX2-5 and HIRA, a conserved histone chaperone, at the shared enhancer of the troponin genes *Tnni2* and *Tnnt3* (referred to as the TTe site) in E12.5 wild-type hearts (Dilg et al. 2016). Previous ChIP experiments in embryonic hearts (Dupays et al. 2015), as well as DamID assays in HL-1 cardiomyocyte-like cells (Bouveret et al. 2015), have demonstrated direct binding of NKX2-5 to the TTe site. These findings support the notion that NKX2-5 contributes to the regulation of muscle fiber-type identity by repressing the expression of the fast skeletal troponin isoforms *Tnni2* and *Tnnt3*, thereby preventing inappropriate activation of a fast skeletal muscle gene program in the heart (Dupays et al. 2015). This may result from a direct effect of reduced NKX2-5 levels on atrial myofibrillar composition. Interestingly, a similar upregulation of fast skeletal troponin isoforms has been reported in several models of cardiac dysfunction triggered by the loss of either the Polycomb Repressive Complex 2 (PRC2) (Delgado-Olguín et al. 2012) or the histone deacetylases HDAC1 and HDAC2 (Montgomery et al. 2007; He et al. 2011). Glucose metabolism plays a pivotal role in regulating histone acetylation and guiding myogenic lineage commitment (Yucel et al. 2019). Therefore, misexpression of fast skeletal troponin isoforms could also be mediated by glucose-induced epigenetic deregulation. Further supporting this HIRA is enriched at the TTe locus in wild-type hearts, and its conditional deletion in the cardiogenic lineage leads to increased expression of the fast skeletal troponin genes *Tnni2* and *Tnnt3* (Dilg et al. 2016). In a separate model with *Notch1* gain of function, elevated *Tnni2* expression was associated with myocardial structural abnormalities (Watanabe et al. 2006). Transcriptomic analysis following HIRA inactivation revealed upregulation of *Tnni2* and *Tnnt3*, with concurrent downregulation of *Tnnt1* (Stone et al. 2007; Dilg et al. 2016).

Together, these findings suggest that maternal hyperglycemia and *Nkx2-5* haploinsufficiency do not simply induce ectopic activation of a fast skeletal muscle program, but more broadly disrupt the maturation of the embryonic cardiac contractile apparatus. While *Tnni2* and *Tnnt3* are aberrantly activated, the concomitant reduction in *Myl2* expression points to an incomplete acquisition of the ventricular contractile phenotype. *Myl2* encodes the ventricular regulatory myosin light chain, a critical component of the sarcomere required for force generation and contractile maturation (W. Huang et al. 2015; Yuan et al. 2018). Thus, the simultaneous gain of skeletal muscle–specific troponin isoforms and loss of a key ventricular sarcomeric component support the concept of a developmental shift toward an immature and functionally compromised contractile state. Consistent with our findings, NKX2-5-null human cardiomyocytes exhibit reduced MYL2 expression together with decreased expression of other ventricular markers, including IRX4 and HAND1, leading to impaired progression toward a specialized ventricular phenotype (Anderson et al. 2018). These data suggest that maternal hyperglycemia further destabilizes an NKX2-5-dependent transcriptional program required for ventricular maturation, resulting in the simultaneous loss of ventricular contractile identity (*Myl2*) and inappropriate activation of fast skeletal muscle genes (*Tnni2* and *Tnnt3*). During embryogenesis, the cardiac transcription factors NKX2-5 and GATA4 are mutual cofactors which cooperate to activate cardiac structural genes and promote ventricular differentiation (Stefanovic and Christoffels 2015; Stefanovic et al. 2014; Durocher et al. 1997). Previous studies demonstrated that NKX2-5 physically interacts with GATA4 to synergistically activate cardiac gene expression (Durocher et al. 1997), while BMP/SMAD signaling cooperates with GATA4 to regulate cardiac transcriptional programs (Moskowitz et al. 2011; Brown et al. 2004). More recently, SMAD4 has also been shown to interact directly with NKX2-5 (Hu et al. 2021), suggesting that these factors function within an integrated transcriptional network controlling cardiomyocyte differentiation and maturation.

Our scMultiome analyses will provide an essential resource for linking non-coding variants identified by GWAS to the regulatory networks of cardiac development. By integrating epigenomic and transcriptomic dimensions, they will allow exploration of how genetic susceptibility interacts with an adverse maternal environment, such as pregestational diabetes, thereby helping to elucidate the mechanisms underlying CHD and guiding the development of new prevention strategies.

## Supplementary data legends

**Supplementary Table 1.** Summary of types of external defects observed in E15.5-E16.6 foetuses. Foetuses were assessed for gross developmental abnormalities and categorized according to maternal blood glucose levels: control (C), moderate (M), high (H), severe (S), severe reversed (SR), or control reversed (CR). SR represents severe hyperglycemia reversed at E3.5 by glibenclamide pellet implantation, while CR represents control females receiving a glibenclamide pellet at E3.5. Statistical significance was determined using a one-sided Fisher’s exact test.

| Types of defects | NG | Moderate | High | Severe | Severe reversed | Control reversed |
| --- | --- | --- | --- | --- | --- | --- |
| Exencephaly | 0 | 0 | 5 | 9 | 0 | 0 |
| Eye | 0 | 0 | 14 | 10 | 1 | 0 |
| Micrognathia | 0 | 4 | 10 | 8 | 0 | 0 |
| Lymphatic | 0 | 5 | 14 | 10 | 4 | 1 |
| Total abnormal | 0 | 6 | 19 | 13 | 4 | 1 |
| Total dead | 0 | 0 | 4 | 11 | 1 | 0 |
| Total normal | 76 | 21 | 25 | 9 | 13 | 21 |
| P value vs control | - | 0.0002 | <0.0001 | <0.0001 | 0.0002 | 0.2245 |
| P value vs Severe | <0.0001 | <0.0001 | 0.0102 | - | 0.0072 | <0.0001 |

**Supplementary Table 2.** Summary of types of heart defects observed under hyperglycemic conditions in the ßV59M model. Cardiac morphology was assessed on serial sections stained with haematoxylin and eosin. ASD, atrial septal defect; mVSD: membranous ventricular septal defect; musVSD: muscular ventricular septal defect; AVSD, atrioventricular septal defect; OFT, outflow tract defects (includes overriding aorta and double-outlet right ventricle). † one with transposition of the great arteries. Statistical significance was tested using two-sided Fisher’s exact test.

Ibrahim, Maurizio *et al.*, Supp. Table 2 Summary of types of heart defects observed under hyperglycemic conditions in the βV59M model.
| Types of defects |  | NG | Moderate | High | Severe | Severe reversed | Control reversed |
| --- | --- | --- | --- | --- | --- | --- | --- |
| Single | ASD | 0 | 0 | 0 | 0 | 0 | 0 |
|  | mVSD | 0 | 0 | 5 | 2 | 3 | 0 |
|  | muscVSD | 0 | 2 | 5 | 1 | 0 | 0 |
| Multiple | OFT + mVSD | 0 | 1 | 9 | 2 | 0 | 0 |
|  | OFT + mVSD + muscVSD | 0 | 1 | 3 | 0 | 0 | 0 |
|  | OFT + muscVSD | 0 | 0 | 0 | 0 | 0 | 0 |
|  | mVSD + ASD | 0 | 0 | 1 | 2 | 0 | 0 |
|  | OFT + AVSD | 0 | 0 | 0 | 6† | 0 | 0 |
|  | *situs inversus* + ASD + mVSD | 0 | 0 | 1 | 0 | 0 | 0 |
|  | *situs inversus* + ASD + mVSD + OFT | 0 | 0 | 1 | 0 | 0 | 0 |
|  | *situs inversus* | 0 | 0 | 1 | 0 | 0 | 0 |
|  | *ectopia cordis* | 0 | 0 | 0 | 1 | 0 | 0 |
| Total abnormal |  | 0 | 4 | 26 | 14 | 3 | 0 |
| Total normal |  | 44 | 21 | 6 | 7 | 11 | 2 |
| P value vs control |  | - | 0.0146 | <0.0001 | <0.0001 | 0.0118 | >0.9999 |
| P value vs Severe |  | <0.0001 | 0.0007 | 0.3290 | - | 0.0153 | 0.1423 |

**Supplementary Table 3.**
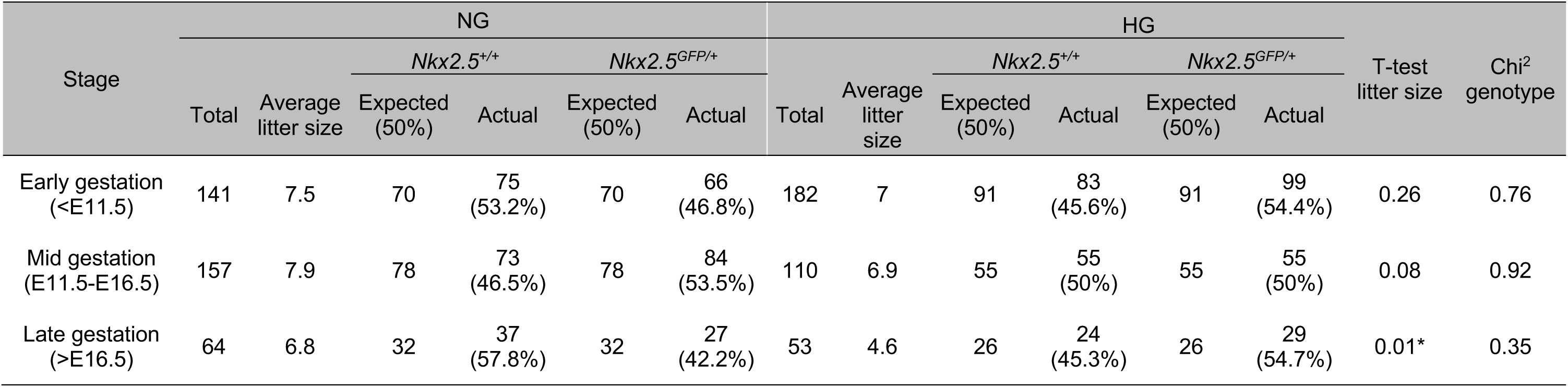
Genotype-dependent effects of maternal glycaemic status on litter size and embryonic viability across gestation. Average litter size in pregnancies derived from *Nkx2-5^GFP/+^*males was compared between normoglycemic (NG) and hyperglycemic (HG) conditions at three gestational stages: early, mid, and late gestation. Within these *Nkx2-5^GFP/+^* paternal crosses, no difference was detected at early or mid gestation, whereas average litter size was significantly reduced under HG conditions at late gestation compared to NG. To assess whether reduced litter size reflected genotype-specific loss, we tested Mendelian transmission of *Nkx2-5^GFP/+^ vs Nkx2-5^+/+^* in offspring and found no deviation from the expected 1:1 ratio in NG or HG cohorts. Thus, the late-gestation reduction in litter size is attributed to hyperglycemia rather than selective mortality of a specific *Nkx2-5* genotype. T-test was performed for litter size; and Chi^2^ for genotype ratios.

**Supplementary Table 4.**
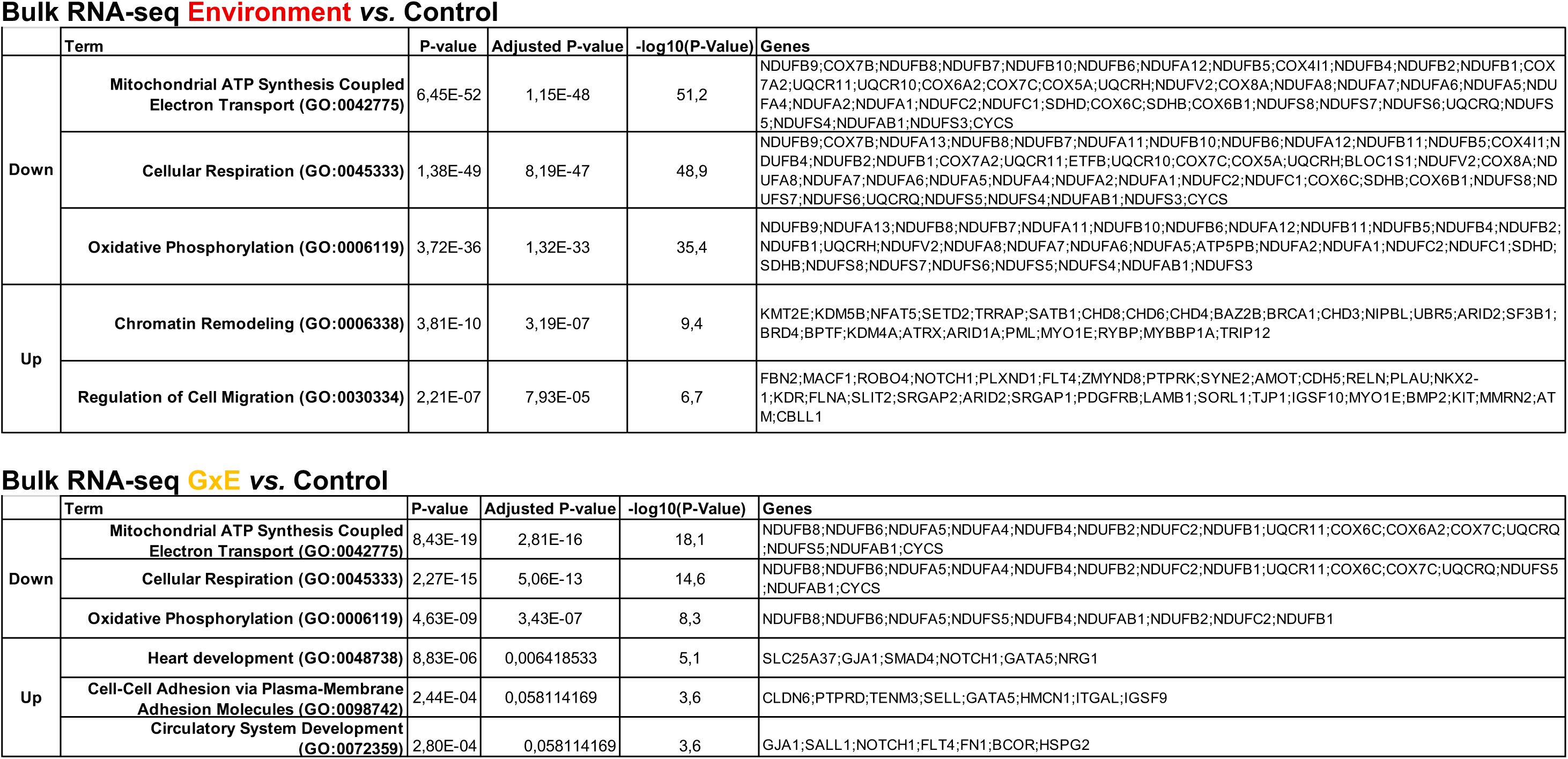
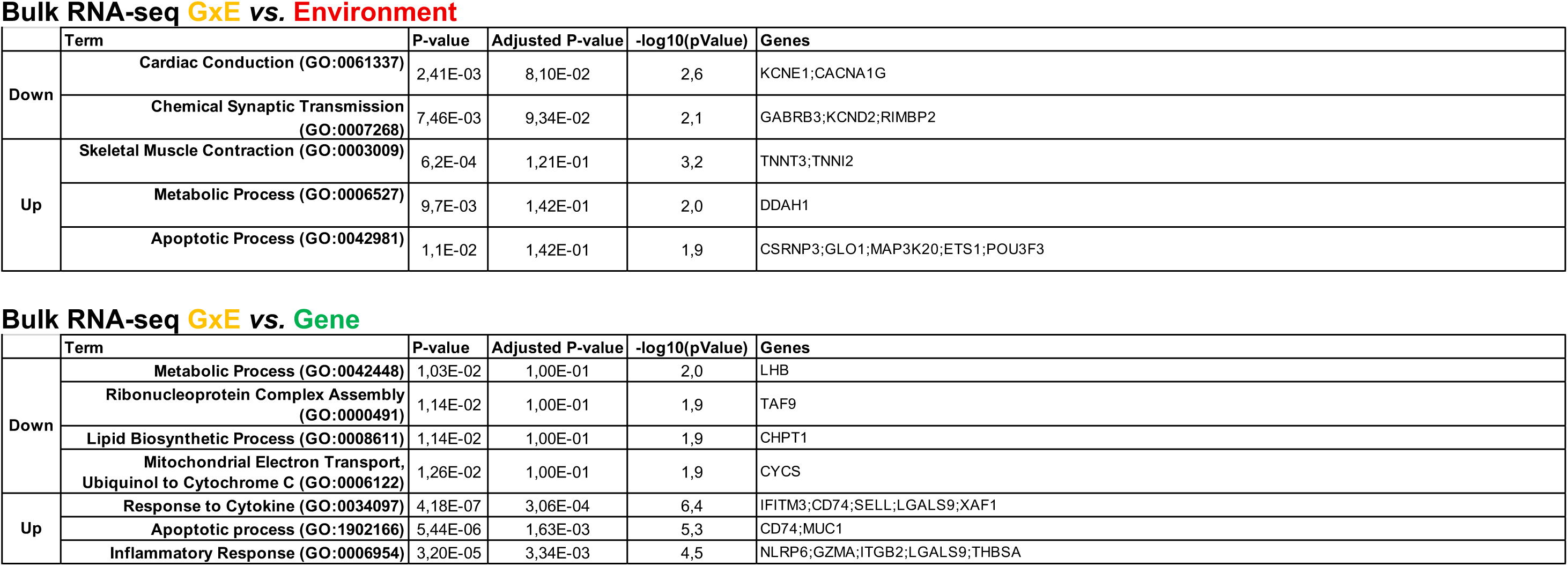
GO enrichment analysis of differentially expressed genes across four comparisons in bulk RNA-seq data from fetal hearts. GO terms were identified using enrichR, and the top 10 enriched terms for each comparison were ranked by adjusted p-value. For clarity, a subset of biologically relevant terms is shown. Associated genes contributing to each GO term are listed, highlighting pathways and processes potentially affected.

**Supplementary Table 5.**
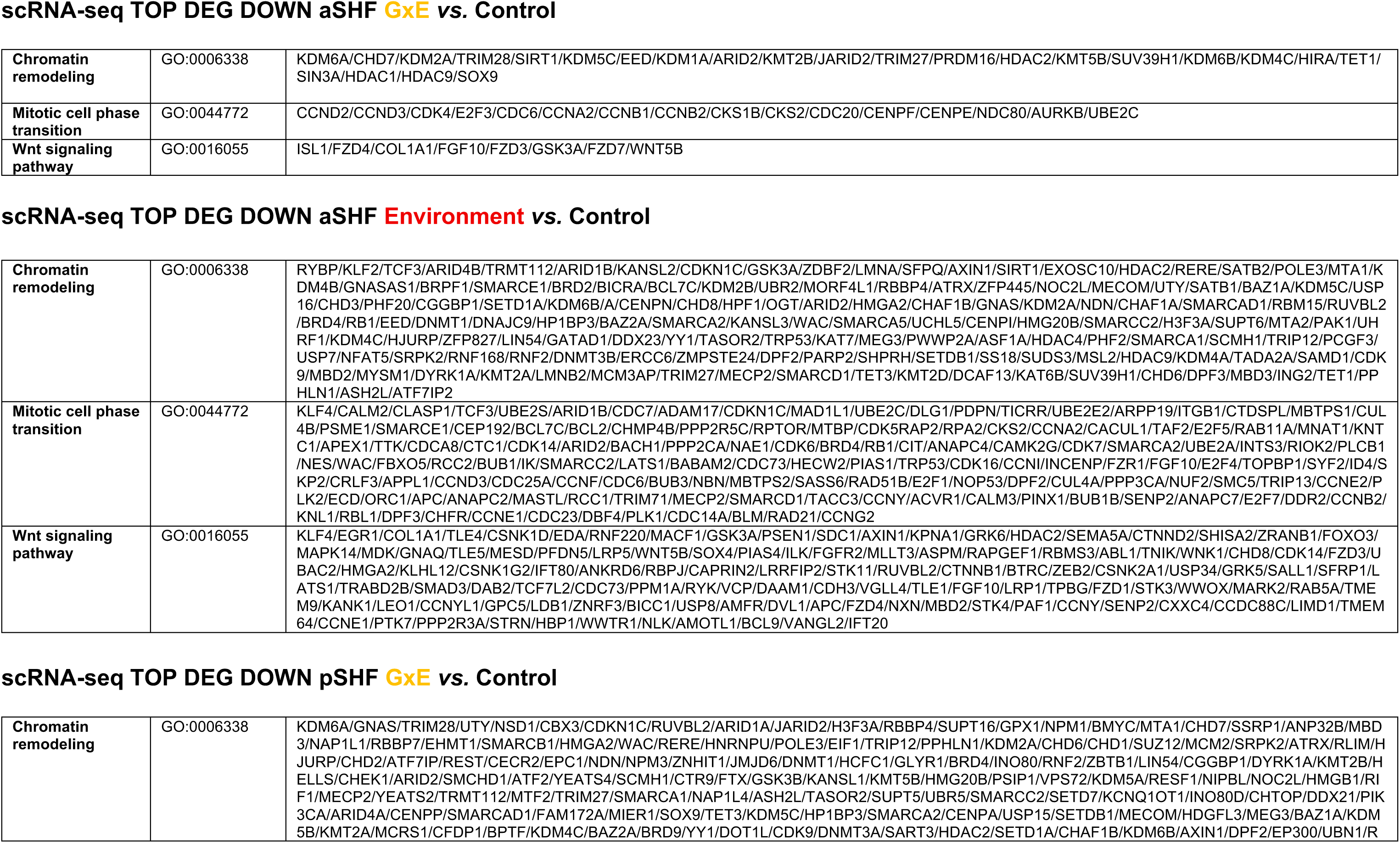
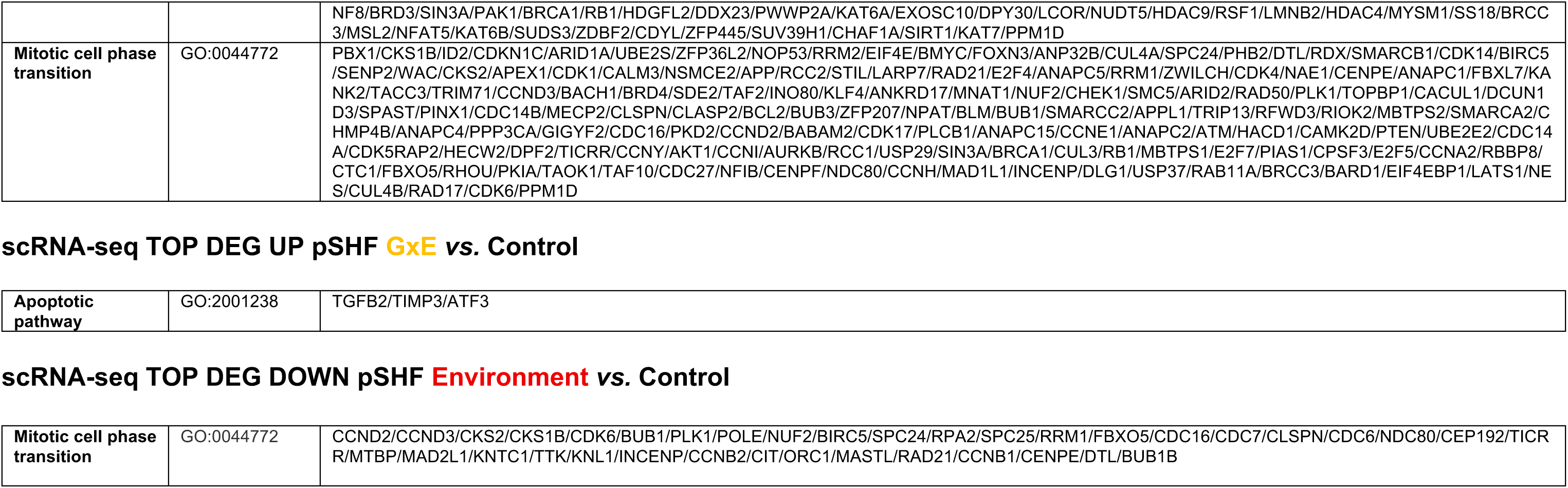
Selected GO terms associated with differentially expressed genes in second heart field progenitors. Genes associated with selected functional categories identified by scRNA-seq analysis of aSHF and pSHF progenitors in the G×E versus Control and Environment versus Control comparisons. Categories include chromatin remodeling, mitotic cell cycle phase transition, Wnt signaling, and apoptosis. To explore biological processes represented in an extended aSHF/pSHF gene set, GO over-representation analysis was performed using both downregulated genes and expressed genes that did not meet the differential expression significance threshold, with the full mouse genome as the background universe.

**Supplementary Table 6.**
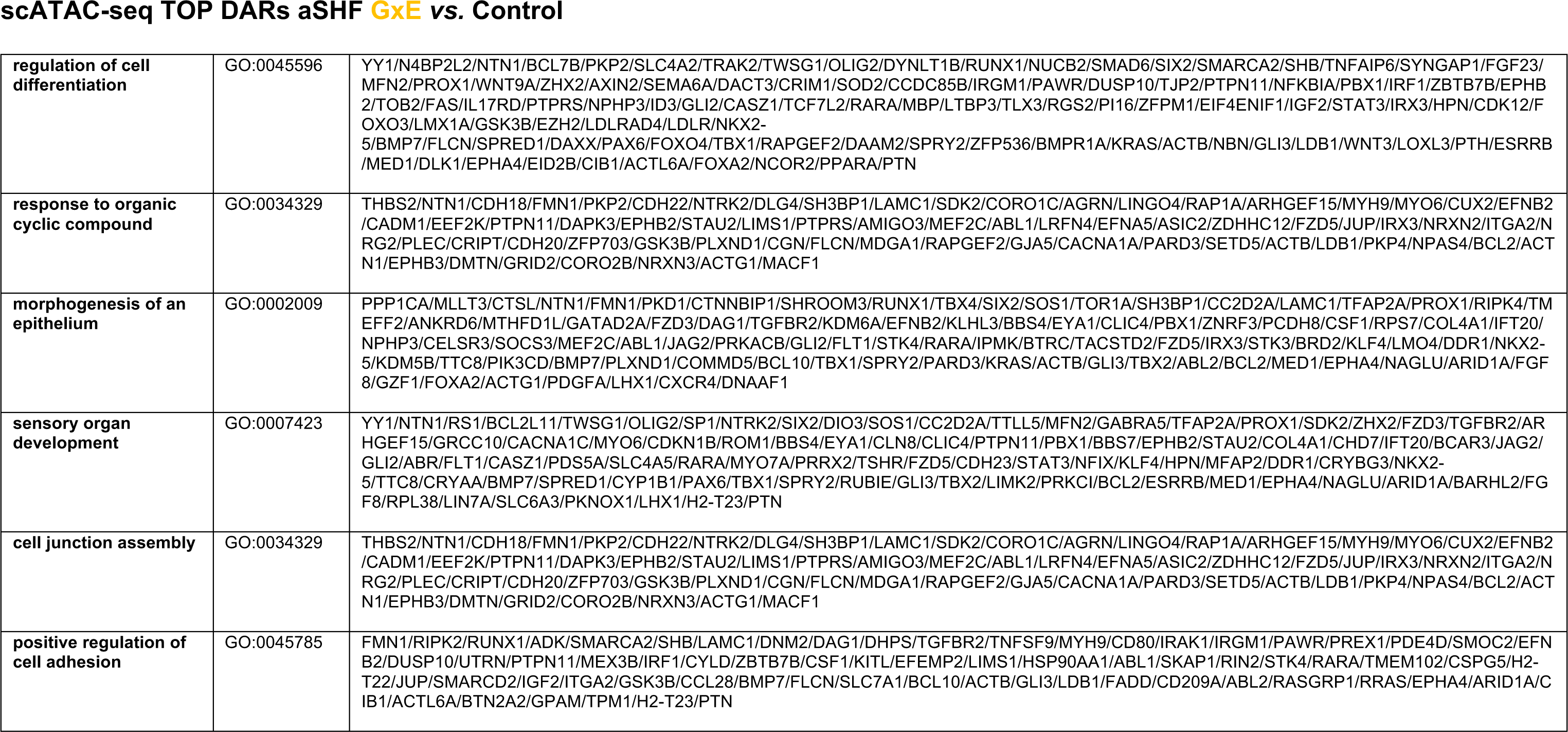
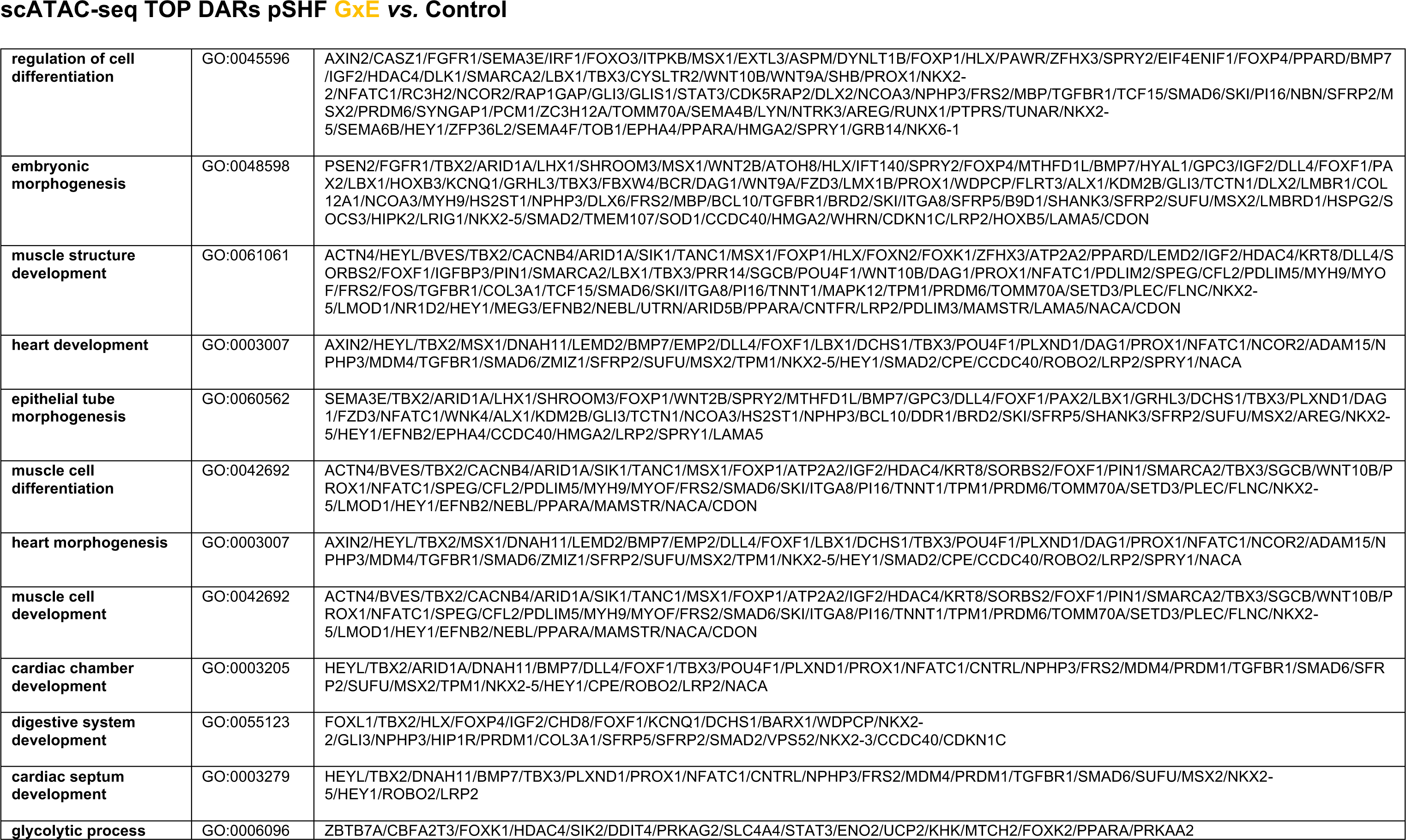
Selected GO terms associated with differentially accessible chromatin regions in second heart field progenitors. Selected GO terms and associated genes identified by functional enrichment analysis of DARs in aSHF and pSHF progenitors in the G×E versus Control comparison. GO term descriptions, identifiers, and genes associated with DARs are listed for each progenitor population.

**Supplementary Table 7.**
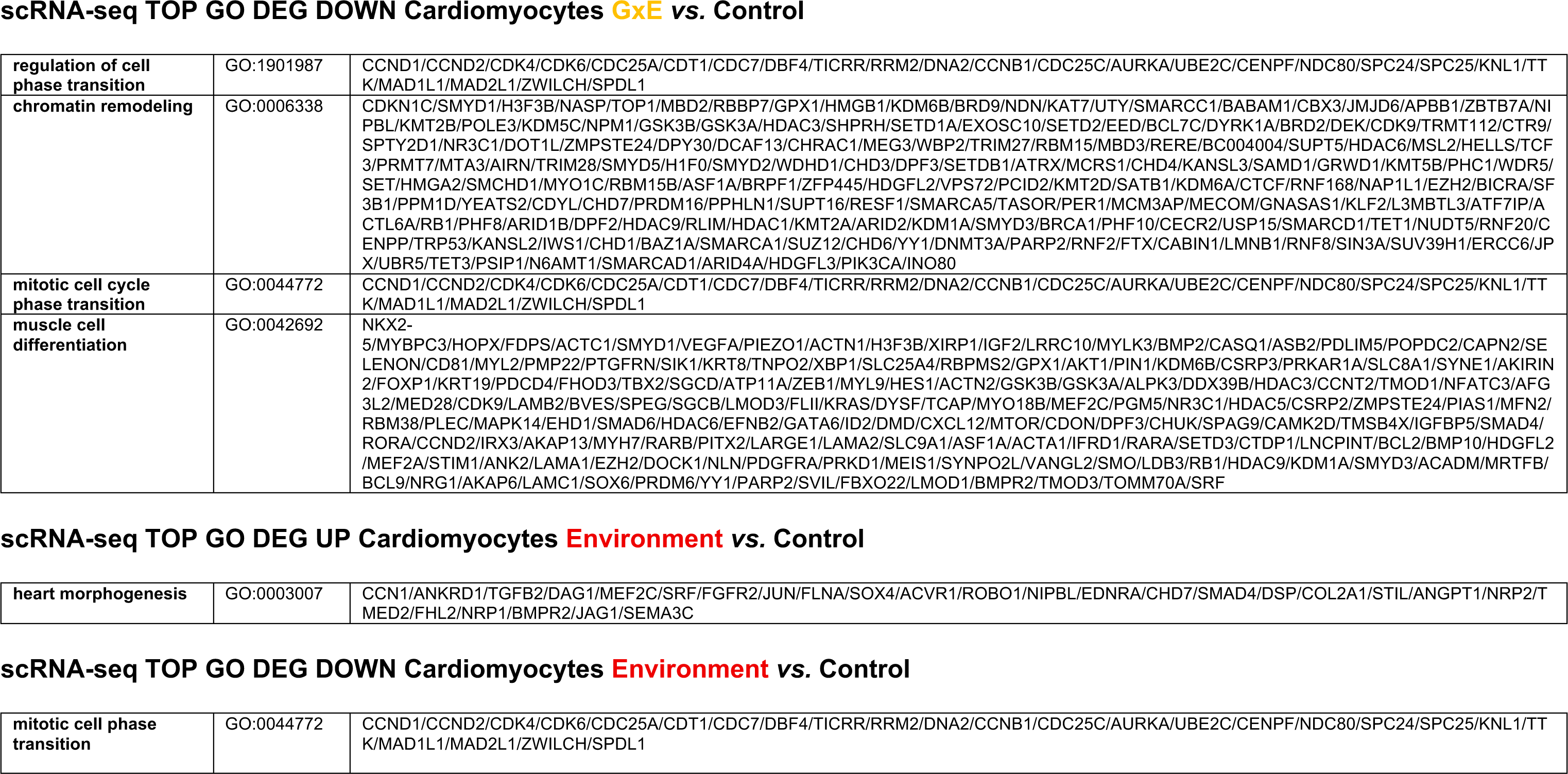
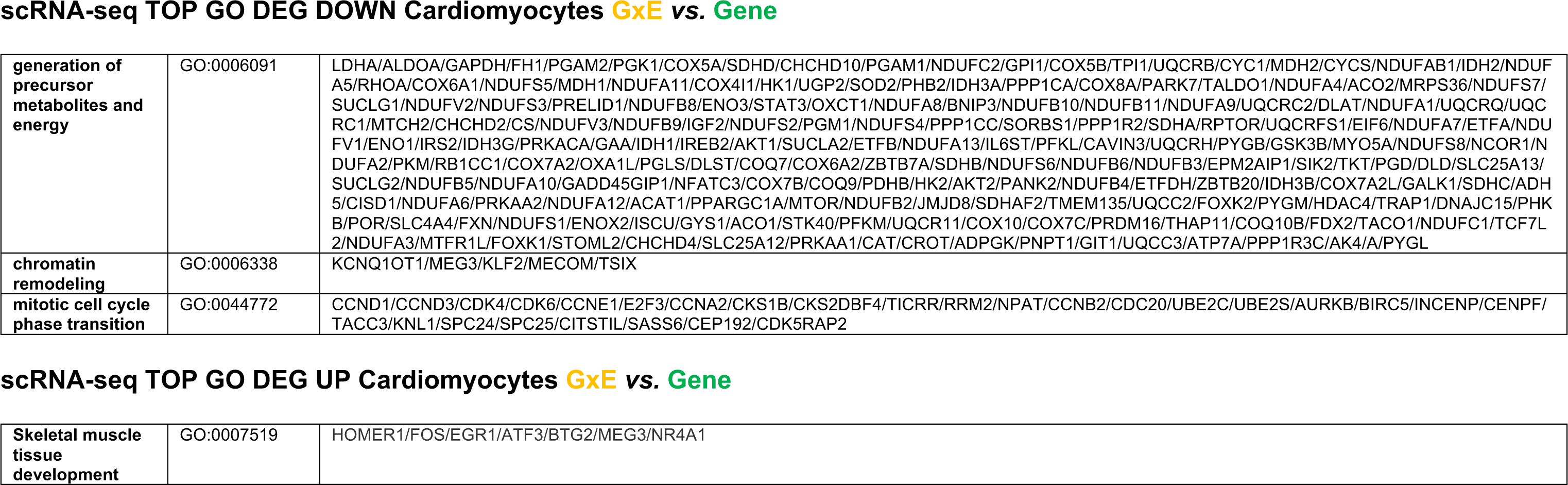
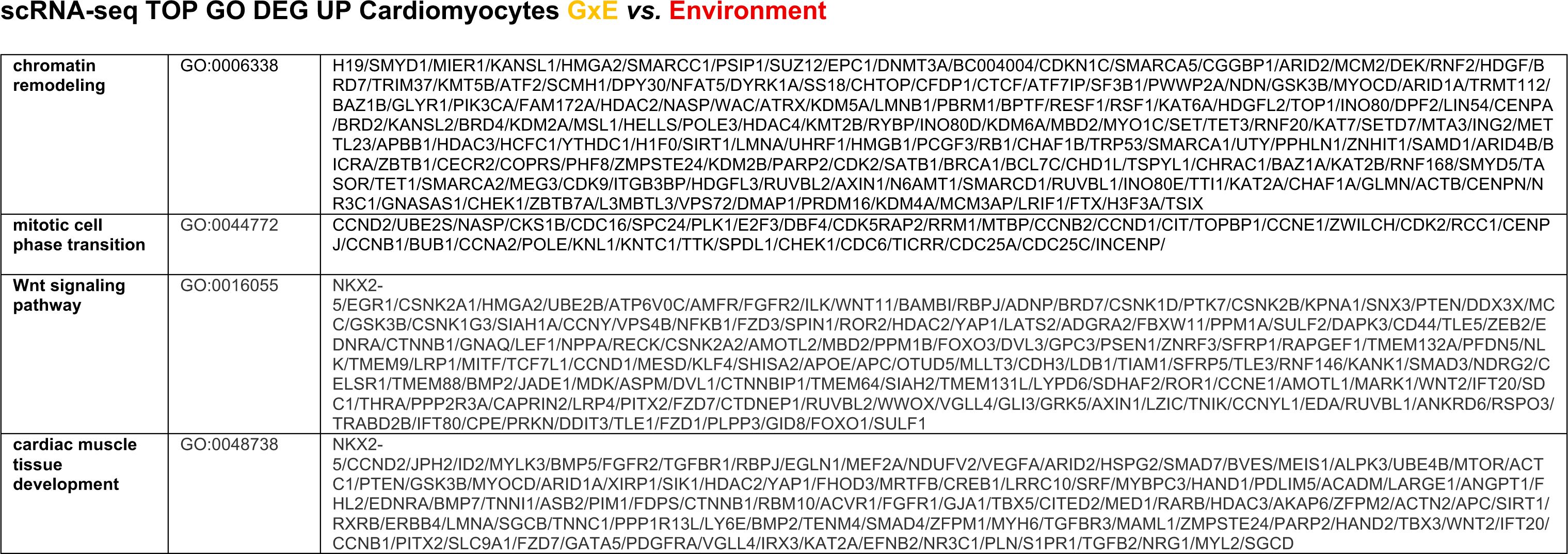
Selected GO terms associated with differentially expressed genes in embryonic cardiomyocytes. Selected GO terms and associated differentially expressed genes identified by scRNA-seq analysis of embryonic cardiomyocytes. Results are presented separately for upregulated and downregulated genes in the G×E versus Control, Environment versus Control, G×E versus Gene, and G×E versus Environment comparisons. GO term descriptions, identifiers, and associated genes are listed. To explore biological processes represented in an extended cardiomyocyte gene set, GO over-representation analysis was performed using both downregulated genes and expressed genes that did not meet the differential expression significance threshold, with the full mouse genome as the background universe.

**Supplementary Table 8.**
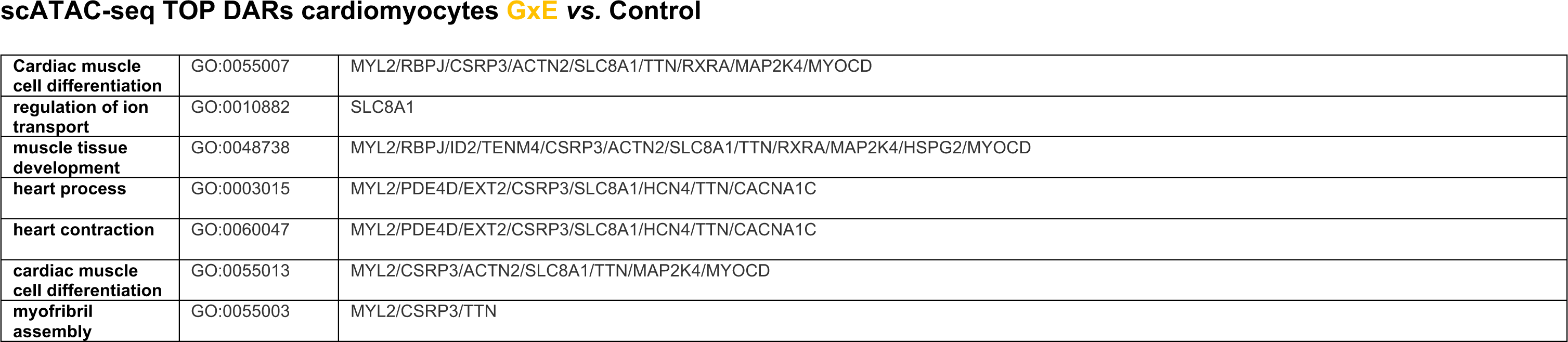
Selected GO terms associated with differentially accessible chromatin regions in embryonic cardiomyocytes. Selected GO terms and associated genes identified by functional enrichment analysis of DARs in embryonic cardiomyocytes in the G×E versus Control comparison. The listed terms relate to cardiac and muscle development, ion transport, cardiac contraction, and myofibril assembly.

**Supplementary Fig. 1.**
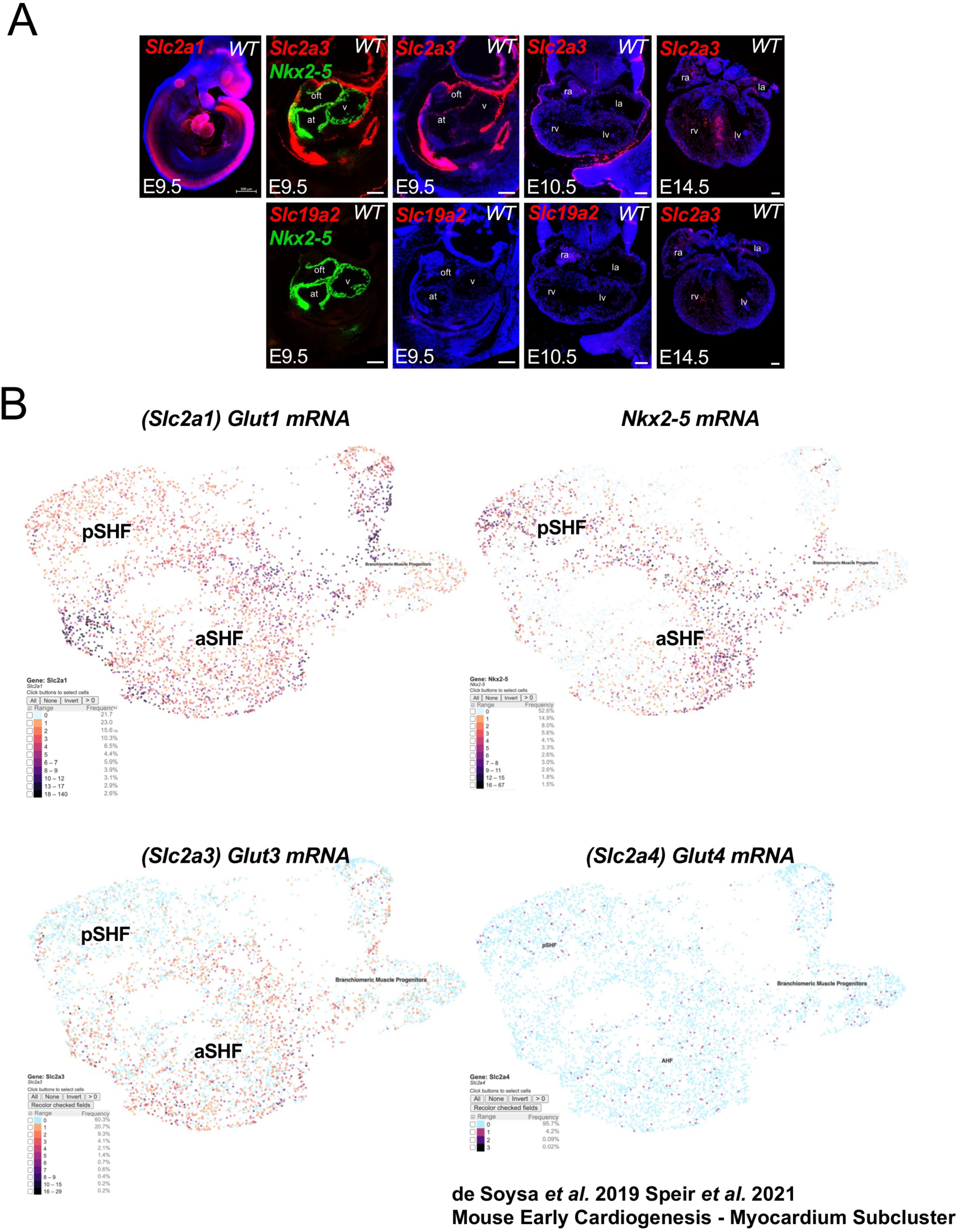
Expression of glucose transporters in early cardiogenesis. (A) Spatial expression of glucose transporter genes during mouse heart development. RNAscope *in situ* hybridization was performed on mouse embryonic heart sections at E9.5, E10.5, and E14.5 to examine the expression of *Slc2a1* (GLUT1), *Slc2a3* (GLUT3), and *Slc19a2*. At E9.5, Slc2a1 (GLUT1) displayed a highly regionalized expression pattern. Co-detection with Nkx2-5 at E9.5 localizes *Slc2a3* expression to the pharyngeal endoderm adjacent to the SHF. *Slc2a3* is highly expressed during early cardiogenesis at E9.5, remains detectable at E10.5, and markedly declines by E14.5. By contrast, *Slc19a2*, which is essential for thiamine-dependent oxidative glucose metabolism, is not expressed in the developing heart throughout cardiogenesis, despite previous reports identifying pathogenic *SLC19A2* variants in patients with congenital heart defects (Conte et al. 2023b). (B) Single-cell myocardium subclusters from published E9.5-E10.5 mouse datasets (de Soysa et al. 2019) are shown to illustrate expression of *Slc2a1* (Glut1), *Slc2a3* (Glut3), and *Slc2a4* (Glut4) across aSHF and pSHF. *Slc2a1* is expressed in SHF progenitors and overlaps with *Nkx2-5*-positive domains. *Slc2a3* is less abundant but detectable mainly in the aSHF. *Slc2a4* is minimal to absent at these stages.

**Supplementary Fig. 2.**
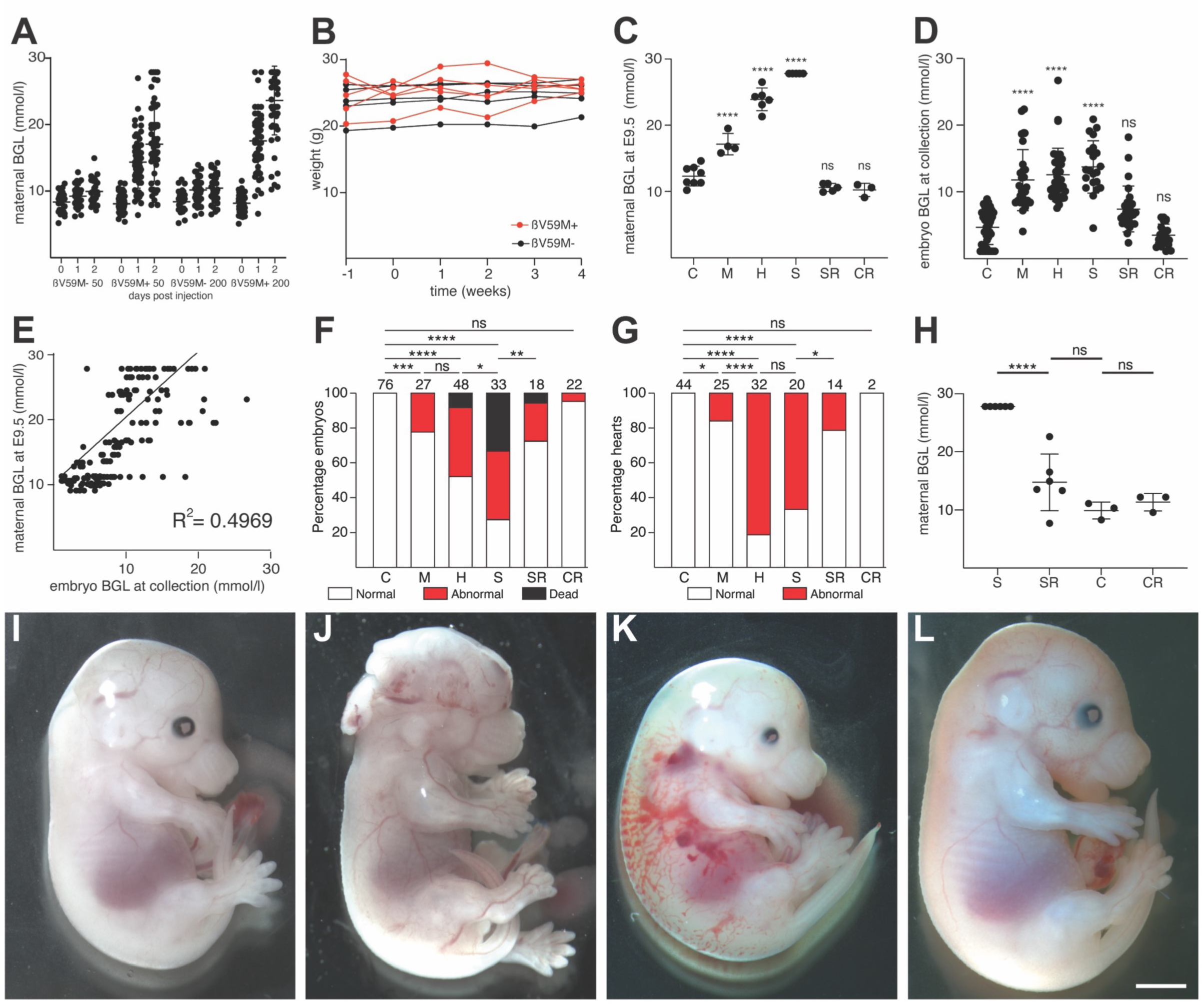
Analysis of the ßV59M mouse model of diabetic embryopathy. (A) Comparison of blood glucose levels between adult female ßV59M mice injected with tamoxifen (50 mg/kg or 200 mg/kg, n=57 each) or corn oil (n=26 and 36, respectively). (B) Comparison of weights of adult female ßV59M mice injected with tamoxifen (n=3) or corn oil (n-=3). (C) Blood glucose levels of pregnant ßV59M mice at E9.5. (D) Comparison of blood glucose levels of E15.5 or E16.5 embryos from control (n=44) or hyperglycemic mothers with moderate (n=27), high (n=35), severe (n=21), severe reversed (n=26) and control reversed (n=22) blood glucose levels. (E) Comparison of maternal and embryonic blood glucose levels. (F,G) Histograms showing significant induction of embryonic abnormality and death (F) and heart defects (G) by increasing levels of maternal hyperglycemia and significant rescue by reversal of severe maternal hyperglycemia prior to embryo implantation. (H) Reversal of severe maternal hyperglycemia by glibenclamide treatment prior to embryo implantation. (I-L) Representative images of a control E15.5 embryo (I) and E16.5 embryos from mothers with high (J), severe (K) and severe reversed (L) hyperglycemia. Statistical significance for (C,D) was tested by Kruskal Wallis ANOVA with Dunn’s multiple comparison post hoc test comparing to control; for (F,G) by two-tailed Fisher’s exact test; (E) by simple linear regression; (H) by AOVA with Tukey’s post hoc test. *, P<0.95; **, P<0.01; ****, P <0.0001; ns, not significant. C, control; M, moderate; H, high; S, severe; SR, severe reversed; CR, control reversed. Scale bar = 2 mm (I-L).

**Supplementary Fig. 3.**
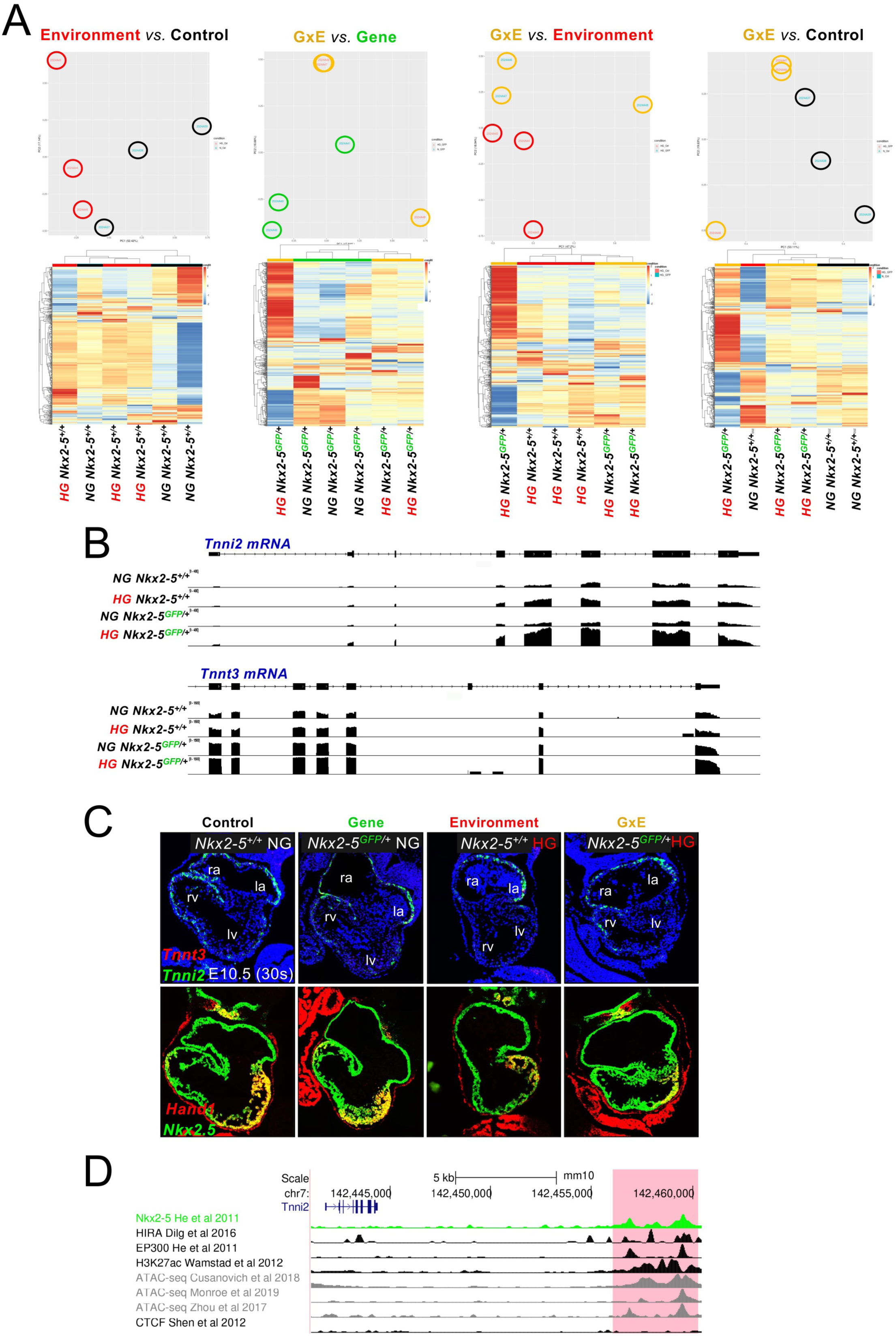
Bulk RNA-seq DEG analysis and early spatial validation of contractile markers and *Hand1* expression. (A) Principal component analysis (PCA) of E15.5 bulk RNA-seq libraries (n = 3 per group) and heat maps of differentially expressed genes (DEG) for the four comparisons (Environment *vs* Control; G×E *vs* Gene; G×E *vs* Environment; G×E *vs* Control). (B) Gene-level expression summaries (normalized counts) for *Tnni2* and *Tnnt3* across the four groups, highlighting upregulation in the G×E condition in the bulk dataset in the browser view. (C) RNAScope on E10.5 (∼30 somite) heart sections showing spatial expression of *Tnni2* and *Tnnt3*. In NG *Nkx2-5^GFP/+^* embryonic hearts, *Tnnt1* expression was undetectable, whereas *Tnni2* was detected in the developing atrial and ventricular myocardium. At this stage, the upregulation observed in bulk RNA-seq data at E15.5 was not yet apparent. Serial sections show co-detection of *Hand1* and *Nkx2-5*. *Hand1*, a marker of the left ventricle, was reduced in the Gene and G×E groups relative to Control. (D) NKX2-5 and HIRA co-occupy at a shared enhancer regulating *Tnni2-Tnnt3* expression (Dilg et al. 2016; Dupays et al. 2015)

**Supplementary Fig. 4.**
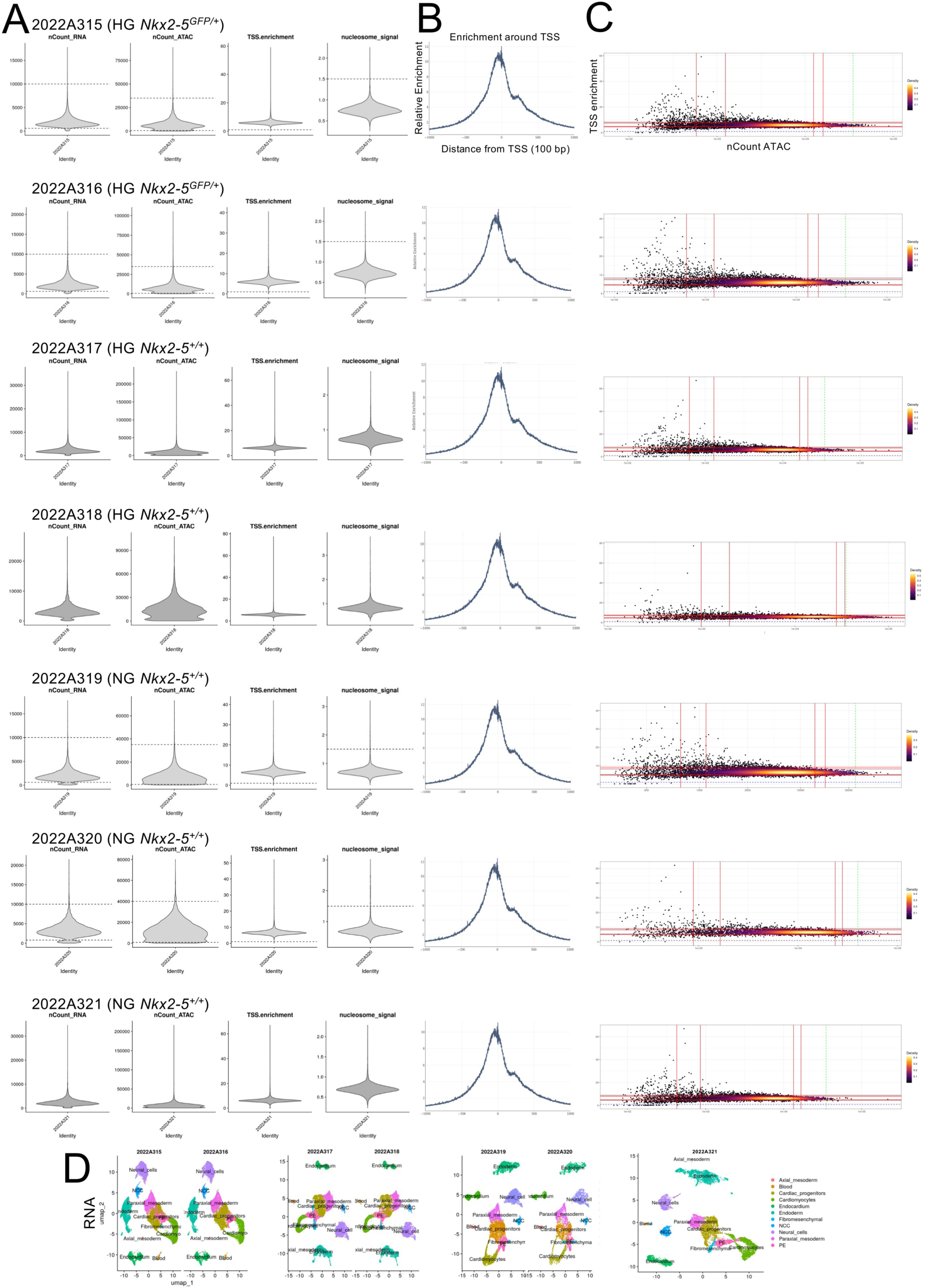
Quality control of single-nucleus multiome datasets. (A) Violin plots showing the distribution of RNA counts, ATAC counts, TSS encrichment scores, and nucleosome signals across samples. (B) Average TSS enrichment plots of snATAC-seq data demonstrating a mean enrichment score of 10-12. (C) Scatte plot of TSS enrichment scores versus ATAC counts for snATAC-seq data, showing the relationship between sequencing depth and signal quality. Red lines represent 5, 10, 90 and 95 % quantiles. (D) snRNA-seq corresponding to the integrated analyses represented in figure 4C. UMAP plots showing individual replicates for RNA. Nuclei and their clustering were consistent across replicates.

**Supplementary Fig. 5.**
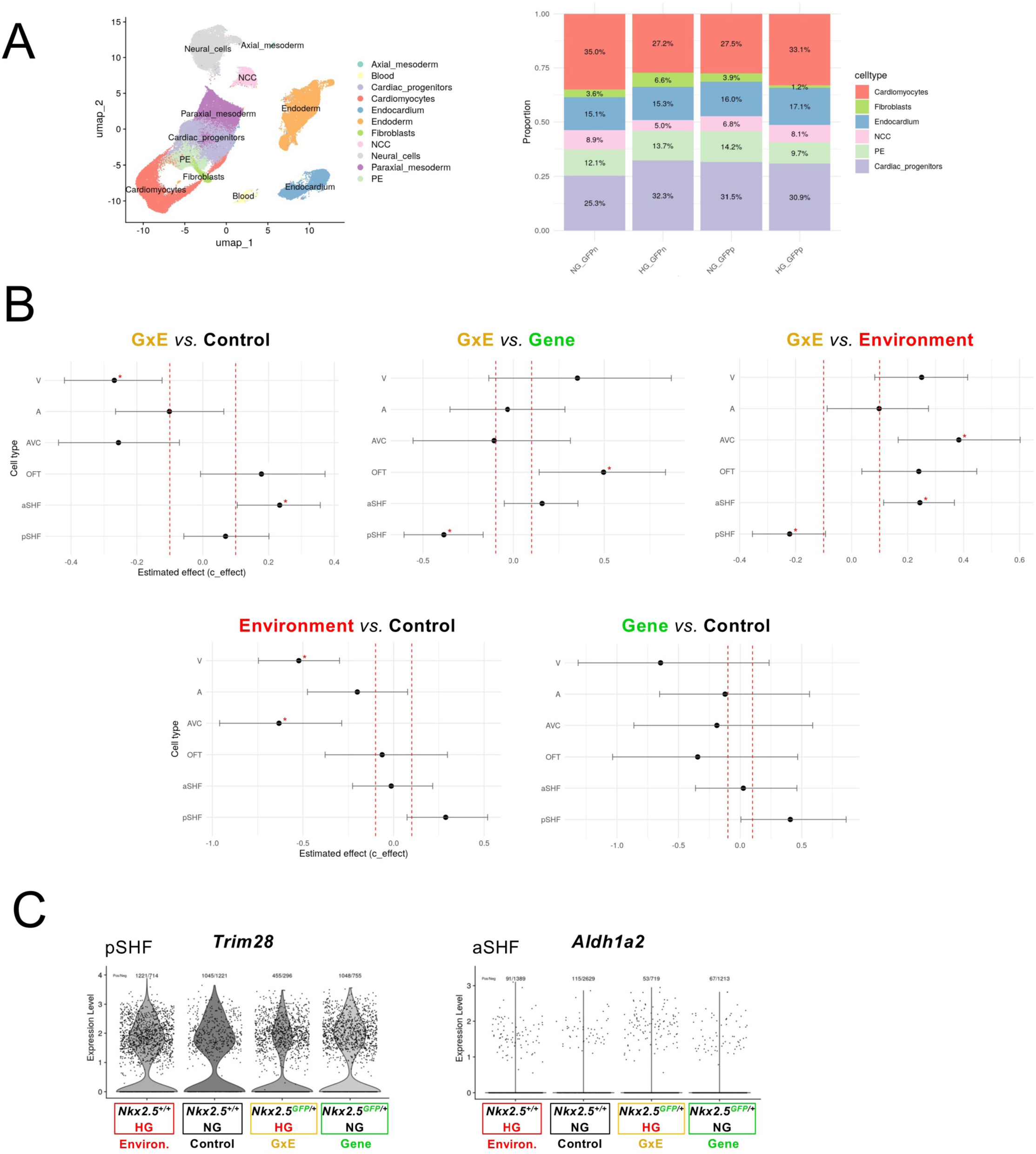
Cell-type composition and differential abundance at E9.5. (A) UMAP of E9.5 snMultiome nuclei from the four groups showing major lineages: cardiomyocytes, cardiac progenitors, endocardium, fibroblasts, neural crest cells (NCC), pro/epicardium (PE), paraxial mesoderm, blood, and endoderm. Cellular distribution within each experimental group is represented by bar plots showing cell-type proportions normalized to total cell number per group. (B) Differential abundance analysis within the cardiomyocytes and cardiac progenitors subclusters was performed on scRNA-seq data. Red asterix indicate significant changes in cellular proportions across the different comparisons, the zone between the two vertical dashed lines marks zero effect. (C) Violin plots displaying single-nucleus RNA expression of *Aldh1a2* in aSHF and *Trim28* in pSHF across the four experimental groups. *Trim28* is involved in DNA methylation (Panzeri et al., 2025). (V) Ventricle, (A) atria, (AVC) atrioventricular canal, (OFT) outflow tract, (aSHF) anterior second heart field, and (pSHF) posterior second heart field state.

**Supplementary Fig. 6.**
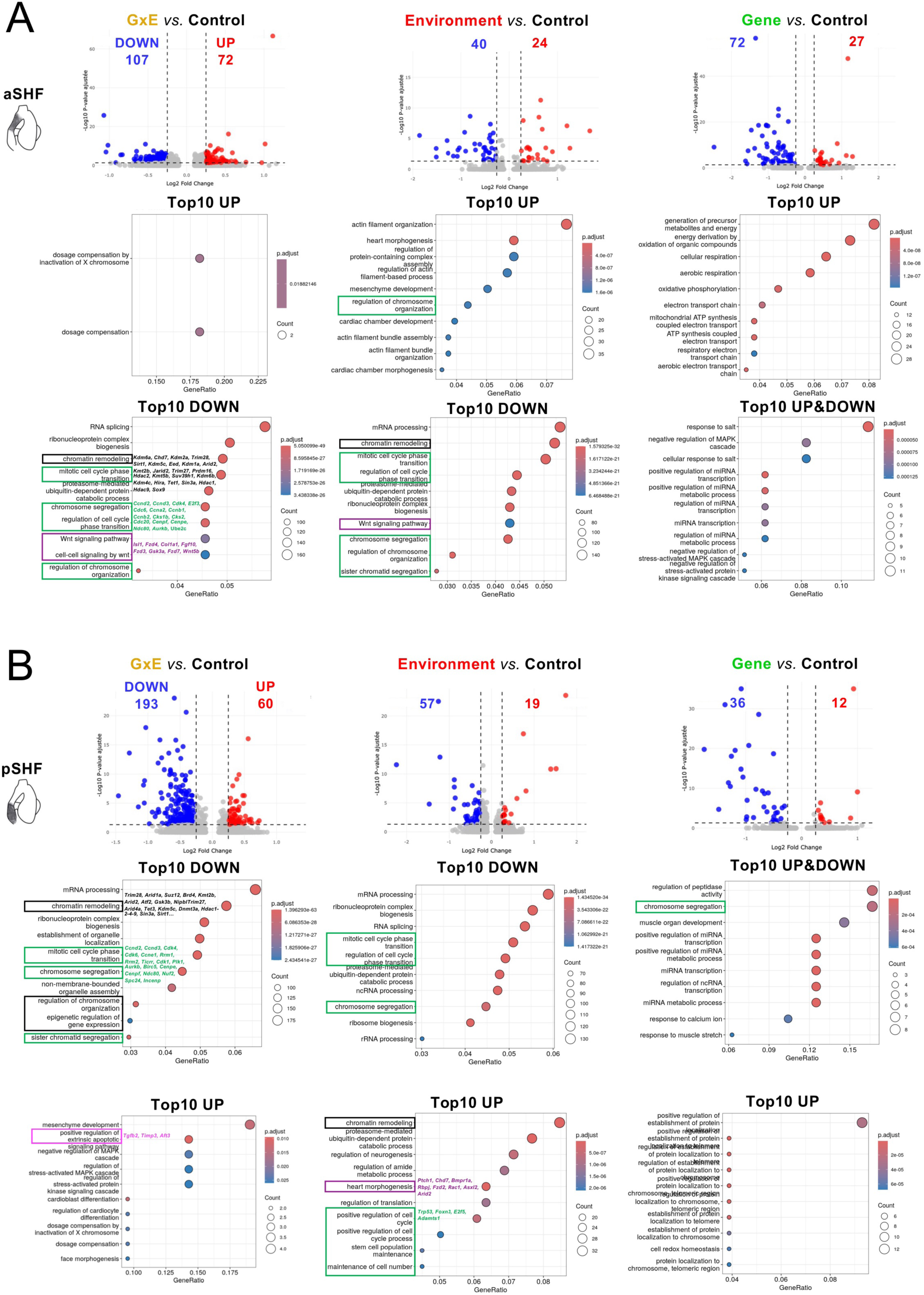
DEG and Top10 GO terms in SHF subclusters. Differential expression analysis was performed in the aSHF and pSHF subclusters by comparing the G×E, Environment, and Gene groups with the Control group. Volcano plots show significantly downregulated (blue) and upregulated (red) genes per contrast (vertical dashed lines = log2FC threshold; numbers indicate DEG counts). GO dot plots summarize the top enriched biological processes for down regulated gene sets in aSHF and pSHF across these 3 comparisons. Recurrent themes include chromatin remodeling, mitotic cell-cycle transition, Wnt signaling, along with apoptosis/ROS programs.

**Supplementary Fig. 7.**
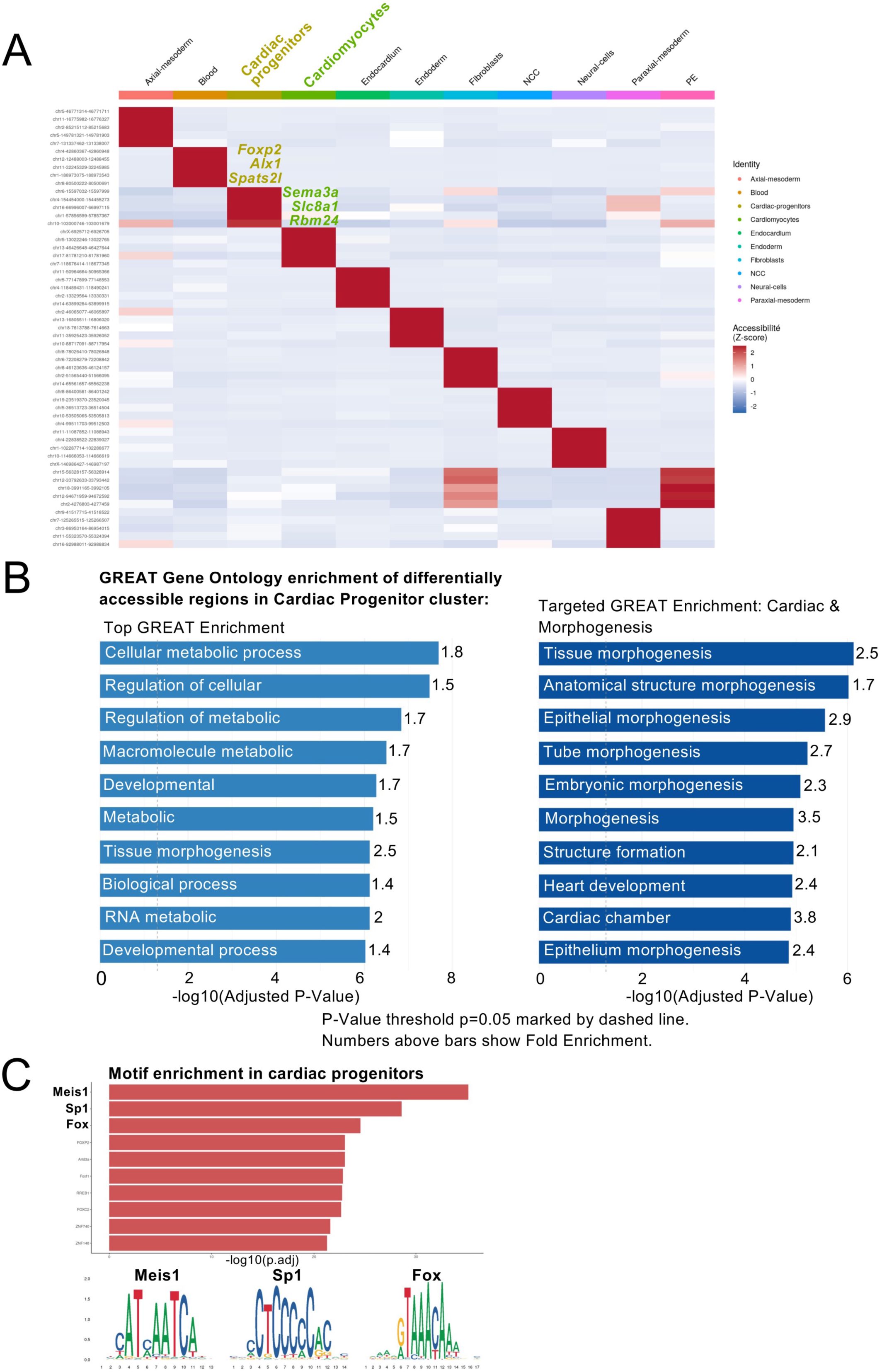
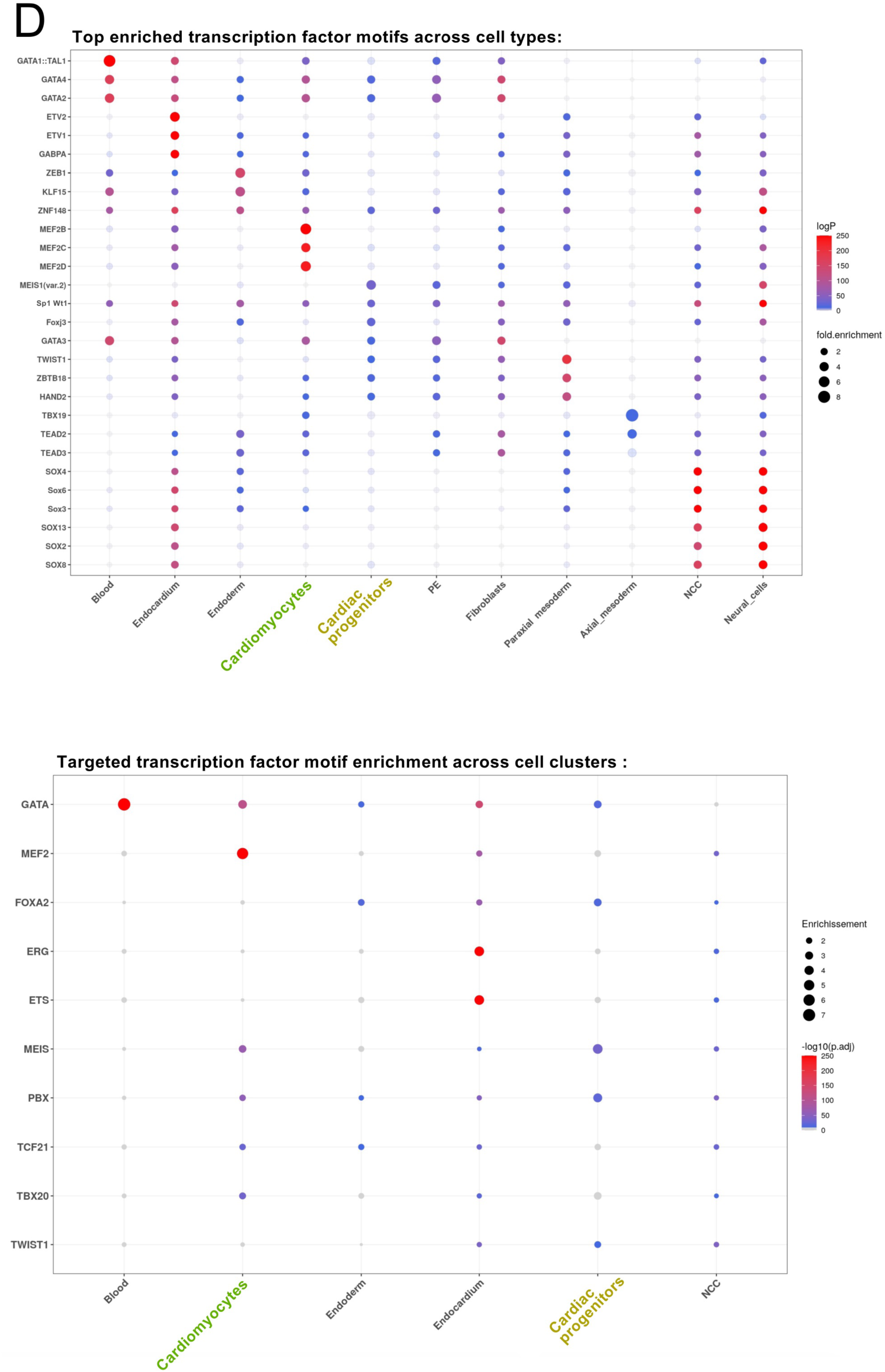
Chromatin accessibility profiles validate the identity of cardiac progenitors in the scATAC-seq dataset. (A) Chromatin accessibility heatmap of the top 5 marker peaks per cell type. Rows represent genomic coordinates of marker peaks, and columns represent cell types. Chromatin accessibility is shown as Z-scores, ranging from low (blue) to high (red). (B) GREAT analysis of DARs identified in cardiac progenitors reveals significant enrichment for developmental and morphogenetic processes, including heart development, cardiac chamber morphogenesis, tissue morphogenesis, and embryonic morphogenesis. (C) Motif enrichment analysis of progenitor-specific accessible regions identifies significant enrichment for binding motifs of cardiac developmental transcription factors, including Meis1, Sp1, and Fox family members. Together, these results confirm that the scATAC-seq dataset captures regulatory landscapes characteristic of embryonic cardiac progenitors. (D) Transcription factor motif enrichment across embryonic cardiac cell populations. Dot plot showing the enrichment of transcription factor binding motifs across major embryonic cardiac cell types identified by scATAC-seq. The upper panel displays the most enriched motifs in each cell population, whereas the lower panel highlights selected cardiac developmental transcription factors. Cardiac progenitors exhibit enrichment for regulatory motifs including Meis, GATA, HAND, T-box, consistent with their developmental identity and supporting the biological relevance of the chromatin accessibility profiles. Dot size indicates fold enrichment and color represents statistical significance (-log10 adjusted *P* value).

**Supplementary Fig. 8.**
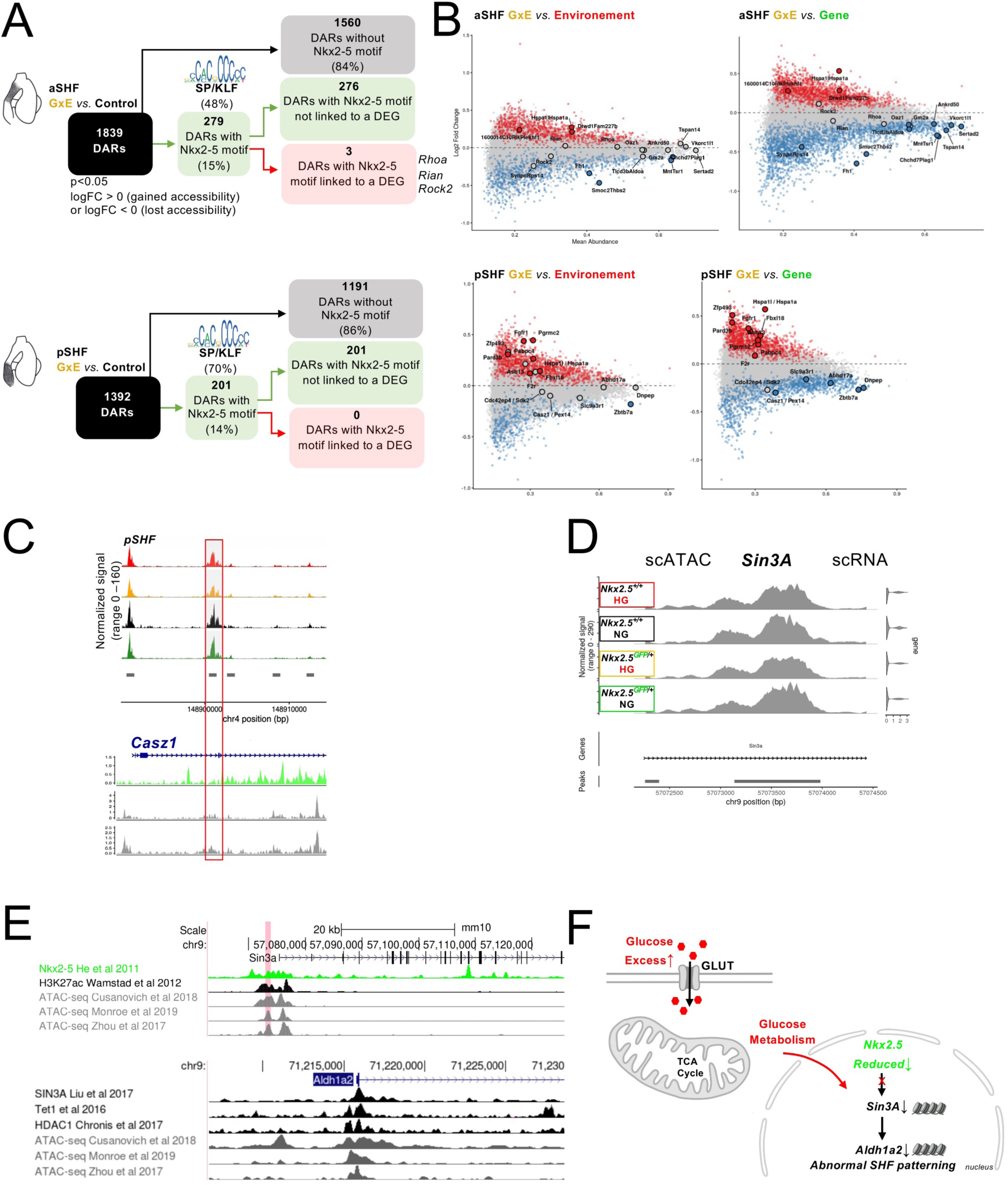
Integrative multiomic analysis identifies candidate regulators linking maternal hyperglycemia to altered developmental signaling. (A) Schematic overview of the integrative analysis used to identify candidate direct Nkx2-5 regulatory targets. DARs identified in aSHF and pSHF progenitors were sequentially filtered to retain regions containing an NKX2-5 consensus binding motif, before integration with differentially expressed genes to generate a high-confidence list of candidate regulatory elements. The schematic also summarizes the transcription factor-binding motifs enriched within candidate regulatory regions in each progenitor population. (B) MA plots showing differential chromatin accessibility in G×E versus Environment and G×E versus Gene comparisons in aSHF and pSHF progenitors. Regions with increased accessibility are shown in red and regions with decreased accessibility in blue (adjusted *P* < 0.05). (C) Genome browser tracks showing representative candidate regulatory regions associated with *Casz*. Differentially accessible regions (DARs) identified in GxE progenitors are displayed together with previously published NKX2-5, HDAC1/2 ChIP-seq tracks. Red boxes highlight the DARs shown in the analysis. (D) Integrated multiomic analysis identifies *Sin3a* as a candidate regulatory locus exhibiting reduced chromatin accessibility in GxE cardiomyocytes together with decreased transcript abundance. snATAC-seq tracks depicting chromatin accessibility at the *Sin3a* locus between groups (differential peak testing using MACS v3.0.2, min.pct = 0.1, adjusted P < 0.05)*. Sin3a* was identified *via* an exploratory analysis filtering for peaks showing at least a ±5% shift in cell frequency (defined as the proportion of cells with at least one fragment overlapping the peak) between G×E and Control conditions. (E) Publicly available Nkx2-5 ChIP-seq and ATAC-seq datasets support occupancy of the *Sin3a* locus by Nkx2-5, whereas published SIN3A ChIP-seq data demonstrate binding at the *Aldh1a2* regulatory region, supporting a potential *Nkx2-5–SIN3A–Aldh1a2* regulatory axis. (F) Working model illustrating the proposed mechanism whereby maternal hyperglycemia, together with reduced Nkx2-5 dosage, decreases *Sin3a* expression, leading to reduced *Aldh1a2* expression and impaired second heart field (SHF) patterning.

**Supplementary Fig. 9.**
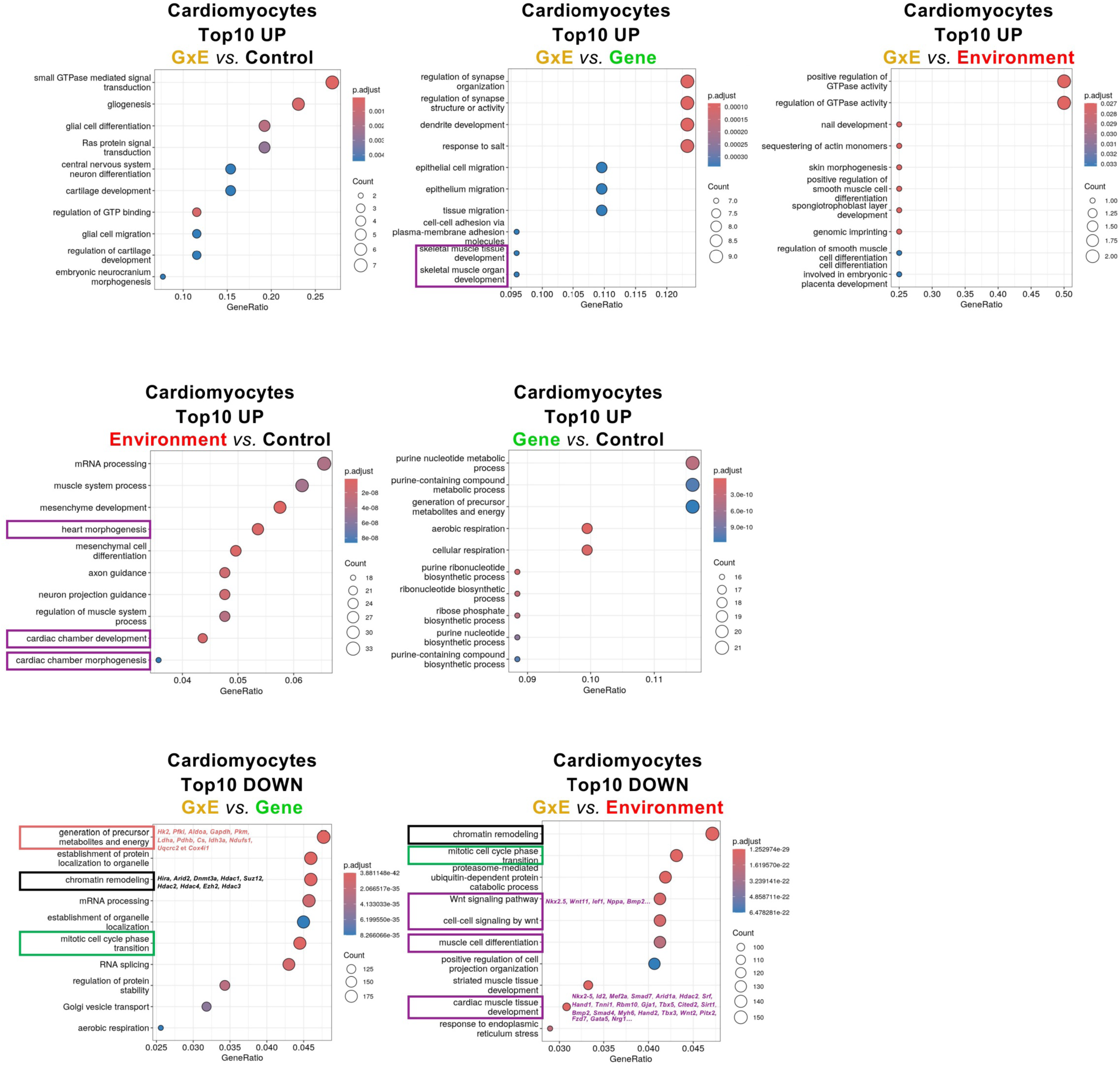
Top 10 GO terms related to disregulated genes in cardiomyocyte cluster. Gene Ontology (GO) enrichment analysis was performed using the enrichGO function from the clusterProfiler R package. The top 10 GO terms enriched among upregulated genes in cardiomyocytes are shown for the G×E *vs.* Control, G×E *vs.* Gene, G×E *vs.* Environment, Environment *vs.* Control, and Gene *vs.* Control comparisons. The top 10 GO terms enriched among downregulated genes are shown for the G×E *vs.* Gene and G×E *vs.* Environment comparisons.

### Supplementary Spreadsheet 1. Marker genes defining embryonic cardiac cell populations and subclusters

Significant marker genes identified across the global cell clusters and fine cell-type annotations, used to characterize embryonic cardiac cell populations and their subclusters.

### Supplementary Spreadsheet 2. Differential gene expression in anterior second heart field progenitors

Lists of differentially expressed genes in aSHF progenitors for the experimental comparisons analysed.

### Supplementary Spreadsheet 3. Differential gene expression in posterior second heart field progenitors

Lists of differentially expressed genes in pSHF progenitors for the experimental comparisons analysed.

### Supplementary Spreadsheet 4. Differential chromatin accessibility in anterior second heart field progenitors

DARs identified in aSHF progenitors for the experimental comparisons analysed.

### Supplementary Spreadsheet 5. Differential chromatin accessibility in posterior second heart field progenitors

DARs identified in pSHF progenitors for the experimental comparisons analysed.

### Supplementary Spreadsheet 6. Differential gene expression in embryonic cardiomyocytes

Lists of differentially expressed genes in embryonic cardiomyocytes for the experimental comparisons analysed.

### Supplementary Spreadsheet 7. Differential chromatin accessibility in embryonic cardiomyocytes

DARs identified in embryonic cardiomyocytes for the experimental comparisons analysed.

## Acknowledgements

We thank Philippe Guichard, zootechnician, and Clotilde Ferreira, for their technical assistance, the animal facility, in particular Dr Erica Lopez (VMD), Perrine André, Thierry Cheyrol, and Clémence Grillot. RNA sequencing and single-cell multiome library preparation were conducted by the the Genomics and Bioinformatics facility GBiM platform; we are grateful to GBiM, namely Dr Valérie Delague, Christel Castro, Nathalie Trévisiol and Coralie Provost, from the U 1251/Marseille Medical Genetics laboratory, for performing quality control and preliminary analyses of the RNA-Seq and scMultiome data. ScMultome sequencing was performed by IntegraGen (Evry). HREM was conducted by the SFR Necker platform (Paris); we thank Pr Lucile Houyel and Pr Bill Chaudry for their help with HREM data analysis. We thank Dr Stéphane Zaffran and Pr Richard Harvey for the *Nkx2-5^GFP/+^* line (https://www.jax.org/strain/024662). This work was supported by INSERM, the French National Research Agency (ANR-JCJC ANR-20-CE13-0018 REPACE to SS), the European Research Area Network on Cardiovascular Diseases (ERA-Net ANR-19-ECVD-0002 DEFIANCE to SS, BT, and AKK), British Heart Foundation Senior Basic Science Research Fellowships FS/17/55/33100 and FS/SBSRF/22/31022 (DBS); Oxford BHF Centre for Research Excellence core infrastructure grants (DBS and JIKS, RE/18/3/34214); the Benefaction for Medicine and the Wellcome Trust Institutional Strategic Support Fund (DBS and NV); and Oxford Novo Nordisk postdoctoral fellowships (NV, JJM). SI was supported by a doctoral fellowship from the Fondation pour la Recherche Médicale (FDT-fin de thèse 202404018474).

## Funding Information

This paper was supported by the following grants:

Fondation de France 00171134/WB-2025-58883, Agence Nationale de la Recherche ANR-20-CE13-0018 to Sonia Stefanovic

Agence Nationale de la Recherche ANR-19-ECVD-0002 to Anne-Karin Kahlert, Sonia Stefanovic, Bernard Thienpont.

Fondation pour la Recherche Médicale FDT-fin de thèse 202404018474 to Stéphanie Ibrahim.

## Author contributions

Conceptualization: S.S, D.B.S, A.K.K, B.T,

Methodology: J.M, S.S, J.J.M,

Software: J.J.M, J.M, C.H

Validation: S.S

Formal analysis: S.S, A.K.K, D.B.S, D.S, J.M, N.V

Investigation: S.I, S.S, D.B.S, D.C, D.S, G.C, J.J.M, N.V, S.A,

Resources: F.M.A.

Data Curation: S.S

Writing - Original Draft: S.S

Writing - Review & Editing: S.S, D.B.S

Visualization: S.S, D.B.S, N.V

Supervision: S.S, D.B.S

Project administration: S.S, D.B.S

Funding acquisition: S.S, D.B.S, A.K.K, B.T

